# A Novel Selective ERK1/2 Inhibitor, Laxiflorin B, Targets EGFR Mutation Subtypes in Non-small-cell Lung Cancer

**DOI:** 10.1101/2022.06.27.497627

**Authors:** Chengyao Chiang, Min Zhang, Junrong Huang, Juan Zeng, Chunlan Chen, Dongmei Pan, Heng Yang, Min Yang, Qiangqiang Han, Wang Zou, Tian Xiao, Yongdong Zou, Feng Yin, Zigang Li, Lizhi Zhu, Duo Zheng

**Author notes:** Authors with equal contributions (Chengyao Chiang, Min Zhang, Junrong Huang, Juan Zeng).

## Abstract

Extracellular regulated protein kinases 1/2 (ERK1/2) are key members of multiple signaling pathways including the ErbB axis. ERK1/2 ectopic activation is responsible for various types of cancer, especially drug resistance to inhibitors of RTK, RAF and MEK, but specific ERK1/2 inhibitors are scarce. In this study, we identified a potential novel ERK covalent inhibitor, Laxiflorin B, a herbal compound with anticancer activity. However, Laxiflorin B is present at low levels in herbs; therefore, we adopted a semi-synthetic process for the efficient production of Laxiflorin B to improve the yield. Laxiflorin B induced mitochondria-mediated apoptosis via BAD activation in non-small-cell lung cancer (NSCLC) cells, especially in EGFR mutant subtypes. Transcriptomic analysis suggested that Laxiflorin B inhibits amphiregulin (AREG) and epiregulin (EREG) expression through ERK inhibition, and suppressed the activation of their receptors, ErbBs, via a positive feedback loop. More importantly, mass spectrometry combined with computer simulation analysis revealed that Laxiflorin B binds covalently to Cys-183 in the ATP-binding pocket of ERK1 through D-ring, and Cys-178 of ERK1 though non-inhibitory binding of A-ring, respectively. Laxiflorin B also exhibited strong tumor suppressive effects with low toxicity in a NSCLC tumor xenograft model in nude mice, and AREG and EREG were identified as biomarkers of Laxiflorin B efficacy. Finally, Laxiflorin B-4, a C-6 modification of Laxiflorin B, exhibited higher affinity for ERK1/2 and stronger tumor suppression. These findings provide a new approach to tumor inhibition using natural anticancer compounds.

## Introduction

Extracellular regulated protein kinases (ERKs), which belong to the mitogen-activated protein kinase (MAPK) family, play pivotal roles in regulating multiple effectors in cell proliferation and survival pathways under both physiological and pathological conditions ^1^. In cancer development and progression, abnormal activation of ERK1/2 by upstream signals caused by events such as mutation or amplification in receptor tyrosine kinases (RTKs), RAS, proto-oncogene serine/threonine-protein kinases (RAFs) and mitogen-activated protein kinases (MAPK)/ERK kinases (MEKs) ^1^, result in malignancy and poor patient prognosis ^2,3^.

Due to the prevalence of activation of the RAS-RAF-MEK-ERK axis in various types of malignancies, kinase inhibitors have been a common strategy for clinical treatment. Three types of RAS activation mutation account for 16% of human cancers ^4^. These mutations are generally regarded as intractable GTPases due to structural and affinity characteristics ^5,6^, although ARS-1602 ^7^ and MRTX849 ^8^ have been shown to be effective treatments in cases with the KRAS12C mutation, which accounts for a small fraction of human malignancies. First-generation RAF inhibitors for the major mutation V600E, such as Vemurafenib, Dabrafenib and Encorafenib, were reported to be associated with the rapid development of drug resistance and paradoxical effects due to upregulation of RAS expression and alternative splicing of BRAF leading to a reduction in the affinity of the inhibitors^9,10^. The second-generation RAF inhibitors, which were designed to intercept RAF-dimer formation, are currently undergoing clinical trials^11^. Multiple mutation sites and compensatory pathways have also been described for MEK, a crucial mitogen-activated kinase activated by RAF, resulting in increased resistance to the MEK inhibitor, AZD6244, and the RAF inhibitor, PLX4720^12^, as well as poor prognosis in human malignancies^3^. Several studies have shown that prolonged administration of inhibitors of RTK, RAF and MEK result in drug resistance and re-activation of ERK1/2 in cancer cells ^10^. Therefore, an effective strategy for targeting ERK is urgently required to improve the prognosis of numerous cancer patients. Hence, in addition to providing a greater understanding of the MAPK pathway, the identification and development of ERK1/2 inhibitors may overcome the challenges to the use of RAF and MEK inhibitors for the treatment of a variety of cancers.

Compared with the upstream RTKs, RAS, RAF and MEK, ERK1/2 has a more conservative amino acid sequence and structure, with very few mutations in cancer cell genomes^13^. By phosphorylation, ERK1/2 regulates widely diverse substrates responsible for evolutionarily conserved cellular processes, such proliferation, differentiation, growth, metabolism, migration and survival. Thus, the ERK1/2-associated signaling cascade is strictly controlled by a series of negative regulators, such as, dual-specificity phosphatases (DUSPs) ^1^. Among the current strategies for ERK1/2 inhibition in malignancies, ATP analogues, such as pyridine-pyrrole, indazole-pyrrolidine and pyrazole amino-pyrimidine derivatives, have been demonstrated to inhibit ERK1/2 effectively by blocking the ATP-binding pocket ^14^. In previous studies, Ulixertinib was shown to inhibit ERK1/2 with high specificity and delayed drug resistance compared with RAF or MEK inhibitors^15^. Moreover, Ulixertinib was found to overcome the drug resistance of RAF inhibition in a patient-derived xenograft model and a phase I clinical trial^16^. Ravoxertinib, FR180204 and LY3214996 mediated effective inhibition of ERK1/2 ^17–19^, with synergistic effects achieved by using a RAF inhibitor compared with monotherapy in RAF- or KRAS-mutant malignancies in phase I clinical trials ^20,21^. Robust efficacy of SCH7779684, MK-8353 and LY3214996 were reported for KRAS- or RAF-mutant cancer cells in a xenograft model ^19,22,23^. On the other hand, effective ERK1/2 inhibitors have rarely been reported in clinical trials ^14^.

*Isodon eriocalyx* (*I. eriocalyx var.* laxiflora) is a widely used herbal medicine in China. Research has implicated *ent*-kaurene diterpenes isolated from *Isodon* plants as novel anticancer drugs with broad potential ^24^ . Laxiflorin B, was shown to inhibit cancer cell viability ^25^. However, comprehensive studies of this compound are hindered by low (0.00061%) abundance^26^.

Here, we developed a new strategy to improve to yield of Laxiflorin B through semi-synthetic production from its analogue, Eriocalyxin B. We demonstrated that Laxiflorin B exerts strong anti-cancer activity by inhibiting the proliferation and promoting apoptosis in non-small-cellular lung cancer (NSCLC) cells expressing a constitutively active EGFR. Moreover, the ErbB pathway was primarily influenced by Laxiflorin B application and ERK1/2 was identified as a novel potential target of Laxiflorin B. The cystine (Cys) residue at position 183 and 178 of ERK1 was identified as the covalent binding sites for Laxiflorin B.

Subsequently, we found that BAD, an initiator of mitochondria-mediated apoptosis regulated by the ERK-RSK axis, was activated by Laxiflorin B-induced ERK inhibition, and this effect was rescued by BAD elimination. Simultaneously, the ERK-driven secretion of growth factors, such as AREG and EREG, was largely abolished by Laxiflorin B-targeted ERK suppression, resulting in a positive feedback loop of ErbB signaling induced by autocrine AREG and EREG downregulation. Our *in vivo* study revealed that Laxiflorin B exerted strong antitumor effects with low toxicity, implying a wide range of possibilities for modification and therapeutic applications.

## Results

### Semi-synthetic production of Laxiflorin B from its natural analogue, Eriocalyxin B

Eriocalyxin B and Laxiflorin B were isolated from dry leaves of *Isodon eriocalyx. laxiflora* with 0.084% and 0.00067% yields^26^, respectively. To obtain large amounts of Laxiflorin B, we designed semi-synthetic route began with the oxidative cleavage of the C6–C7 carbon-carbon bond of Eriocalyxin B. By treating the reflux solution of Eriocalyxin B in DCM with Dess–Martin periodinane, the vicinal diols at C6-C7 were oxidized to aldehyde. The aldehyde functional group of the oxidation product from the final step was selectively reduced with NaBH_4_ under acidic conditions. Column chromatography showed that the total yield of Laxiflorin B was 70% (Fig. 1).

**Fig. 1.**
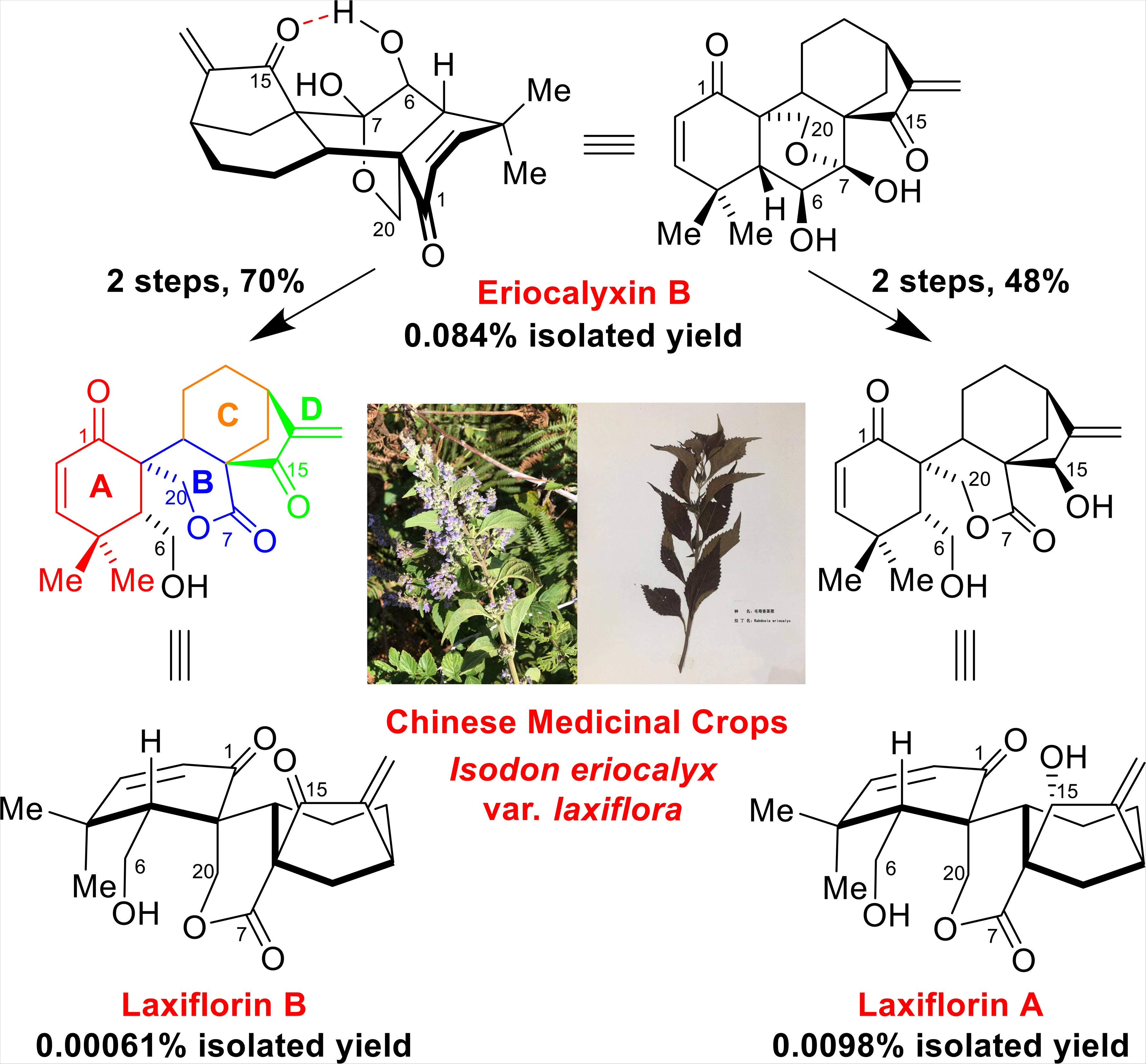
Semi-synthetic production of Laxiflorin B from Eriocalyxin B. Laxiflorin B was prepared at scale from Eriocalyxin B through oxidative cleavage of the C6–C7 carbon-carbon bond with Dess–Martin periodinane, followed by selective reduction of aldehyde with NaBH_4_ under acidic condition.

### Laxiflorin B inhibits the progression and promotes apoptosis of NSCLC cells

Laxiflorin B has been shown to inhibit cancer cell viability ^25^. We first examined the anticancer efficacy of Laxiflorin B in NSCLC cell lines. The growth of PC9, HCC827, H1650 (Fig. 2a), A549 and H1975 (Fig. S1a and Table 1) cells was inhibited by Laxiflorin B in a dose-dependent manner. This dose-dependent inhibition of cell viability was confirmed in 2-D clonogenic assays using PC9, HCC827, H1650 (Fig. 2b), A549 and H1975 (Fig. S1b) cells. Laxiflorin B also inhibited the anchorage-independent growth of PC9 cells (Fig. 2c). These results indicated that Laxiflorin B is a natural compound with strong anticancer effects at low concentration.

**Fig. 2.**
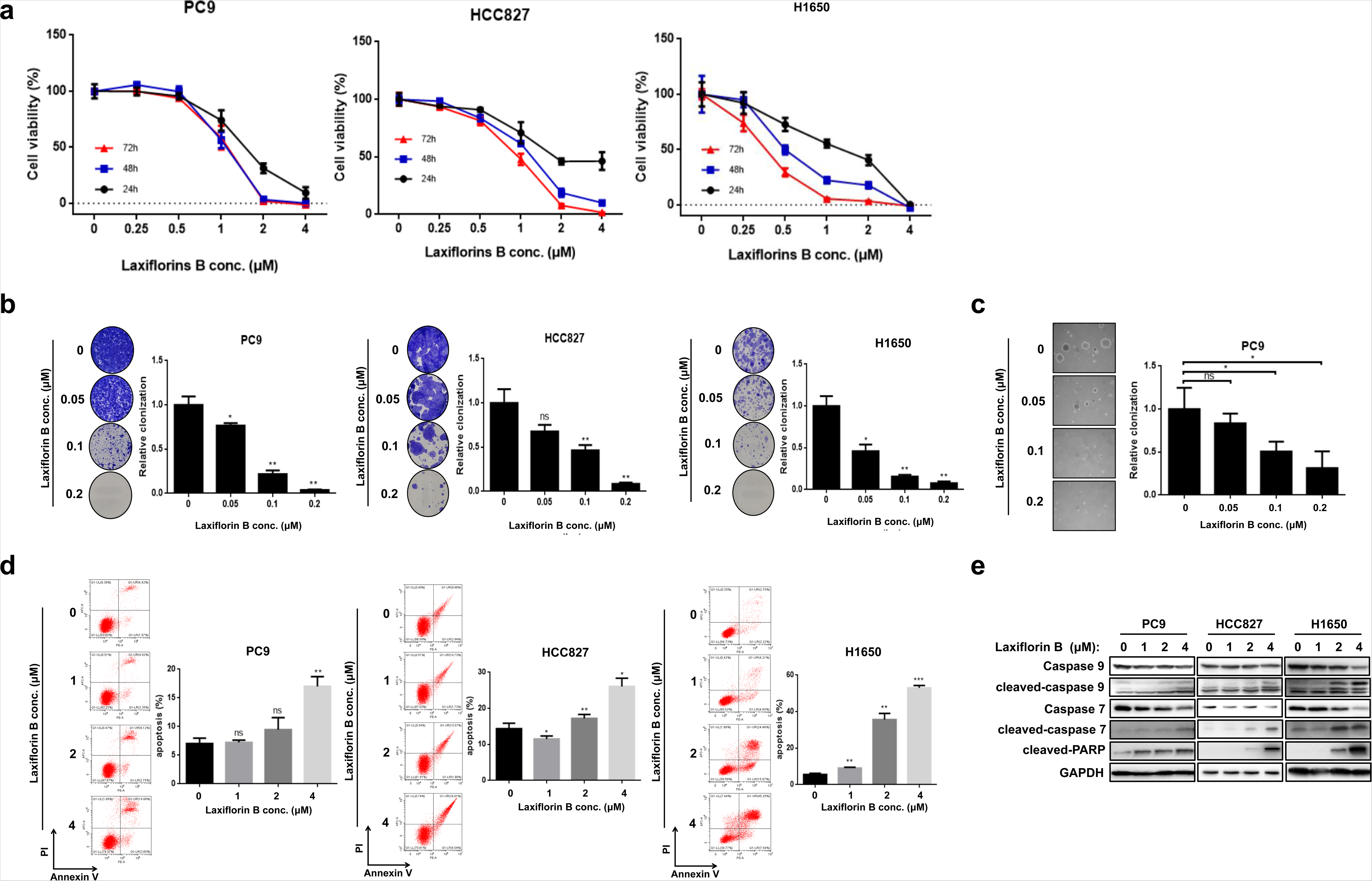
Inhibitory effects of Laxiflorin B in non-small cellular lung cancer cells. **a** Viability of PC9, HCC827 and H1650 cells after Laxiflorin B treatment. **b** Clonogenic assays of PC9, HCC827 and H1650 cell growth after Laxiflorin B treatment for 2–3 weeks. **c** Anchorage-independent growth of PC9 cells after Laxiflorin B treatment for 31 days. **d** Flow cytometric analysis of apoptosis of PC9, HCC827 and H1650 cells following Laxiflorin B treatment for 48 h. **e** Western blot analysis of the expression of apoptosis-related proteins, caspase 9, 7 and PARP, following Laxiflorin B treatment for 48 h. \**P* < 0.05; \*\**P* < 0.01; \*\*\**P* < 0.001.

**Table 1.** IC_50_ of Laxiflorin B in non-small cellular lung cancer cell lines.

| Status |  | PC-9 | HCC827 | H1650 | H1975 | A549 |
| --- | --- | --- | --- | --- | --- | --- |
| EGFR |  | Ex19del | Ex19del | Ex19del | L858R,<br>T790M | WT |
| KRAS |  | WT | WT | WT | WT | KRAS<br>(G12S) |
| IC<sub>50</sub><br>(μM) | 24h | 1.503 | 2.554 | 1.148 | 3.147 | 2.298 |
|  | 48h | 1.049 | 1.161 | 0.5669 | 1.023 | 1.596 |
|  | 72h | 1.066 | 0.9231 | 0.3647 | 0.716 | 1.878 |

To assess the effects of Laxiflorin B treatment on apoptosis, PC9, HCC827 and H1650 cells were incubated with Laxiflorin B for 48 h. Flow cytometric analysis showed that apoptosis was induced in these cell lines in a dose-dependent manner (Fig. 2d). Next, we showed that the apoptotic markers, cleaved-caspase 7, 9 and PARP (Fig. 2e), were upregulated following Laxiflorin B treatment, while cleaved-caspase 3 was not (Fig. S1c and d), suggesting that Laxiflorin B-induced apoptosis was mediated via the mitochondrial pathway. Taken together, these results clearly demonstrate the inhibitory effects of Laxiflorin B on the growth and survival of NSCLC cell lines, especially in EGFR mutant subtypes.

### ERK1/2 were identified as novel targets of Laxiflorin B

We next focused on the signaling pathways and molecular mechanism underlying the effects of Laxiflorin B. We analyzed the changes in gene expression in Laxiflorin B-treated PC9 by RNA sequencing (Fig. S2a, b and c). The enrichment of biological processes (Fig. 3a) and associated pathways (Fig. 3b) in PC9 cells after Laxiflorin B treatment were classified by gene ontology and KEGG pathway analyses, respectively. Genes associated with ErbB-related signaling were significantly (*P* < 0.005) downregulated after Laxiflorin B administration (Fig. 3b and Table S1). Thus, we screened the status of ErbB-associated molecules in NSCLC cells. The phosphorylation level of members in ErbB signaling pathway were downregulated, while the levels of phospho-ERK1/2 and phospho-RSK were consistently and distinctly reduced in PC9, HCC827 and H1650 cells (Fig. 3c and S3b). The level of phosphorylated BAD, an apoptosis-induced factor ^1^ regulated by the ERK-RSK axis, was also reduced (Fig. 3c). In contrast, the stability of ERK1/2 proteins was not influenced by Laxiflorin B treatment (Fig. S3a). We also monitored the status of the p38 and STAT pathways in NSCLC cells after Laxiflorin B treatment (Fig. S3c) using DMSO treatment was as a negative control (Fig. S3d).

**Fig. 3.**
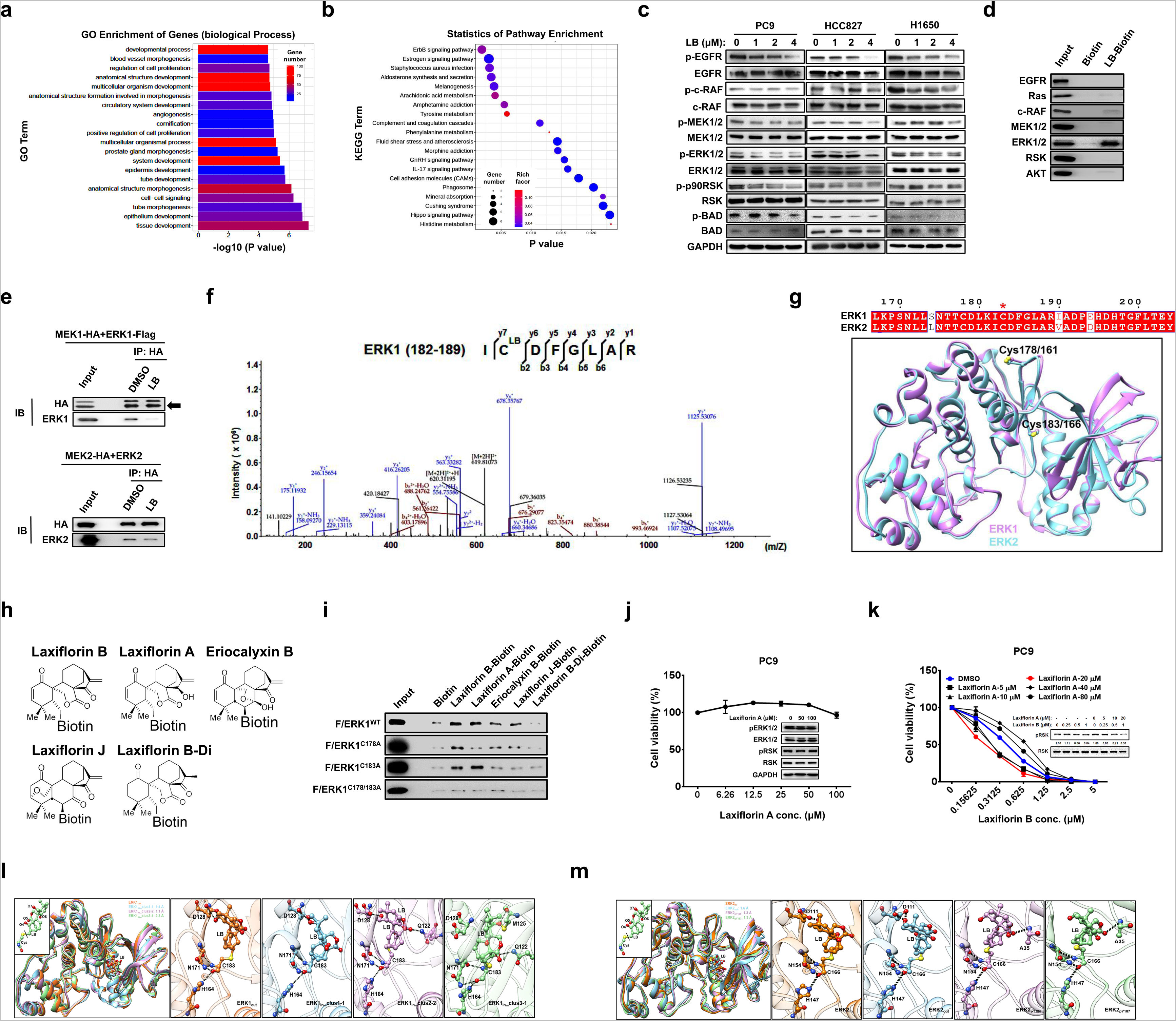
ERK1/2 as a potential novel target of Laxiflorin B. **a-b** GO and KEGG pathway analyses of the effects of 4 µM Laxiflorin B treatment for 48 h on the transcription of genes related to cellular functions and signaling pathways in PC9 cells. **c** Western blot analysis of EGFR-associated pathway proteins in PC9, HCC827 and H1650 cells following Laxiflorin B treatment for 24 h. **d** Pull-down assays of the interaction of Laxiflorin B with EGFR pathway-associated molecules. **e** Co-immunoprecipitation analysis of the the inhibitory effect of Laxiflorin B in the interaction of MEK1 with ERK1 or MEK2 with ERK2. **f** HPLC-MS/MS analysis showing that Laxiflorin B (molecular weight: 344.1624) binds covalently with ERK1 on Cys-183. **g** Structural models of ERK1 and ERK2. The sequences of the conserved Cys-178/161 and Cys-183/166 in ERK1 and ERK2 are aligned at the top of model. **h** Structure of the biotin-labeled Laxiflorin B analogues. **i** Pull-down assays of the interaction of Laxiflorin B analogues with wild-type and mutated ERK1. **j** CCK-8 assay and Western blot analysis of the cytotoxic effects of Laxiflorin A on PC9 cells. **k** CCK-8 assay and Western blot analysis of drug sensitivity and phospho-RSK expression, respectively, of PC9 cells co-treated with Laxiflorin A and B. **l** The predicted interactional model of EKR1 and Laxiflorin B from MD simulations. All representative structures are superimposed in the first panel. **m** The predicted interactional structures of EKR2 and Laxiflorin B from MD trajectories. The representative structures are superimposed in the first panel and models of the interactions between Laxiflorin B and EKR2 are shown in the following four panels. The fourth and fifth panels illustrate the interactions between Laxiflorin B and phosphorylated EKR2. The hydrogen bonds in K and L are represented by dotted lines.

Laxiflorin B treatment significantly affected the ErbB signaling pathway, indicating that Laxiflorin B directly targets key molecules involved in ErbB signal transduction. The binding of Laxiflorin B with ErbB downstream molecules was investigated using biotin pull-down assays, which showed that ERK1/2 binds Laxiflorin B with high affinity compared with other molecules (Fig. 3d). Immunoprecipitation of the HA tag on MEK1 and MEK2 showed that Laxiflorin B blocked the interaction between MEK1 and ERK1, and partially blocked the MEK2-ERK2 interaction (Fig. 3e). However, the interaction of MEK1 with ERK2 or MEK2 with ERK1 was not influenced by Laxiflorin B treatment (Fig. S3e). Furthermore, Cysteine 178 (Cys-178, Fig. S4a) and Cysteine 183 (Cys-183, Fig. 3f) of ERK1 were identified as potential binding sites for Laxiflorin B. Similarly, Cys-183 was previously reported as a covalent targeting site on ERK1^27^. The similarity of amino acid sequence (homology) between ERK1 and ERK2 reached 86% (Fig. S4b), and the amino acid sequence from Cys-161 to Cys-166 of ERK2 was identical to the sequence from Cys-178 to Cys-183 of ERK1 (Fig. 3g). Cys-161 and Cys-166 of ERK2 were also identified as potential binding sites for Laxiflorin B (Fig. 3g). Similarly, Cys-166 was previously identified as binding site for FR148083, an irreversible covalent inhibitor of ERK1/2 ^30^. More importantly, both Cys-166 of ERK2 and Cys-183 of ERK1 located within the ATP-binding pockets of ERK1 and ERK2, respectively (Fig. 3g). These results implicated ERK1/2 as the potential Laxiflorin B target proteins that mediate the inhibitory effects on the MEK-ERK axis.

To validate the significance of the chemical structure of Laxiflorin B, we investigated the structures of Laxiflorin B analogues, Laxiflorin A, Eriocalyxin B and Laxiflorin J^25,26,28^. Similar to Laxiflorin B, Eriocalyxin B retains both the A- and D-rings, although the ring junction differed from that of Laxiflorin B. In Laxiflorin A, the enone functional group of the D-ring was selectively reduced. Laxiflorin J has a similar stereochemical structure to Eriocalyxin B, although the enone functional group of the A-ring is displaced. Laxiflorin B-Di, synthesized by hydrogenation of Laxiflorin B, displaced the enone functional groups of the A- and D-rings (Fig. 3h and S5). We next conducted pull-down assays using biotin-labeled Laxiflorin B and these four analogues ( Laxiflorin A, Eriocalyxin B, Laxiflorin J and Laxiflorin B-Di) to investigate the capacity of binding wild-type (ERK1^WT^-Flag), and mutant forms (ERK1^C178A^-Flag, ERK1^C183A^-Flag, ERK1^C178A/C183A^-Flag) of ERK1 overexpressed in HEK293T cells (Fig. 3i). The results indicated that Laxiflorin B interacted strongly with ERK1^WT^, ERK1^C178A^ and ERK1^C183A^ due to its functional A- and D-rings. Laxiflorin A interacted only with ERK1^WT^ and ERK1^C183A^ due to the A-ring targeted Cys-178 on ERK1. Eriocalyxin B and Laxiflorin J, which share similar ring junctions, but differ from Laxiflorin B, interacted with ERK1 WT and ERK1 C178A, but not ERK1 C183A in pull-down assays, indicating that the D-rings played a significant role in targeting Cys-183 on ERK1. In addition, disruption of both the A- and D-rings of the Laxiflorin B-Di, or dual mutation of Cys-178 and Cys-183 on ERK1, resulted in the dissociation of Laxiflorin B analogues from ERK1 (Fig. 3i). Taken together, these results revealed that the D-ring of Laxiflorin B links covalently with the side-chain of Cys-183 in the ATP-binding pocket of ERK1 resulting in inhibition of ERK1 activity in the MEK-ERK-RSK axis of NSCLC cells.

Further, Laxiflorin A, the analogue of Laxiflorin B without the D-ring (Fig. S6a), revealed low levels of cytotoxicity against PC9 NSCLC cells (Fig. 3j and S6b and c), which was consistent with previous findings ^25^. The results from the molecular docking method and molecular dynamic (MD) simulation show that Laxiflorin A occupies the small cavity around Cys-178. The Leu132, Hie 142 and Phe146 of ERK1 provide hydrophobic environment for the binding of Laxiflorin A (Fig. S6d). Gln136, Gln149, Asn175 and Thr176 form stable hydrogen bonds with the backbone and side chain atoms in Laxiflorin A (Fig. S6e and f). However, the drug sensitivity and inhibition of the ERK-RSK axis of PC9 were promoted by co-treatment with Laxiflorin A and Laxiflorin B (Fig. 3k), which confirmed that occupation of Cys-178 by the A-ring of Laxiflorin A facilitated efficient targeting of the Cys-183 of ERK1 by the D-ring of Laxiflorin B.

Finally, we used MD simulation to generate a model of the interaction between ERK1/ERK2 and Laxiflorin B. Due to the free rotation of Laxiflorin B, Laxiflorin B exists in “in” and “out” conformations (Fig. S7a). Regardless of the conformation, the representative interactional models based on docking were found to be stable during the 200-ns unbiased MD simulation (Fig. S7b and c) in ERK1 and ERK2. The overlapped representative structures from cluster analysis showed that the interactions between Laxiflorin B and ERK1/ERK2 were almost identical (Fig. 3l and m). Laxiflorin B covalently binds to Cys-183 in ERK1 and Cys-166 in ERK2 to occupy the ATP-binding pocket and inhibit ERK activity. The probability that the backbone atoms of Cys-183/Cys-166 form several hydrogen bonds with His-164/His-147, Asn-171/Asn-154 was found to exceed 80% (Fig. S7b and c). The oxygen atom labeled O7 in A-ring also has the ability to form conserved hydrogen bonds with Asp-128/Asp-111 in ERK1/2. Furthermore, the interactions are dynamic in ERK1, where the O7 atom of A-ring and O4 atom of D-ring can also form hydrogen bonds with Met-125 and Gln-122, respectively. We also examined the effect of phosphorylation at Thr185 and Tyr-187 of ERK2 on the binding of Laxiflorin B and found that the hydrogen bonds between O7 and Asp-111 were broken, while O5 atom of B-ring could form hydrogen bonds with the backbone atoms of Ala-35 (Fig. 3m). Overall, we showed that Laxiflorin B binds covalently to Cys-183/Cys-166 of ERK1/ERK2 and expels the ATP molecule from the binding pocket, thereby inhibiting the activity of ERKs in the MEK-ERK-RSK axis of NSCLC cells.

### Laxiflorin B-induced apoptosis was mediated by BAD via ERK1/2 inhibition

Our previous data implied that Laxiflorin B-induced apoptosis in a caspase 3-independent and caspase 7-dependent manner (Fig. S1c, d and 2e), and the level of phosphorylated BAD was also reduced (Fig. 3c). Unphosphorylated BAD functions as a critical mediator of ERK-related apoptosis by promoting mitochondria-mediated cell death ^1^. We validated the status of BAD in the inhibition of the MEK-ERK axis using chemical and molecular biological approaches. Treatment of NSCLC cells with Laxiflorin B (1 µM), Trametinib (50 nM, MEK inhibitor) or Ulixertinib (1 µM, ERK inhibitor) resulted in similar reduction in the levels of phosphor-BAD (Fig. 4a). Similar effects were also observed following dual knockdown (KD) of ERK1 and ERK2 by specific shRNAs in both PC9 and HCC827 cells (Fig. S8a, b and Fig. 4b). However, KD either ERK1 or ERK2 was insufficient to diminish the level of BAD phosphorylation (Fig. 4b) in NSCLC cells.

**Fig. 4.**
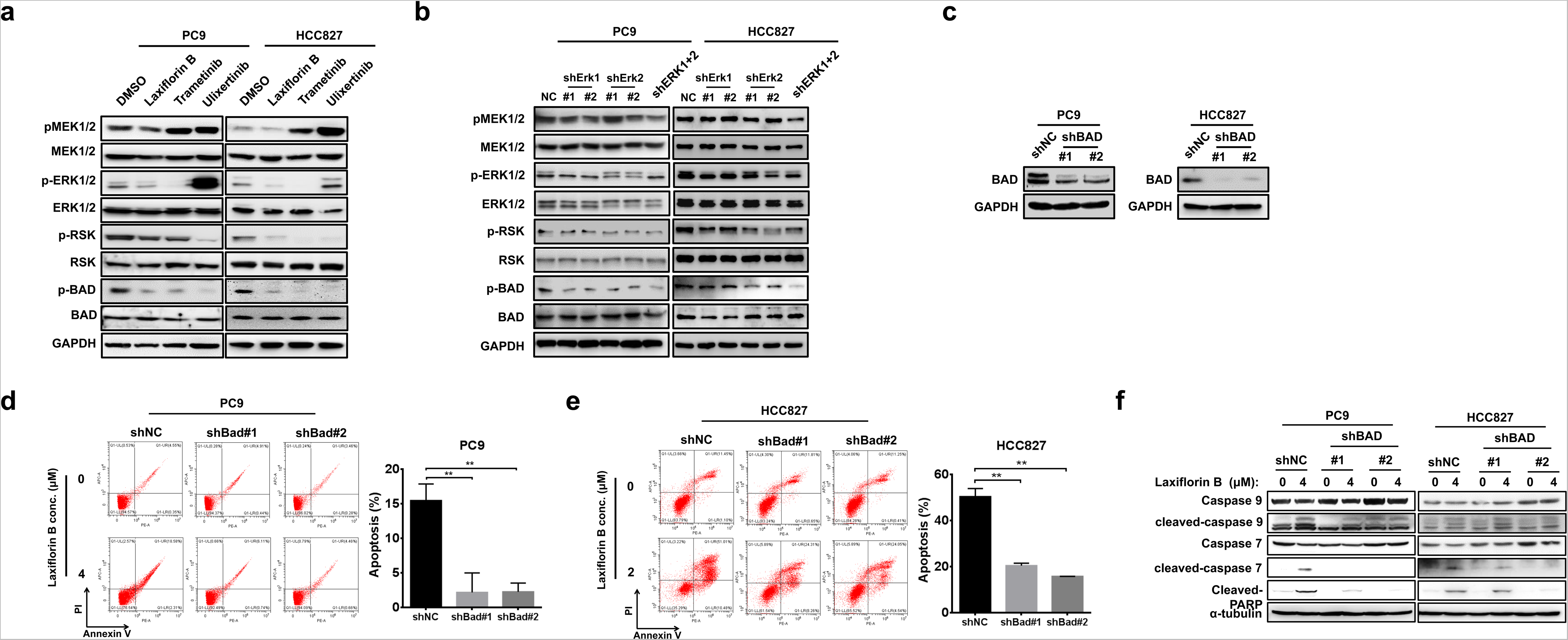
Laxiflorin B-induced apoptosis was mediated by BAD via ERK1/2 inhibition. **a** Western blot analysis of the MEK-BAD axis in PC9 and HCC827 cells after treatment with Laxiflorin B, Trametinib (a MEK inhibitor, 50 nM), and Ulixertinib (an ERK1/2 inhibitor, 100 nM) for 24 h. **b** Western blot analysis of the MEK-BAD axis-related proteins in PC9 and HCC827 cells after Laxiflorin B treatment and ERK-knockdown (KD). **c** Western blot confirmation of the efficiency of BAD-KD in PC9 and HCC827 cells. **d-e** Flow cytometric analysis of apoptosis induced in PC9 and HCC827 with stable BAD-KD stable cells by Laxiflorin B treatment for 48 h. **f** Western blot analysis of apoptotic proteins (caspase 9, 7 and PARP) in PC9 and HCC827 cells with stable BAD-KD after Laxiflorin B treatment for 48 h. \**P* < 0.05; \*\**P* < 0.01; \*\*\**P* < 0.001.

We next investigated the significance of BAD in Laxiflorin B-induced apoptosis by specific shRNA-mediated KD of BAD. The efficiency of BAD-KD was confirmed at the mRNA (Fig. S8c) and protein (Fig. 4c) levels. BAD-KD in PC9 and HCC827 cells was associated with dose- and time-dependent increases in cell viability (Fig. S8d, e, f and Table S2) and colony formation capacity (Fig. S8g and h) following Laxiflorin B treatment. Expression of cleaved-caspase 7, 9 and PARP in PC9 and HCC827 cells was effectively induced by Laxiflorin B treatment, whereas BAD-KD significantly blocked Laxiflorin B-induced apoptosis and expression of cell death markers in both PC9 and HCC827 (Fig. 4d-f). Taken together, BAD was found to play a key role in the induction of apoptosis induced by Laxiflorin B-mediated inhibition of ERK1/2.

### Autocrine production of ERK1/2-downstream growth factors was abolished by Laxiflorin B

In our previous data, we observed that Laxiflorin B treatment influenced the phosphorylation status of ERK upstream signaling molecules, such as phospho-EGFR, phospho-RAF and phospho-MEK1/2 (Fig. 3c). KEGG pathway analysis of gene differential gene expression in Laxiflorin B-treated PC9 (Fig. S2a, b and c) showed that *AREG* and *EREG* ligands of the ErbB family, were downregulated (4.41-, and 2.40-fold, respectively) after Laxiflorin B administration (Fig. S9a and Table S1). *AREG* and *EREG,* which encode the ERK-regulated growth factors ARGE and EREG, respectively, have been shown to promote cancer progression^18,29^. The expression of *AREG* and *EREG* was significantly repressed by Laxiflorin B treatment at both the mRNA (Fig. 5a and Fig. S9b) and protein (Fig. 5b) levels in NSCLC cells. In addition, conditional culture medium collected from Laxiflorin B-treated NSCLC cells did not induce EGFR activation by phosphorylation (Fig. 5c). ERK-specific shRNAs suppressed the expression of both AREG and EREG by PC9 cells in a manner that resembled the inhibitory effect of Laxiflorin B (Fig. 5d). These results demonstrated that the positive feedback loop of the ERK-driven autocrine pathway is impaired by Laxiflorin B treatment.

**Fig. 5.**
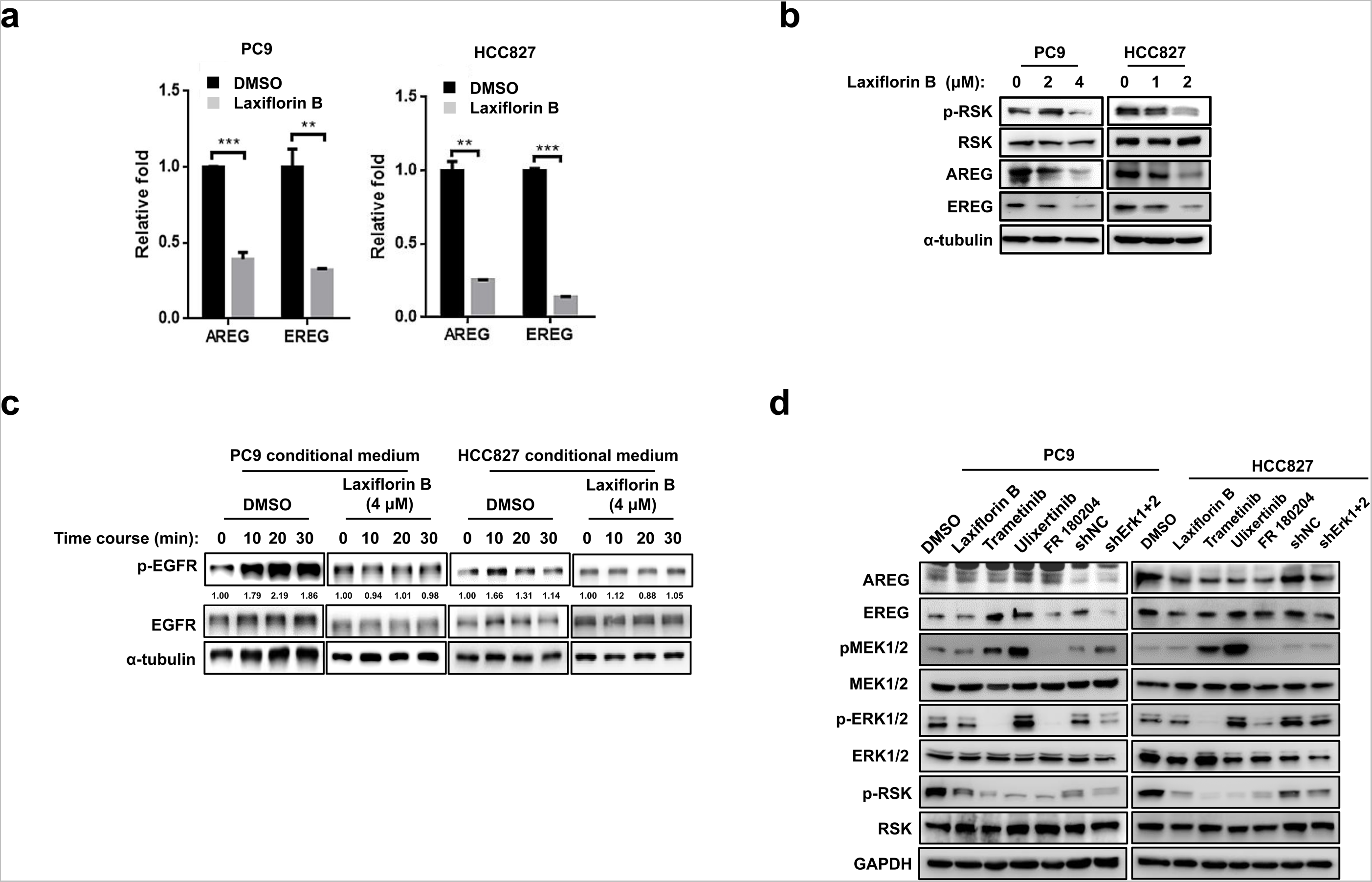
Autocrine production of ERK1/2-downstream growth factors was abolished by Laxiflorin B. **a** Real-time PCR analysis of the expression of ERK1/2 downstream growth factors after Laxiflorin B treatment for 48 h. **b** Western blot analysis of the expression of ERK downstream growth factors after Laxiflorin B treatment for 48 h. **c** Western blot analysis of phosphorylated and unphosphorylated EGFR expression in starved PC9 or HCC827 cells treated with conditioned medium collected from cultures of PC9 or HCC827 cells treated with DMSO or Laxiflorin B for 48 h. **d** Western blot analysis of components of the MEK-ERK axis in starved PC9 cells treated with conditioned medium collected from cultures of PC9 or HCC827 cells treated with Laxiflorin B, ERK shRNA, and ERK or MEK inhibitors. \*\**P* < 0.01; \*\*\**P* < 0.001.

### Laxiflorin B exerted anticancer effects in vivo

Next, we evaluated modifications of Laxiflorin B to achieve the increase in hydrophilic characteristics required to investigate its effects in a xenograft model *in vivo*. Compared with the conservative structure of Eriocalyxin B, Laxiflorin B has widely modifiable space due to its flexible structure and the presence of a primary alcohol at C-6 position (Fig. 1). Laxiflorin B-PI and Laxiflorin B-Ala were modified by the addition of piperazine and an alanine group, respectively, at the C-6 position of Laxiflorin B to increase solubility (Fig. S10a). Although cell viability assays showed the inhibitory effect of Laxiflorin B-Ala was lower than those of Laxiflorin B and Laxiflorin B-PI (Fig. S10b and Table S3), the ERK-related axis was suppressed effectively by Laxiflorin B-Ala (Fig. S10c). Considering the toxicity in the *in vivo* study, Laxiflorin B-Ala was selected as the most promising compound due to the release of the non-toxic alanine group after metabolism in mice.

We next evaluated the *in vivo* efficacy of Laxiflorin B-Ala and its therapeutic potential in PC9 cell xenograft model established in nude mice. A schematic diagram of the experimental protocol is shown in Fig. 6a. The Laxiflorin B-Ala injections in the groups receiving 10 and 20 mg/kg were terminated of days 15 and 12, respectively, due to a slight decreasing in body weight. Subsequently the body weight in these groups was restored (Fig. 6b and Fig. S10d). The tumor growth was significantly suppressed by Laxiflorin B-Ala treatment (Fig. 6b, c, d), which was also found to be associated with low toxicity (Fig. S10e). We performed IHC staining of the cell proliferation marker Ki67, phospho-RSK, AREG and EREG in the xenografts. The results showed that the expression of Ki67 and the ERK downstream markers was inhibited by Laxiflorin B-Ala treatment *in vivo* (Fig. 6e). A similar inhibitory effect was also found in PDX model (Fig S10f, g and h). These findings indicate the potential value of AREG and EREG levels as appropriate biomarkers for evaluating the efficacy of Laxiflorin B treatment *in vivo*.

**Fig. 6.**
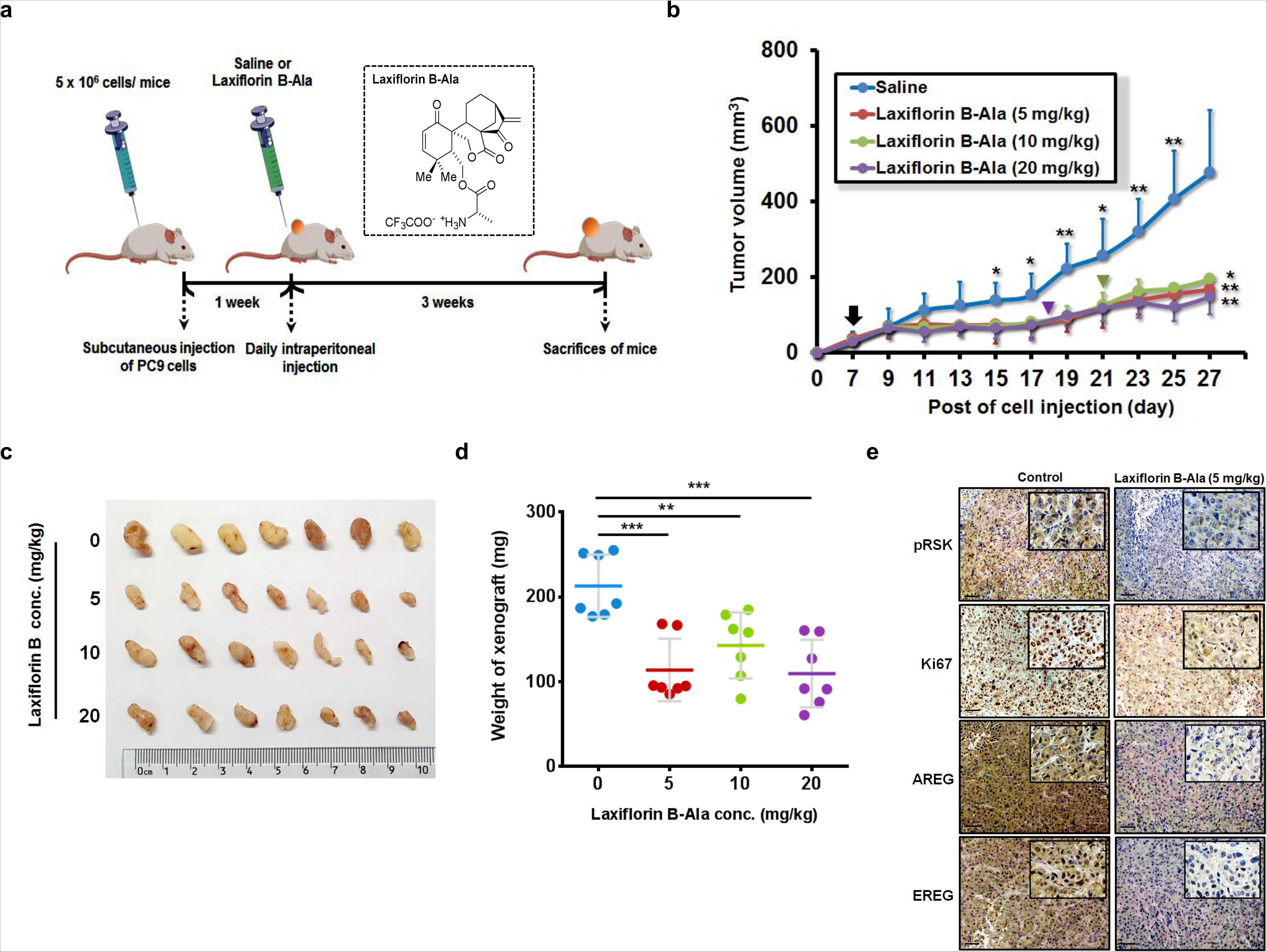
Anti-cancer effect of Laxiflorin B *in vivo*. **a** Schematic diagram of the experiment designed to evaluate the *in vivo* efficacy of Laxiflorin B-Ala and its therapeutic potential in PC9 cell xenograft model established in nude mice. **b** Tumor volume was measured once every two days post-injection of Laxiflorin B-Ala for 3 weeks. Black arrow indicates the initiation of Laxiflorin B administration, green and purple inverted triangles indicate the termination of administration in the 10 and 20 mg/kg groups on days 15 and 12, respectively. **c** Tumors harvested from the control and Laxiflorin B-Ala-treated groups. **d** Tumor weight was measured after 3 weeks of Laxiflorin B treatment. **e** Immunohistochemical staining of the expression of Ki67, phospho-RSK, AREG and EREG in xenografts. \**P* < 0.05; \*\**P* < 0.01; \*\*\**P* < 0.001.

### Evaluation of modifiable capacity and efficacy of Laxiflorin B analogues

As a 6,7-seco-*ent*-kauranoid, Laxiflorin B is suitable for modification because of less spatial hindrance of the characteristic functional group on C6. Therefore, we next modified the C-6 position hydroxyl group of Laxiflorin B to obtain more effective compounds for further investigation (Fig. 7a). Compared with unmodified Laxiflorin B, Laxiflorin B-4 and Laxiflorin B-5 had lower IC_50_ in both PC9 (Fig. 7b and Table S4) and HCC827 (Fig. S11a). We also used computational methods to understand the interactions between Laxiflorin B-4 and ERK1/2 (Fig. 7c, S11b and c). We found that the additional benzene heterocycle in Laxiflorin B-4 occupied the adenosine-binding pocket that remains vacant in the Laxiflorin B-bound state of ERK1/2. These findings confirmed that Laxiflorin B partially occupies the ATP-binding pocket of ERK1/2 ^30,31^. Chatterjee et. al. proposed that the binding of a covalent inhibitor occurs in a stepwise process, which includes non-covalent and covalent binding stages ^32^. While similar covalent bonds are formed during the binding of Laxiflorin B and Laxiflorin B-4 with ERKs, we focused on the relative binding free energy of the non-covalent binding step. As shown in Fig. S11d, the free energy of non-covalent binding of Laxiflorin B-4 with both ERK1 and ERK2 was found to be approximately 5 kcal/mol greater than that of the corresponding interactions of Laxiflorin B. The additional benzene heterocycle in Laxiflorin B-4 contributes more than 20 kcal/mol hydrophobic energy. Interestingly, for both Laxiflorin B and Laxiflorin B-4, the binding free energy of binding with ERK2 was approximately 5 kcal/mol greater than that with ERK1, which was consistent with our experimental results. Based on the non-covalent binding free energy, the proposed free energy changes during the interaction of Laxiflorin B and Laxiflorin B-4 with ERKs is depicted in Fig. 7d. Moreover, Laxiflorin B-4 prolonged the inhibition of RSK phosphorylation (Fig. 7e) and enhanced the efficiency of cell growth inhibition (Fig. 7b), which also implied that the additional functional group on Laxiflorin B-4 provides more stability for ERK1/2 targeting than Laxiflorin B (Fig. 7c). Considering the small volume of Laxiflorin B, the large binding pocket of ERK1/2 offers the opportunity for the addition of chemical groups, indicating the enormous potential for modification of Laxiflorin B to improve the binding affinity with ERK1/2.

**Fig. 7.**
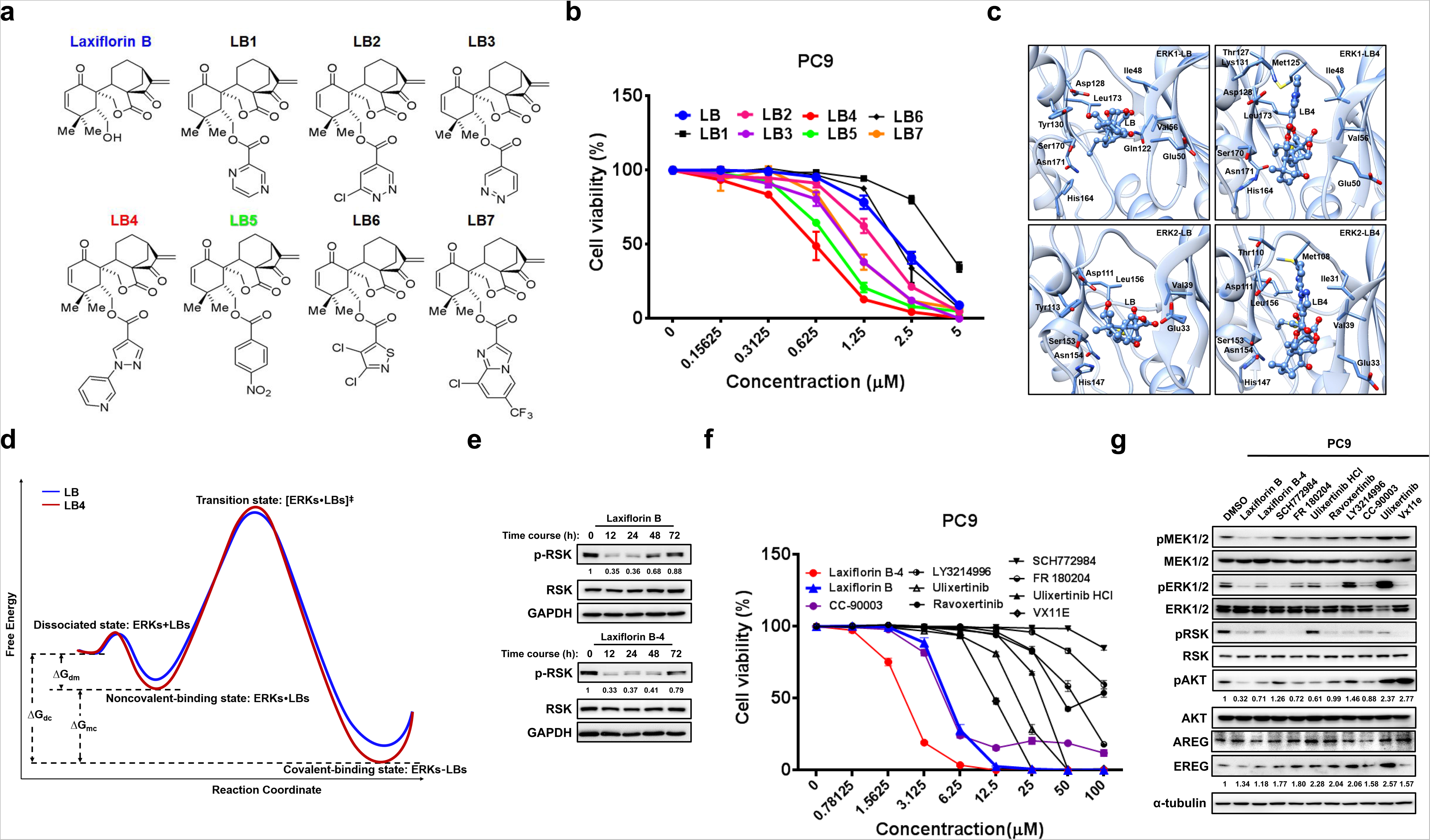
The modifiable capacity of Laxiflorin B. **a** Molecular structures of Laxiflorin B analogues modified by the addition of different functional groups. **b** IC_50_ of Laxiflorin B analogues against PC9 cells in CCK-8 assays. **c** Ball-stick model of the stable interaction between ERK1/2 and Laxiflorin B/Laxiflorin B-4. ERK1/2 is in cartoon model; the key residues interacting with Laxiflorin B and Laxiflorin B-4 are labeled. **d** The proposed free energy profile of the reaction pathway for Laxiflorin B and Laxiflorin B-4 was represented. **e** Western blot analysis of the inhibition of phosphorylated-RSK by Laxiflorin B and Laxiflorin B-4. **f** CCK-8 assay of the inhibitory effects of commercial ERK inhibitors, Laxiflorin B and Laxiflorin B-4 on PC9 cell viability. **g** Western blot analysis of the status of the MEK-ERK-RSK axis after treatment with commercial ERK inhibitors and Laxiflorin B analogues at 1 µM in PC9 cells.

Next, we selected several commercial ERK inhibitors as standards to evaluate the inhibitory effects of Laxiflorin B and Laxiflorin B-4 in NSCLC cells. Interestingly, both Laxiflorin B and Laxiflorin B-4 exhibited superior efficacy in suppressing NSCLC cell viability than the other ERK inhibitors (Fig. 7f, S11e and Table S5). Furthermore, pathway screening of NSCLC cells implied that Laxiflorin B and Laxiflorin B-4 not only suppressed RSK phosphorylation to a large extent, but also abolished AKT compensation due to a lower level of AREG and EREG than that induced by other ERK inhibitors at the same concentration (Fig. 7g and S11f).

Taken together, our findings provide evidence that, in cancer progression, the MEK-ERK-RSK axis is activated to inhibit apoptosis via the EGFR signaling cascade through BAD phosphorylation and increased production of ErbB ligands, such as AREG and EREG, leading to the maintenance of cell growth and survival. However, our results indicate that Laxiflorin B stimulates the primary response by either downregulating AREG /EREG or blocking BAD phosphorylation through covalent targeting of ERK1/2. The dephosphorylated BAD then initiates cell death via the BAD-caspase 7-PAPR axis. The secondary response is then driven by downregulation of AREG and EREG to shut down the autocrine feedback loop resulting in inhibition of EGFR signaling in NSCLC. Laxiflorin A, an analogue of Laxiflorin B, synergized with Laxiflorin B to enhance drug sensitivity of cancer cells by its non-inhibitory competition (NIC) effect. (Fig. 8).

**Fig. 8.**
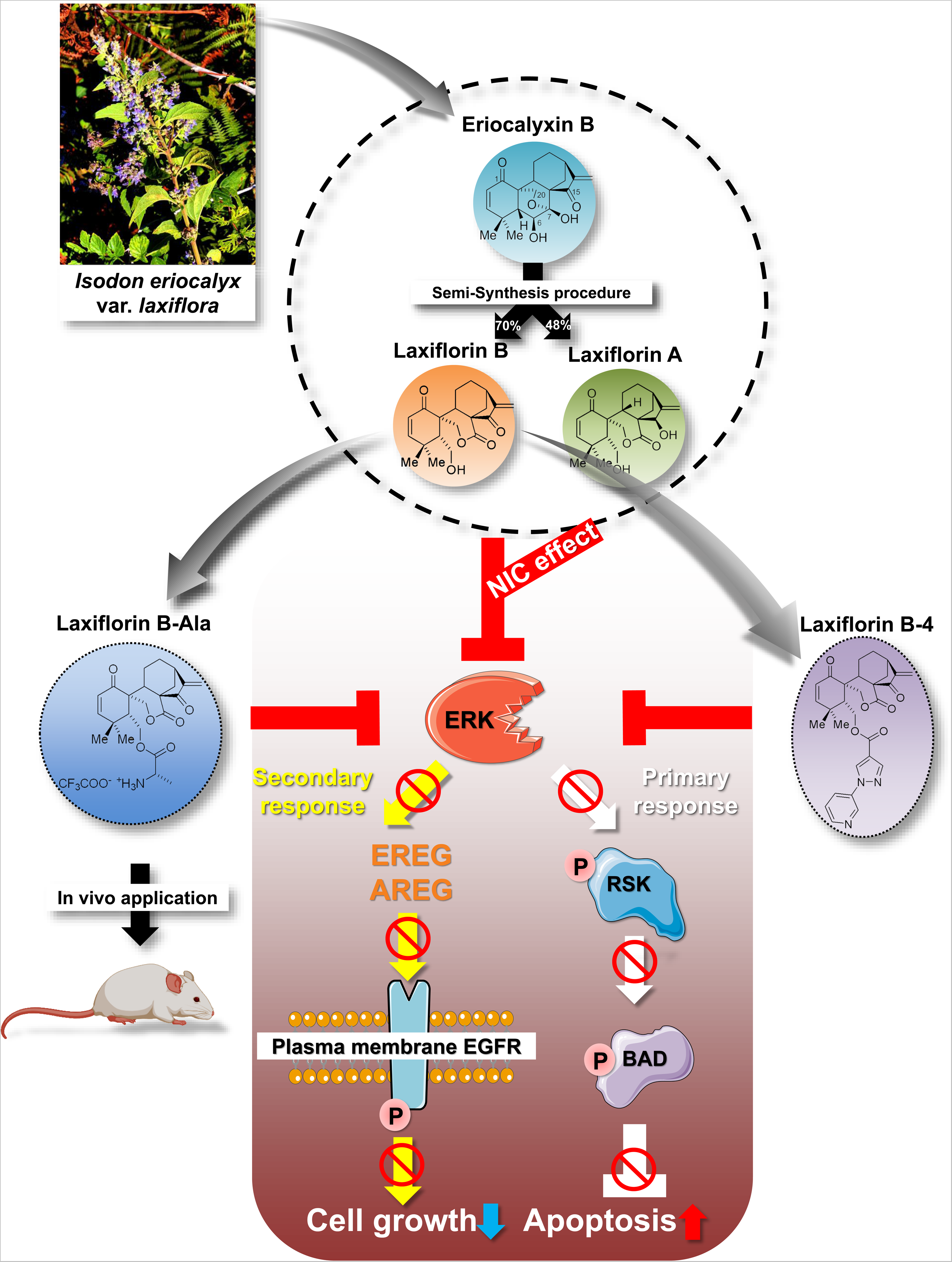
Proposed mechanism underlying the inhibitory effects of Laxiflorin B on the EGFR-related pathway in non-small-cell lung cancer.

## Discussion

In this study, we identified Laxiflorin B as the first ERK 1/2 inhibitor derived from a natural plant, and Cys-183 on ERK1 (corresponding to Cys-166 on ERK2) was identified as a potential target site of this inhibitor. The docking and MD simulation results show that Laxiflorin B occupies the ATP-binding pocket and forms stable interactions with ERK1. However, ERK1 point mutation studies revealed that approximately 50% of Laxiflorin B is degraded at the Cys-178 residue outside the ATP-binding pocket of ERK1, which highlights the potential for structural modification of Laxiflorin B to enhance pharmacologic efficacy. Simulation of the potential binding mode of the ERK2 crystal structure has also shown that Cys-166 residues (corresponding to Cys-183 on ERK1) are located in a relatively inaccessible region of the ATP-binding pocket^33^. In contrast, sulfonamides may contribute to the formation of appropriate bonds, thus highlighting the significance of Cys-166 in the ERK2 binding model. For example, FR148083^34^ and CC-90003^27^ were reported as irreversible covalent inhibitors of Cys-166 on ERK2 ^35^ and Cys-183/184 on ERK1/2, respectively. Interestingly, due to the lack of D-ring activity, Laxiflorin A binds strongly to Cys-178/161 of ERK1/2 through the A-ring, which is non-toxic to cells. We applied the reported isolation ratio of Laxiflorin A and B from herbs of 16:1 ^26^ in our co-treatment-experiment, and found that the sensitivity to Laxiflorin B treatment was enhanced by non-toxic Laxiflorin A via non-inhibitory competition in NSCLC cells. The ERK-targeting ability and lack of toxicity indicate that Laxiflorin A has important potential for development for ERK-related applications.

Human ERK1/2 are composed of 379/360 amino acids with molecular weights of 44/42 kDa and a sequence similarity of approximately 85% (higher at the ATP-binding sites). Pervious study mentioned that ERK1/2 activation is highly negatively correlated with the prognosis of cancer patients ^36^. Currently, the designation of ERK1/2 inhibitors is based mainly on modification of the existing compound skeleton. By combining a high-throughput screening platform with molecular dynamics simulation to rank a range of lead compounds for further research, a detailed model of the binding mode can be resolved at the atomic level. Among the previously reported ERK1/2 inhibitors ^37^, tetrahydropyridine, indazole and tetrahydropyrazole pyridine compounds, such as MK8353 (SCH77298414 analogue) ^40^, BVD-523 ^15^ and GDC-0994 ^17^, provide the most suitable backbones for further research, and some have shown efficacy in the preclinical stage of development^37,38^, whereas, most of ERK1/2 inhibitors are reversible and noncovalent, except FR148083^34^ and CC-90003^27^. According to kinome analysis^39^, despite plenty of kinases have cysteine residues in and around the ATP-binding site, which offer numerous opportunities to covalent reaction with compounds harboring an electrophilic Michael Acceptor in the proper position, only a few cysteine positions have thus far been targeted by covalent inhibitors. Hence, covalently binding of ERK1/2 by Laxiflorin B is an excited finding in our study, which suggests the possibility of further modification of Laxiflorin B as ERK1/2 inhibitor.

Due to the significance of the MEK-ERK axis in tumor development and progression, MEK inhibitors, such as trametinib, selumetinib and MEK162, have been widely studied in RAS- or RAF-driven cancers. As the unique downstream effector of MEK, ERK1/2 activation is critical for cancer progression, and its inhibitors, such as LY3124996 and SCH772984, have been extensively investigated in lung cancer, colorectal cancer and pancreatic cancer with KRAS mutations ^23,40^. However, inhibition of the MEK-ERK axis often results in compensatory AKT activation^41^, which is associated with negative therapeutic outcomes. Therefore, we speculate that MEK-ERK inhibition synergized with TKI or AKT inhibitors would maximize the therapeutic efficacy. In our study, we demonstrated that Laxiflorin B mediates irreversible covalent targeting of ERK1/2 to suppress its activation, leading to downregulation of its downstream target genes, such as, *AREG* and *EREG*. In turn, this inhibition attenuates not only the RAF-MEK-ERK axis, but also AKT compensation via inhibition of the positive feedback loop.

It is not difficult to understand phosphorylated-ERK1/2 plays a vital factor in cancer progression^36^. Data thus far, phosphorylated-ERK1/2 interacts with Importin7 (Imp7), facilitating its nuclear translocation, which could be blocked by myr-EPE peptide^42^. Then, nuclear ERK1/2 directly bind on specific DNA elements to influence genes expression among immune cells^43^, pluripotent embryonic stem cells^44^, and cancer cells^45,46^. Comparing with other ERKi in Fig. 7g and S11f, Laxiflorin B strongly repressed phosphorylated status of ERK1/2, which might sequentially result in inhibition of AKT compensation due to AREG and EREG suppression. Thus, Laxiflorin B is more effective than non-covalent ERK1/2 inhibitors due to the secondary feedback inhibition in addition to its primary effects. However, whether phospho-ERK1/2 is associated with ARGE and EREG expression and nuclear translocation of ERKi-bound ERK1/2 still demand further investigations. Our data suggested phospho-ERK1/2 level and ERK1/2 downstream genes expression, such as ARGE and EREG, should be included in evaluating ERKi efficacy in the future, instead of only phospho-RSK status.

Previous studies showed that expression of *AREG* and *EREG*, which encode secretory proteins, is regulated by FOSL1 ^47^ and ETS1^48^, respectively, which are transcription factors of MEK-ERK axis. High *AREG* or *EREG* expression is associated with a poor prognosis and lower survival rate in lung cancer patients, thus implicating these genes as targets for personalized cancer therapy^29,49^. In our study, AREG, EREG and AKT compensation was repressed by Laxiflorin B treatment due to ERK1/2 inhibition. Therefore, it can be speculated that direct detection of AREG or EREG levels in blood or body fluids represents a valuable marker to evaluate the efficacies of Laxiflorin B and other ERK inhibitors in future clinical applications, although EREG might be as a more accurate indicator because AREG can be adsorbed by extracellular matrix heparin sulfate proteoglycans in the extracellular space ^50^.

Laxiflorin B is a liposoluble molecule with low solubility in normal saline for peritoneal injection. To improve the water-solubility, alanine trifluoroacetate was used to modify C-6 hydroxyl group of Laxiflorin B through esterification to generate Laxiflorin B-Ala. This modification facilitated pharmacodynamic evaluation of Laxiflorin B *in vivo*, despite its slightly inferior inhibitory activity *in vitro*. After diffusing into the plasma, the C-6 position ester group of Laxiflorin B-Ala is hydrolyzed by esterase, resulting in Laxiflorin B release. Since alanine trifluoroacetate accounts for 35% of the molecular weight of Laxiflorin B-Ala, the actual concentration of Laxiflorin B in our model was far below than that of Laxiflorin B-Ala.

Laxiflorin B was the first covalent-natural compound shown to exhibit ERK-targeting and dramatic anticancer activities. Although previous studies have shown that Laxiflorin A, an analogue of Laxiflorin B, has little anti-tumor activity ^25^, our clarification of the molecular mechanism has redefined the value of Laxiflorin A for this purpose. First, since Laxiflorin A binds to Cys-178/161 on ERK1, removal of A-ring activity on Laxiflorin B might minimize the ineffective binding to ERK1/2. Therefore, the drug sensitivity of tumor cells could be promoted by combined treatment with Laxiflorin A and B. Moreover, as a molecular probe of ERK, Laxiflorin A might offer great potential for the design of compounds for further research or clinical applications. The synergistic activity of multiple components of herbs has long been a focus of research into their clinical application^51^. This phenomenon, which is consistent with the compound concept of traditional Chinese medicine, first prompted us to propose the existence of non-inhibitory competition (NIC) between the herbal compounds Laxiflorin A and Laxiflorin B. This type of interaction between natural small molecules might greatly improve the efficacy of active ingredients in natural substances.

In this study, we have developed comprehensive procedures for the synthesis and modification of Laxiflorin B and validated its activity as an ERK1/2 inhibitor both *in vitro and in vivo*. With the assistance of computer simulation and the immense potential for modification, we propose that Laxiflorin B represents an important compound that can be manipulated to improve specificity and affinity. As a novel ERK inhibitor for suppressing MEK-ERK-dependent malignancies, Laxiflorin B might have potential of medication alone or in combination with TKI or other inhibitors for both antitumor therapy and addressing resistance in the future. Our finding also provides a reference for further investigation of the anticancer effects of compounds derived from herbs.

## Material and Methods

### General coumpound synthesis

Unless otherwise noted, all air- and water-sensitive reactions were carried out under nitrogen and with dry solvents under anhydrous conditions, respectively. Reactions were monitored by thin-layer chromatography (TLC) on 0.25 mm silica gel plates (60F-254) that were analyzed by fluorescence at 254 nm or by staining with KMnO_4_ (200 mL H_2_O containing 1.5 g KMnO_4_, 10 g K_2_CO_3_, and 1.25 mL 10% aqueous NaOH). Silica gel 60 (particle size 0.040–0.063 mm) was used for flash column chromatography.

All the chemicals were purchased commercially and used without further purification. Anhydrous solvents were distilled according to standard procedures. Details of general methods, compound characterization, HPLC analysis, and ^1^H and ^13^C NMR spectra of the compounds prepared in this study are provided in the Supplementary Materials and Methods.

### Reagents, enzymes, and antibodies

Chemical reagents and reaction buffers were obtained from Sigma–Aldrich (St. Louis, MO, USA). Q5^®^ High-Fidelity DNA Polymerase, DNA ligase and the restriction enzymes were purchased from New England BioLabs (NEB, Ipswich, MA, USA). A list of antibodies used is provided in the Supplementary Materials and Methods.

### Construction of plasmids and mutagenesis

pBABEpuro-HA-MEK1 (#49328), pMM9-MAPKK2-WT (MEK2, #40805), pFLAG-CMV-hERK1 (#49328), pHAGE-MAPK1 (ERK2, #116760) were purchased from Addgene (MA, USA). Point mutations were introduced using Q5 DNA polymerase to generate the ERK1^C178A^, ERK1^C183A^ and ERK1^C178A/C183A^ mutants. The primary DNA template was digested with *Dpn*I for 2 h at 37°C before DH5α competent cells were transformed with mutant ERK1 to amplify the construct. The sequences were confirmed by Sanger sequencing. Details of the primers used for mutagenesis are provided in the Supplementary Materials and Methods.

### Cell culture and stable pool selection

HEK293T cells were obtained from the Cell Bank of Chinese Academy of Sciences and cultured in Dulbecco’s modified Eagle’s medium (DMEM; Life Technologies) supplemented with 10% fetal bovine serum (FBS; PAN Seratech, Aidenbach, Germany) and 100 U/ml penicillin-streptomycin (Life Technologies). The human non-small-cell lung cancer (NSCLC) cell lines PC9, HCC827, A549, NCI-H1975 and NCI-H1650 were obtained from the American Type Culture Collection (ATCC) and cultured in RPMI1640 (Life Technologies) containing 10% FBS (PAN Seratech) and 100 U/mL penicillin-streptomycin (Life Technologies). HEK293T, PC9, HCC827, A549, NCI-H1975, NCI-H1650 and stable pools were incubated at 37°C under 5% CO_2_. All cell lines were authenticated by short tandem repeat (STR) profiling (Guangdong Huaxi Forensic Physical Evidence Judicial Appraisal Institute, Guangdong, China). The cell lines were passaged fewer than 10 times or for a maximum of 6 months after initial revival from frozen stocks; all cell lines tested negative for mycoplasma.

For generation of stable pools with BAD or ERK1/2 knockdown, NSCLC cell lines were infected with lentiviruses expressing BAD or ERK1/2 shRNA and selected by culturing in RPMI1640 complete medium containing 10 µg/mL puromycin (Selleck, TX, USA) for 1 week. BAD or ERK1/2 expression was evaluated by qPCR and Western blot analyses. Functional assays were performed after confirmation of gene expression. Details of the shRNA oligonucleotides are provided in the Supplementary Materials and Methods.

### Real-time PCR

Total RNA was extracted using TRIzol reagent (Life Technologies) and cDNA was synthesized using RevertAid First Strand cDNA Synthesis Kit (Thermo Scientific, Waltham, MA, USA) according to the manufacturer’s instructions. All real-time PCRs were performed using the CFX Connect Real-Time PCR Detection System (Bio-Rad, Hercules, CA, USA) and the amplifications were carried out using the ChamQ Universal SYBR qPCR Master Mix (Vazyme, Nanjing, China). Each real-time PCR reaction was repeated three times, and target genes were normalized using GAPDH as an internal reference. Details of the primers used for real-time PCR are shown in the Supplementary Materials and Methods.

### Immunoprecipitation, pull-down assays and Western blot analysis

For immunoprecipitation (IP), HEK293T cells transfected with the indicated plasmids were cultured in a 100-mm dishes. Cells were collected at 80% confluence using IP lysis buffer. Following clarification, the supernatant fractions were immunoprecipitated by incubation with magnetic beads conjugated with anti-HA or anti-Flag antibodies at 4°C overnight with rotation at 5 rpm (MedChemExpress, Monmouth Junction, NJ, USA). After washing the magnetic beads several times with wash buffer (25 mM Tris-HCl, pH 7.5, 100 mM NaCl, 0.1% NP-40), the immunoprecipitate was finally eluted with sample buffer containing 1% SDS.

For pull-down assays, Laxiflorin B-biotin was conjugated with streptavidin magnetic beads and then incubated with total lysate from HEK293T or PC9 cells harvested at 80% confluence using IP lysis buffer. After washing the magnetic beads several times with wash buffer (25 mM Tris-HCl, pH 7.5, 100 mM NaCl, 0.05% Triton-X100), the immunoprecipitate was finally eluted with sample buffer containing 1% SDS.

Western blotting was performed following IP. The protein concentration was determined using a BCA Protein Assay Kit (Beyotime Biotechnology, Shanghai, China). All protein samples were separated by sodium dodecyl sulfate-polyacrylamide gel electrophoresis (SDS-PAGE), transferred to polyvinylidene fluoride (PVDF) membranes (Merck Millipore, Burlington, MA, USA), and hybridized with the corresponding antibodies. IP bands were visualized using Pierce ECL Western blotting substrate (Thermo Scientific) and detected by Fluor Chem Q (ProteinSimple, USA). AlphaEaseFC version 4.0 software was applied for quantifying the raw data of western blotting.

### Mass spectrometry (MS) analysis of Laxiflorin B binding with ERK1

HEK293T cells overexpressing a Flag-ERK1 fusion protein were purified using magnetic beads conjugated with an anti-Flag antibody (MedChemExpress, Monmouth Junction, NJ, USA) and incubated with 10 µM Laxiflorin B for 16 h at 4°C. The beads samples were then washed three times with cold PBS containing 0.05% TritonX-100 before incubation with the reaction buffer (1% SDC/100 mM Tris-HCl, pH 8.5/10 mM TCEP/40 mM CAA) at 95°C for 10 min for protein denaturation, cysteine reduction and alkylation. The eluates were diluted with an equal volume of H_2_O and subjected to trypsin digestion overnight at 37°C following the addition of 1 μg trypsin. The peptide was purified using self-made SDB desalting columns. The eluate was vacuum-dried and stored at -20°C for later use.

Mass spectrometry was performed using a TripleTOF 5600+ LC-MS/MS system (SCIEX). The peptide sample was injected using an autosampler and bound to a C18 capture column (5 μm, 5 × 0.3 mm), followed by elution to the analytical column (300 μm × 150 mm, 3 μm particle size, 120 Å pore size, Eksigent) for separation. Two mobile phases (mobile phase A: 3% DMSO, 97% H_2_O, 0.1% formic acid and mobile phase B: 3% DMSO, 97% ACN, 0.1% formic acid) were used to establish the analytical gradient. The flow rate of the liquid phase was set to 5 μL/min. For mass spectrometry IDA mode analysis, each scan cycle involved a full MS scan (m/z range 3501250, ion accumulation time 250 ms), followed by 40 MS/MS scans (m/z range 100–1500, ion accumulation time 50 ms). The conditions for MS/MS acquisition were set to a parent ion signal >120 cps and a charge number of +2 to +5. The exclusion time for ion repeat acquisition was set to 18 s.

Raw data generated by the TripleTOF 5600 were analyzed with ProteinPilot (V4.5) using the Paragon database search algorithm and the integrated false discovery rate (FDR) analysis function. Spectra files were searched against the UniProt Human reference proteome database using the following parameters: sample type, identification; Cys alkylation, iodoacetamide; digestion, trypsin; search effort, rapid ID. Search results were filtered with unused ≥1.3. Decoy hits and protein contaminants were removed; the remaining identifications were used for further analysis.

### Cell growth, 2-D clonogenic and soft assays

For analysis of cell growth, cells were seeded in 96-well plates at 5 × 10^3^ cells per well in a final volume of 100 µL culture medium and incubated at 37°C. Cell viability was evaluated at 0, 24, 48, and 72 h after cell attachment. Cell Counting Kit-8 (CCK-8, Meilun Biotechnology, China) reagent (10 µL) was added to each well, and the plates were incubated for 2 h at 37°C. The absorbance was measured at 450 nm by Maestronano spectrophotometer (TriStar2LB942, Berthold, Germany).

For 2-D clonogenic assays, cells were seeded in 6-well plates at 500 cells per well in a final volume of 4 mL culture medium and incubated at 37°C. After incubation for 8 to 10 days, the 6-well plates were washed with 1× PBS before fixation in 100% methanol at 4°C for 10 min. The cells were then stained with 0.5% crystal violet for 10 min at room temperature before being washed twice in double-distilled (dd) H_2_O.

For soft agar assays, cells were seeded in 0.4% low-melting agarose (Sigma–Aldrich) in 6-well plates at 5×10^3^ cells per well. The cells were incubated at 37°C for 2–3 weeks, then stained with 0.005% crystal violet for 1 h at room temperature before being destained in ddH_2_O.

### Molecular docking and molecular dynamic simulation

In this study, the HADDOCK server ^52^ was used to simulate covalent docking of Laxiflorin B analogues onto ERK1/2. Both the “in” and “out” conformations of Laxiflorin B analogues were studied. The A chain of crystal structures 6GES ^53^ and 6G54 ^53^ were used as the templates of ERK1 and ERK2 for docking with Laxiflorin B analogues. During docking, the clustering method was based on RMSD and the cutoff was set to 1 Å. For scoring parameters, Evdw 1 and Eelec 3 were changed to 1.0 and 0.1, respectively. Several advanced sampling parameters were reset. The initial temperature for the second and third TAD cooling step with flexible side-chain and full flexibility at the interface were set as 500 K and 300 K, respectively. The numbers of molecular dynamic (MD) steps for rigid body high temperature TAD and during the first rigid body cooling stage were both set as 0. All other parameters were unchanged. For each docked complex, the representative structure was selected as the initial structure of MD simulation.

All MD simulations were performed in the AMBER package^54^. The force field FF14SB^55^ was selected to build the force field of standard amino acids in proteins. The parameters for Laxiflorin B analogues were prepared based on the General AMBER force field (GAFF) ^56^. The atomic charges of Laxiflorin B analogues were fitted based on the electrostatic potential (ESP) determined by quantum calculation. In accordance to the atomic charge from the FF14SB force field, the HF-6-31G* basis set was selected to obtain the ESP. Each docked complex was then immersed in a TIP3P water box ^57^ (size 10 Å). The counter ions were added to neutralize the system, which included approximately 50,000 atoms in total. The whole system was firstly minimized to convergence. After minimization, all heavy atoms were restrained by force constant 100 kcal/(mol·rad^2^), and the temperature was gradually increased to 300 K in the NVT ensemble. The whole system was then relaxed in the NPT ensemble without restraint. The duration of the simulation was 200 ns. A snapshot was collected every 1 ps. All analyses were conducted using AMBER in-built modules and in-house code. The cluster analysis was performed using the linkage method to select the representative structure. The cutoff was set to 0.8 Å.

The noncovalent binding free energy was calculated using the molecular mechanics generalized Born surface area (MM-GBSA) method ^58^. The entropic contribution was not considered in the total binding energy; therefore, the total binding free energy between Laxiflorin B analogues and ERK1/2 included only the electrostatic contribution (EEL), the van der Waals (VDW) interaction contribution in the gaseous phase and the solvation free energy equal to the sum of the polar (EGB) and nonpolar (ESURF) contributions.

### Cell apoptosis analysis

5 x 10^5^ PC9 and HCC827 cells were seeded into 6-well plates. After 24 h culturing at 37°C incubator, the attached cells were treated with Laxiflorin B at 0, 1, 2 and 4 μM for 48 h. The cells were then stained with Annexin V and PI using the Annexin V-FITC Apoptosis Detection Kit (BioVision, USA) and analyzed using the FACSCalibur platform (BD Biosciences).

### Xenografts in nude mice

All animal research procedures were performed according to the rules of the Animal Care and Use Ethics Committee of Shenzhen University Health Science Center, and all animals were treated in strict accordance with protocols approved by the Institutional Animal Use Committee of the Health Science Center, Shenzhen University. Female *nu*/*nu* nude mice (aged 4–6 weeks) were subcutaneously injected in the dorsal region with 5×10^6^ PC9 cells. When the tumor volume reached 50 mm^3^ (after 7 days), saline and Laxiflorin B-Ala (5, 10, 20 mg/kg) were injected intraperitoneally once per day. Tumor volume and mouse body weight were measured once every two days. Tumor volume was calculated according to the formula: volume = length × width^2^/2. After 3 weeks of Laxiflorin B treatment, mice were euthanized, and their tumor xenografts were harvested for weighing and immunohistochemical staining for phospho-RSK, AREG, EREG, and Ki-67

### Statistical analyses

All *in vitro* experiments were performed at least in triplicate. The results of each experiment were presented as the mean ± standard deviation. Data analyses were performed using Microsoft Excel 2010 Professional Plus (Version 14.0.7237.5000) and GraphPad Prism 6 (Version 6.01). Two-tailed, unpaired Student’s *t*-tests were used to compare differences between two groups with similar variances. For all tests, a *P*-value <0.05 was considered to indicate a statistically significant difference.

## Authors’ contributions

Conceived and designed the experiments: Zheng Duo, Zhu Lizhi and Chiang Chengyao. Chemical compounds synthesis: Huang Junrong, Yang Min. Carried out most of the experiments and analyzed the data: Zhang Min, Chiang Chengyao, Huang Junrong, Chen Chunlan, Pan Dongmei, Heng Yang. Construction of plasmids and lentivirus packaging: Zhang Min, Chiang Chengyao, Chen Chunlan. Animal studies: Chiang Chengyao and Yin Feng. Performed the LC-MS/MS and proteomic analysis: Han Qiangqiang, Wangzou, Zou Yongdong. Performed the computer simulation of Laxiflorin B and ERK1/2 crystal structures: Zeng Juan. Planning and discussion of the project: Zheng Duo, Zhu Lizhi. Supervised the entire project: Zheng Duo, Zhu Lizhi. Wrote the manuscript, designed the layout of Fig.s and tables: Chiang Chengyao, Zheng Duo, Li Zigang, Zeng Juan.

## Acknowledgments

The authors would like to thank Jessica Tamanini of ETediting for editing the manuscript prior to submission and the Instrumental Analysis Center of Shenzhen University for the support. This work was supported by the Natural Science Foundation of Guangdong Province (No. 2017A030313042, 2019A1515010210, 2021A1515011046, 2021A1515011154, and 2021A1515010996), the Guangdong Provincial Science and Technology Program (No. 2017B030301016), the Shenzhen Municipal Government of China (No. JCYJ20180507182427559 and JCYJ20210324093408024), the Shenzhen Key Medical Discipline Construction Fund (No. SZXK060).

