## Supplementary Materials and Methods for "A Novel Selective ERK1/2 Inhibitor, Laxiflorin B, Targets EGFR Mutation Subtypes in Non-small-cell Lung Cancer"

### List of Contents:

**Antibodies list**
**Oligo-nucleutide list**
**Synthesis procedure of Laxiflorin B analogues**

**Antibodies list**

| Antibody | Cat. no. | Supplier | Dilution factor |
|---|---|---|---|
| HA-tag | sc-7392 | Santa Cruz Biotechnology (Dallas, TX, USA) | 1:2000 |
| Flag-tag | F1804 | Sigma-Aldrich (St. Louis, MO, USA) | 1:2000 |
| β-actin | CW0096A | CoWin BioSciences (Cambridge, MA, USA) | 1:2000 |
| $\alpha$-tubulin | 11224-1-AP | ProteinTech Group, Inc. (Rosemont, IL, USA) | 1:2000 |
| phospho-Akt | 4060 | Cell Signaling Techonlogy (Danvers, MA, USA) | 1:1000 |
| Akt | 9272 | Cell Signaling Techonlogy (Danvers, MA, USA) | 1:1000 |
| phospho-Erk1/2 | 4070 | Cell Signaling Techonlogy (Danvers, MA, USA) | 1:1000 |
| Erk1/2 | 4695 | Cell Signaling Techonlogy (Danvers, MA, USA) | 1:1000 |
| Ki67 | GB13030-2 | Servicebio (Wuhan, China) | 1:200 |
| Phosphor-EGFR | 3777 | Cell Signaling Techonlogy (Danvers, MA, USA) | 1:1000 |
| EGFR | 4267 | Cell Signaling Techonlogy (Danvers, MA, USA) | 1:1000 |
| Phosphor-MEK1/2 | 9154 | Cell Signaling Techonlogy (Danvers, MA, USA) | 1:1000 |
| MEK1/2 | 9122 | Cell Signaling Techonlogy (Danvers, MA, USA) | 1:1000 |
| Phosphor-c-Raf | 9427 | Cell Signaling Techonlogy (Danvers, MA, USA) | 1:1000 |
| c-Raf | 53745 | Cell Signaling Techonlogy (Danvers, MA, USA) | 1:1000 |
| Phosphor-p90RSK | 11989 | Cell Signaling Techonlogy (Danvers, MA, USA) | 1:1000 |
| RSK | 9355 | Cell Signaling Techonlogy (Danvers, MA, USA) | 1:1000 |
| GAPDH | 60004-1-Ig | ProteinTech Group, Inc. (Rosemont, IL, USA) | 1:2000 |
| Phosphor-Bad | 9291 | Cell Signaling Techonlogy (Danvers, MA, USA) | 1:500 |
| Bad | 9268 | Cell Signaling Techonlogy (Danvers, MA, USA) | 1:500 |
| Ras | ab52939 | Abcam (Cambridge, England) | 1:1000 |
| Amphiregulin | GTX100986 | GeneTex (Southern California, USA) | 1:250 |
| Epiregulin | GTX16256 | GeneTex (Southern California, USA) | 1:250 |
| Caspase 3 | 9665 | Cell Signaling Techonlogy (Danvers, MA, USA) | 1:1000 |
| Caspase 7 | 9492 | Cell Signaling Techonlogy (Danvers, MA, USA) | 1:1000 |
| Caspase 9 | 9502 | Cell Signaling Techonlogy (Danvers, MA, USA) | 1:1000 |
| Cleaved-PARP | 5625 | Cell Signaling Techonlogy (Danvers, MA, USA) | 1:1000 |
| Cyclin D1 | 55506 | Cell Signaling Techonlogy (Danvers, MA, USA) | 1:1000 |
| p21 | Ab7960 | Abcam (Cambridge, England) | 1:500 |
| p27 | 3686 | Cell Signaling Techonlogy (Danvers, MA, USA) | 1:1000 |
| Phosphor-p38 | 4511 | Cell Signaling Techonlogy (Danvers, MA, USA) | 1:1000 |
| p38 | 8690 | Cell Signaling Techonlogy (Danvers, MA, USA) | 1:1000 |
| Phosphor-STAT3 | 9145 | Cell Signaling Techonlogy (Danvers, MA, USA) | 1:1000 |
| STAT3 | 9139 | Cell Signaling Techonlogy (Danvers, MA, USA) | 1:1000 |

| | | | |
|---|---|---|---|
| p85 | 4257 | Cell Signaling Techonlogy (Danvers, MA, USA) | 1:1000 |
| anti-rabbit | 111-035-008 | Jackson ImmunoResearch Laboratories, Inc (West Grove, Pennsylvania, USA) | 1:5000 |
| anti-mouse | 115-035-008 | Jackson ImmunoResearch Laboratories, Inc (West Grove, Pennsylvania, USA) | 1:5000 |

#### Oligo-nucleutide list

| **Cloning primers** | |
|---|---|
| Erk1 C178A-Forward | 5’- ACCTGCTCAGCAACACCACCGCCGACCTTAAGATTTGTGATT -3’ |
| Erk1 C178A-Reverse | 5’- AATCACAAATCTTAAGGTCGGCGGTGGTGTTGCTGAGCAGGT-3’ |
| Erk1 C183A-Forward | 5’- CCACCTGCGACCTTAAGATTGCTGATTTCGGCCTGGCCCGGA -3’ |
| Erk1 C183A-Reverse | 5’- TCCGGGCCAGGCCGAAATCAGCAATCTTAAGGTCGCAGGTGG-3’ |
| Erk1 C178/183A-Forward | 5’- AGCAACACCACCGCCGCCCTTAAGATTGCTGATTTCGGCCTG-3’ |
| Erk1 C178/183A-Reverse | 5’- CAGGCCGAAATCAGCAATCTTAAGGTCGGCGGTGGTGTTGCT-3’ |
| **qPCR primers** | |
| AREG-Forward | 5’-CCACAGTGCTGATGGATTTG-3’ |
| AREG-Reverse | 5’-AGCCAGGTATTTGTGGTTCG-3’ |
| EREG-Forward | 5’-TCCCAGGAGAGTCCAGTGAT-3’ |
| EREG-Reverse | 5’-AGTGTTCACATCGGACACCA-3’ |
| HBEGF-Forward | 5’-GGTGGTGCTGAAGCTCTTTC-3’ |
| HBEGF-Reverse | 5’-GCTTGTGGCTTGGAGGATAA-3’ |
| Bad-Forward | 5’-CCGAGTGAGCAGGAAGACTC-3’ |
| Bad-Reverse | 5’- GGTAGGAGCTGTGGCGACT-3’ |
| Erk1-Forward | 5’- ACAGTCTCTGCCCTCCAAGA-3’ |
| Erk1-Reverse | 5’-CTCATCCGTCGGGTCATAGT-3’ |
| Erk2-Forward | 5’-CCAGACCATGATCACACAGG-3’ |
| Erk2-Reverse | 5’-CTGGAAAGATGGGCCTGTTA-3’ |
| GAPDH-Forward | 5’-GTCTCCTCTGACTTCAACAGCG-3’ |
| GAPDH-Reverse | 5’-ACCACCCTGTTGCTGTAGCCAA-3’ |
| **Oligo-nucleutide for knockdown** | |
| shBad-#1-Top | 5’-CCGGGAGGATGAGTGACGAGTTTGTCTCGAGACAAACTCGTCACTCATCCTCTTTTTG-3’ |
| shBad-#1-Bottom | 5’-AATTCAAAAAGAGGATGAGTGACGAGTTTGTCTCGAGACAAACTCGTCACTCATCCTC-3’ |
| shBad-#2-Top | 5’-CCGGGTTTGTGGACTCCTTTAAGAACTCGAGTTCTTAAAGGAGTCCACAAACTTTTTG-3’ |
| shBad-#2-Bottom | 5’-AATTCAAAAAGTTTGTGGACTCCTTTAAGAACTCGAGTTCTTAAAGGAGTCCACAAAC-3’ |
| shErk1-#1-Top | 5’-CCGGGCCATGAGAGATGTCTACATTCTCGAGAATGTAGACATCTCTCATGGCTTTTTG-3’ |

| | |
|---|---|
| shErk1-#1-Bottom | 5’-AATTCAAAAAGCCATGAGAGATGTCTACATTCTCGAGAATGTAGACATCTCTCATGGC-3’ |
| shErk1-#2-Top | 5’-CCGGGGATCAGCTCAACCACATTCTCTCGAGAGAATGTGGTTGAGCTGATCCTTTTTG-3’ |
| shErk1-#2-Bottom | 5’-AATTCAAAAAGGATCAGCTCAACCACATTCTCTCGAGAGAATGTGGTTGAGCTGATCC-3’ |
| shErk2-#1-Top | 5’-CCGGGGACCTCATGGAAACAGATCTCTCGAGAGATCTGTTTCCATGAGGTCCTTTTTG-3’ |
| shErk2-#1-Bottom | 5’-AATTCAAAAAGGACCTCATGGAAACAGATCTCTCGAGAGATCTGTTTCCATGAGGTCC-3’ |
| shErk2-#2-Top | 5’-CCGGGCACCATTCAAGTTCGACATGCTCGAGCATGTCGAACTTGAATGGTGCTTTTTG-3’ |
| shErk2-#2-Bottom | 5’-AATTCAAAAAGCACCATTCAAGTTCGACATGCTCGAGCATGTCGAACTTGAATGGTGC-3’ |

### Synthesis procedure of Laxiflorin B analogues

#### Semi-synthesis of Laxiflorin A and Laxiflorin B

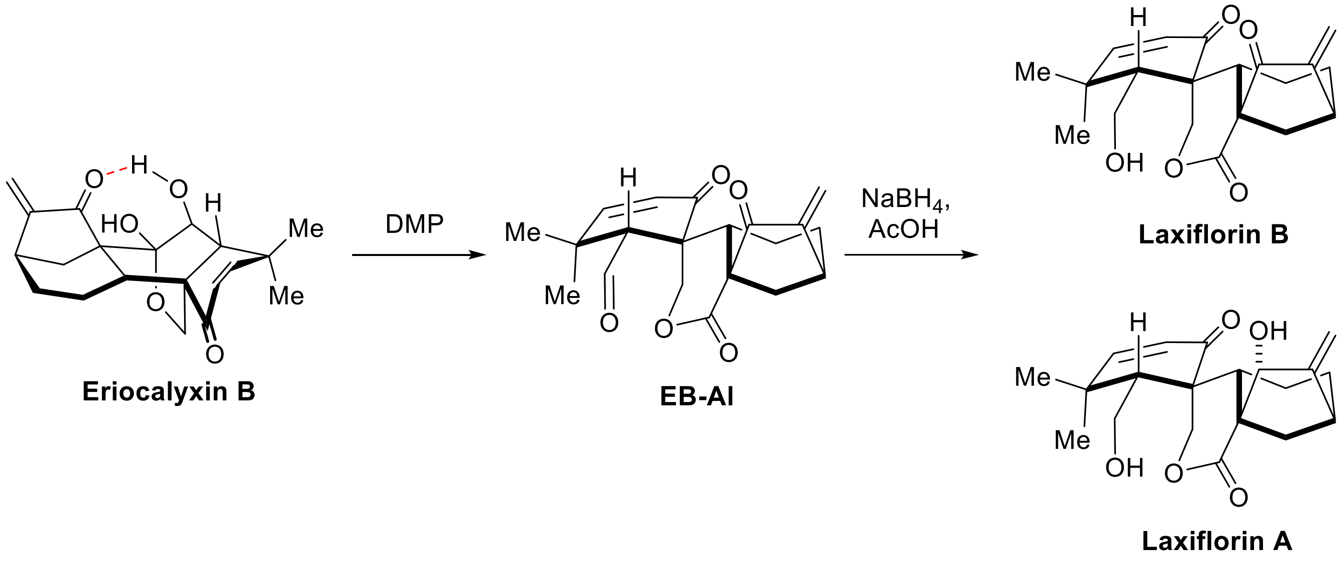


To a stirred solution of Eriocalyxin B (344 mg, 1 mmol) in DCM (10 mL) at room temperature was added Dess-Martin periodane (852 mg, 2 mmol). The resulting solution was warmed up the temperature to 40 ºC for another 0.5 h. After the reactant was disappeared which was monitored by TLC, the solution was cooled down to room temperature. Then the reaction was quenched by addition of saturated an aqueous $Na_2S_2O_3$ (2 mL) and saturated brine (10 mL). The aqueous phase was extracted by diethyl ether (10 mL×3) and the combined organic extracts were washed with brine (10 mL), dried over $Na_2SO_4$, filtered and concentrated under reduced pressure. Silica gel flash column chromatography (silica gel, Hexanes/Ethyl acetate = 1:1) of the residue gave a white solid (257 mg, 0.75 mmol). To a solution of the white product from last step in THF (10 mL) and AcOH (0.4 mL) was added sodium borohydride (30 mg，0.75 mmol) at 0 ℃. Then the solution was warmed up to room temperature and kept stirring for another 1 h. After the reactant was disappeared monitored by TLC, the reaction was quenched by addition of saturated an aqueous $NaHCO_3$ (2 mL) and saturated brine (10 mL). The aqueous phase was extracted by ethyl acetate (10 mL×3) and the combined organic extracts were washed with brine, dried over $Na_2SO_4$, filtered and concentrated under reduced pressure. Silica gel flash column chromatography (silica gel, Hexanes/Ethyl acetate = 1:1) of the residue gave Laxiflorin B (LB) as a white solid (241 mg, 0.70 mmol, totally yield 70%) with trace Laxiflorin A (LA) (18 mg, 0.05 mmol，totally yield 5%). The spectra data of LB was matched with the reported literature[1-3].

To a stirred solution of Eriocalyxin B (344 mg, 1 mmol) in DCM (10 mL) at room temperature was added Dess-Martin periodane (852 mg, 2 mmol). The resulting solution was warmed up the temperature to 40 ºC for another 0.5 h. After the reactant was disappeared which was monitored by TLC, the solution was cooled down to room temperature. Then the reaction was quenched by addition of saturated an aqueous $Na_2S_2O_3$ (2 mL) and saturated brine (10 mL). The aqueous phase was extracted by diethyl ether (10 mL×3) and the combined organic extracts were washed with brine (10 mL), dried over $Na_2SO_4$, filtered and concentrated under reduced pressure. Silica gel flash column chromatography (silica gel, Hexanes/Ethyl acetate = 1:1) of the residue gave a white solid (257 mg, 0.75 mmol). To a solution of the white product from last step in THF (20 mL) and AcOH (1.0 mL) was added sodium borohydride (80 mg，2.0 mmol) in 3 portions at 0 ℃. Then the solution was warmed up to room temperature and kept stirring for another 1 h. After the reactant was disappeared monitored by TLC, the reaction was quenched by addition of saturated an aqueous $NaHCO_3$ (2 mL) and saturated brine

(10 mL). The aqueous phase was extracted by ethyl acetate (10 mL×3) and the combined organic extracts were washed with brine, dried over $Na_2SO_4$, filtered and concentrated under reduced pressure. Silica gel flash column chromatography (silica gel, Hexanes/Ethyl acetate = 1:1) of the residue gave Laxiflorin A (LA) as a white solid (173 mg, 0.50 mmol, totally yield 50%) **LA**: $^1$H NMR (500 MHz, Chloroform-*d*) δ 6.56 (d, *J* = 10.1 Hz, 1H), 5.84 (d, *J* = 10.2 Hz, 1H), 5.14 (s, 1H), 5.10 (s, 1H), 4.90 (d, *J* = 10.9 Hz, 1H), 4.52 (s, 2H), 4.46 (d, *J* = 10.9 Hz, 1H), 4.00 (dd, *J* = 12.3, 3.8 Hz, 1H), 3.92 (dd, *J* = 12.3, 3.8 Hz, 1H), 3.43 (s, 1H), 2.72 (dd, *J* = 8.3, 5.3 Hz, 1H), 2.66 (dd, *J* = 12.7, 4.1 Hz, 1H), 2.27 (d, *J* = 12.4 Hz, 1H), 2.15 – 2.01 (m, 3H), 2.00 (s, 1H), 1.55 – 1.44 (m, 1H), 1.40 (tt, *J* = 12.8, 7.5 Hz, 2H), 1.23 (m, 6H). $^{13}$C NMR (125 MHz, $CDCl_3$) δ 200.8, 176.0, 158.9, 158.3, 124.8, 109.3, 82.2, 69.7, 59.0, 52.4, 51.1, 47.3, 36.4, 36.2, 35.4, 32.8, 31.9, 30.6, 24.3, 17.1. HRMS (ESI/[M+Na]$^+$) calcd. for $C_{20}H_{26}NaO_5$: 369.1678, found 369.1675.

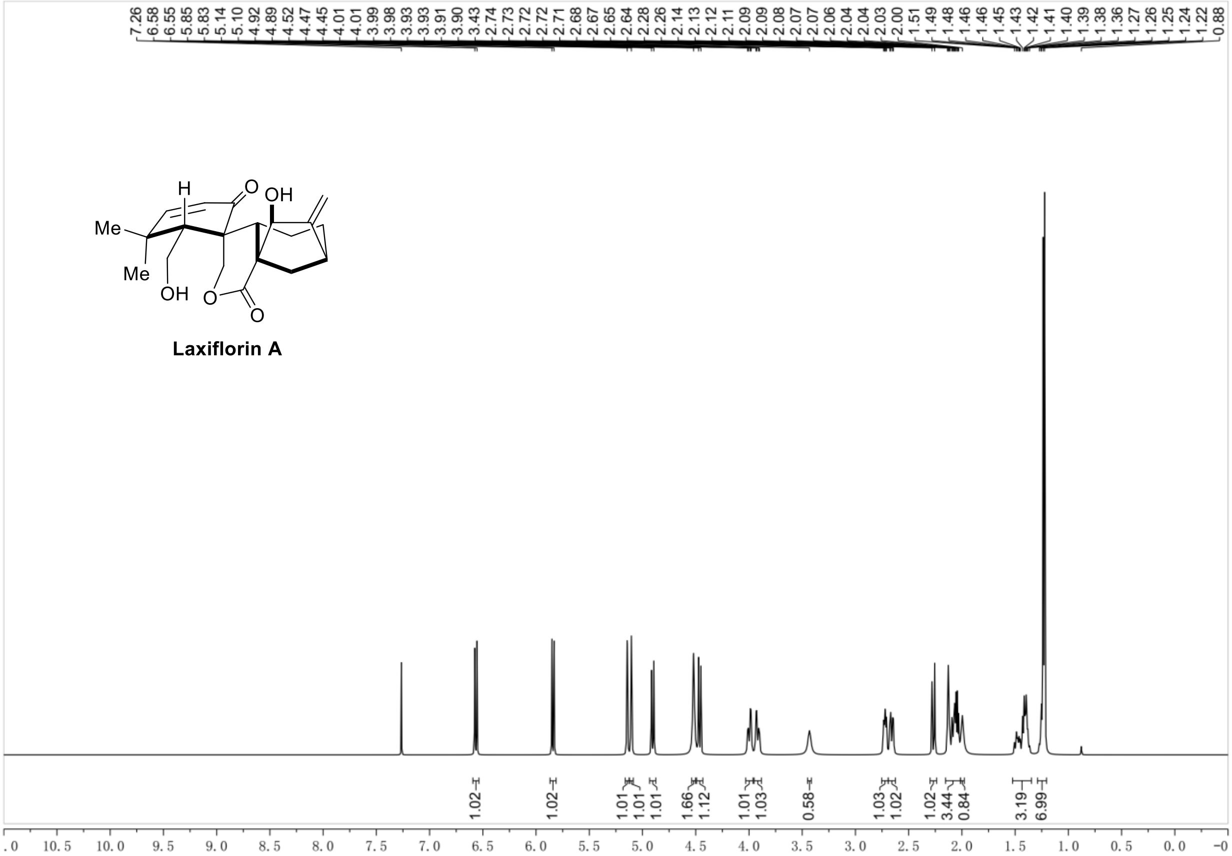
Laxiflorin A
Me
Me
OH
H
O
OH
O
O
7.26 6.58 6.55 5.85 5.83 5.14 5.10 4.92 4.89 4.52 4.47 4.45 4.01 4.01 3.99 3.98 3.93 3.93 3.91 3.90 3.43 2.74 2.73 2.72 2.72 2.71 2.68 2.67 2.65 2.64 2.28 2.26 2.14 2.13 2.12 2.11 2.09 2.09 2.08 2.07 2.07 2.06 2.04 2.04 2.03 2.00 1.51 1.49 1.48 1.46 1.46 1.45 1.43 1.42 1.41 1.40 1.39 1.38 1.36 1.27 1.26 1.25 1.24 1.22 0.88
1.02 1.02 1.01 1.01 1.01 1.66 1.12 1.01 1.03 0.58 1.03 1.02 1.02 3.44 0.84 3.19 6.99
10.5 10.0 9.5 9.0 8.5 8.0 7.5 7.0 6.5 6.0 5.5 5.0 4.5 4.0 3.5 3.0 2.5 2.0 1.5 1.0 0.5 0.0

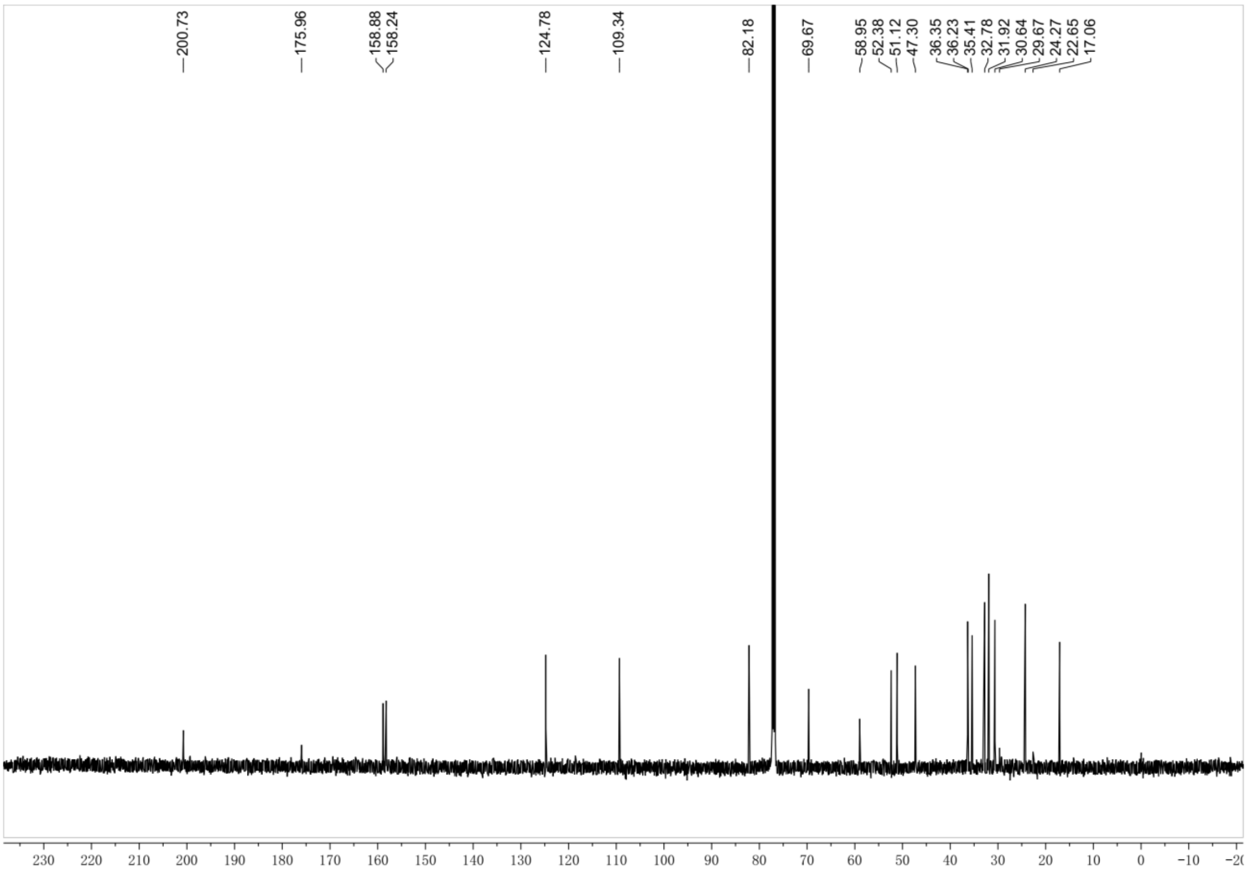
200.73 175.96 158.88 158.24 124.78 109.34 82.18 69.67 58.95 52.38 51.12 47.30 36.35 36.23 35.41 32.78 31.92 30.64 29.67 24.27 22.65 17.06
230 220 210 200 190 180 170 160 150 140 130 120 110 100 90 80 70 60 50 40 30 20 10 0 −10

**Synthesis of Laxiflorin B-Alanine**

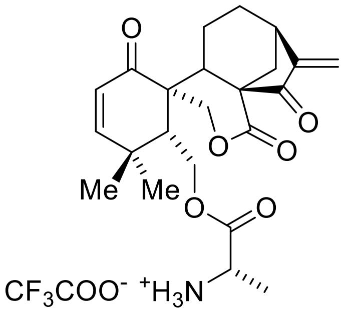


**LB-Ala**

To a stirred solution of Laxiflorin B (17 mg, 0.05 mmol) in DCM (1 mL) at room temperature was added Boc-L-alanine (9.5 mg, 0.05 mmol), EDCI (9.6 mg, 0.05 mmol) and DMAP (0.6 mg, 0.005 mmol). The resulting solution was stirred another 12 h until LB was disappeared monitored by TLC. The reaction was quenched by addition of saturated an aqueous $NaHCO_3$ (2 mL), followed by saturated brine (10 mL) and ethyl acetate (10 mL). The aqueous phase was extracted by ethyl acetate (10 mL×3) and the combined organic extracts were washed with brine, dried over $Na_2SO_4$, filtered and concentrated under reduced pressure. Silica gel flash column chromatography (silica gel, Hexanes/Ethyl acetate = 1:2) of the residue gave a white solid. To a solution of white solid from last step in DCM (1 mL) was added TFA (0.5 mL) at room temperature. Keep the solution stirring until the reactant was disappeared monitored by TLC, then remove the solvent under reduce pressure. To the resulting crude solid was added diethyl ether (10 mL), and the mixture was stirred for 15 min followed by filtered to remove the solvent. This wash operation was carried out for another twice, and the desired product was given LB-Ala as a white solid. **Laxiflorin B-Ala:** (24 mg, 0.045 mmol, 90%). $^1$H NMR (300 MHz, Methanol-$d_4$) δ 6.80 (d, $J$ = 10.2 Hz, 1H), 6.01 (s, 1H), 5.90 (d, $J$ = 10.2 Hz, 1H), 5.62 (s, 1H), 4.74 (d, $J$ =

11.3 Hz, 1H), 4.67 – 4.45 (m, 3H), 4.08 (q, *J* = 7.3 Hz, 1H), 3.16 (dd, *J* = 9.3, 4.8 Hz, 1H), 2.71 (d, *J* = 12.5 Hz, 1H), 2.48 (t, *J* = 4.2 Hz, 1H), 2.38 (dd, *J* = 12.3, 4.6 Hz, 1H), 2.29 (dd, *J* = 12.4, 5.2 Hz, 2H), 1.78 – 1.40 (m, 5H), 1.32 – 1.12 (m, 7H).$^{13}$C NMR (75 MHz, MeOD) δ 202.5, 199.1, 170.3, 169.2, 158.6, 150.9, 143.0, 123.5, 118.2, 90.1, 69.3, 62.3, 58.1, 51.1, 44.1, 41.9, 36.0, 34.7, 30.1, 29.7, 29.3, 24.3, 22.5, 17.3, 14.4.HRMS (ESI/$[M+H]^+$) calcd. for $C_{23}H_{30}NO_6$: 416.2068, found 416.2074.

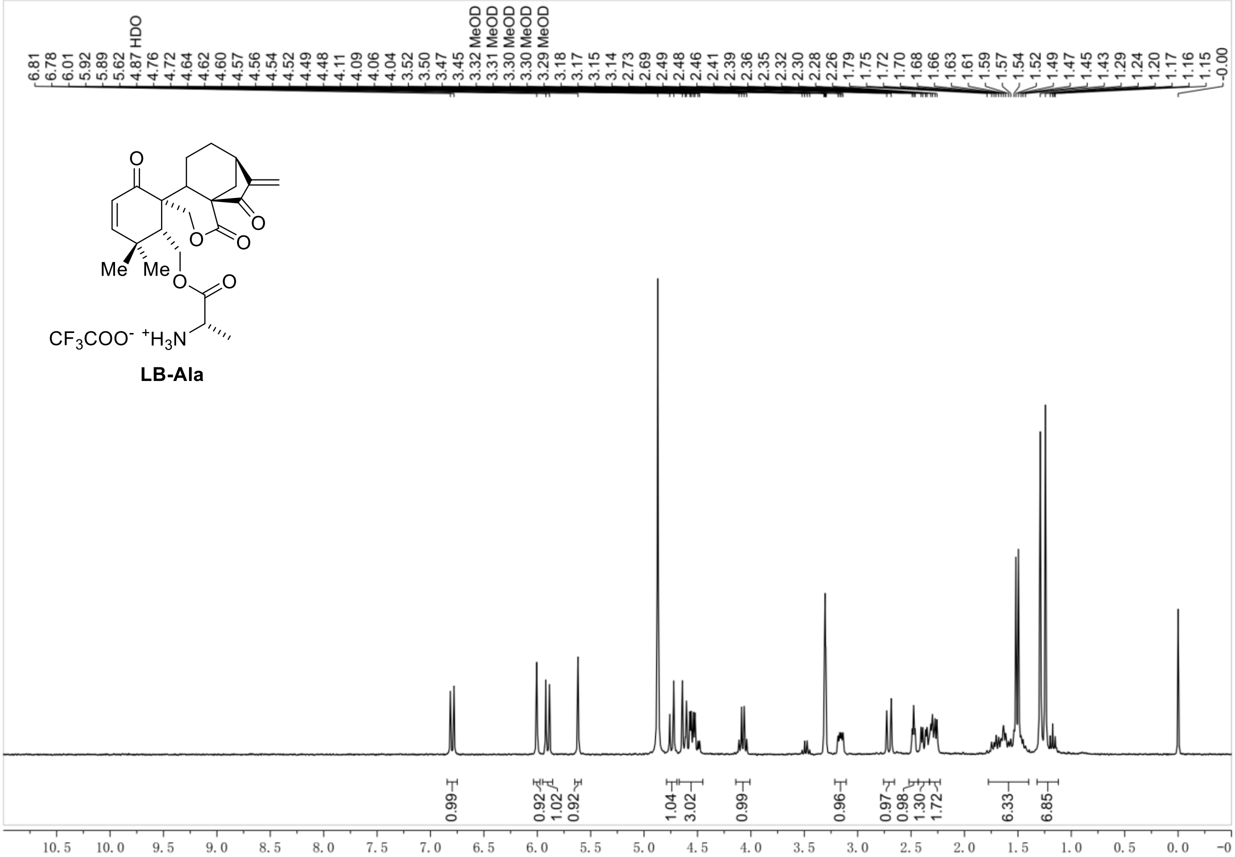
6.81
6.78
6.01
5.92
5.89
5.62
4.87 HDO
4.76
4.72
4.64
4.62
4.60
4.57
4.56
4.54
4.52
4.49
4.48
4.11
4.09
4.06
4.04
3.52
3.50
3.47
3.45
3.32 MeOD
3.31 MeOD
3.30 MeOD
3.30 MeOD
3.29 MeOD
3.18
3.17
3.15
3.14
2.73
2.69
2.49
2.48
2.46
2.41
2.39
2.36
2.35
2.32
2.30
2.28
2.26
1.79
1.75
1.72
1.70
1.68
1.66
1.63
1.61
1.59
1.57
1.54
1.52
1.49
1.47
1.45
1.43
1.29
1.24
1.20
1.17
1.16
1.15
-0.00
O
O
O
O
Me
Me
O
O
$CF_3COO^-$ $^+H_3N$
LB-Ala
0.99
0.92
1.02
0.92
1.04
3.02
0.99
0.96
0.97
0.98
1.30
1.72
6.33
6.85
10.5
10.0
9.5
9.0
8.5
8.0
7.5
7.0
6.5
6.0
5.5
5.0
4.5
4.0
3.5
3.0
2.5
2.0
1.5
1.0
0.5
0.0

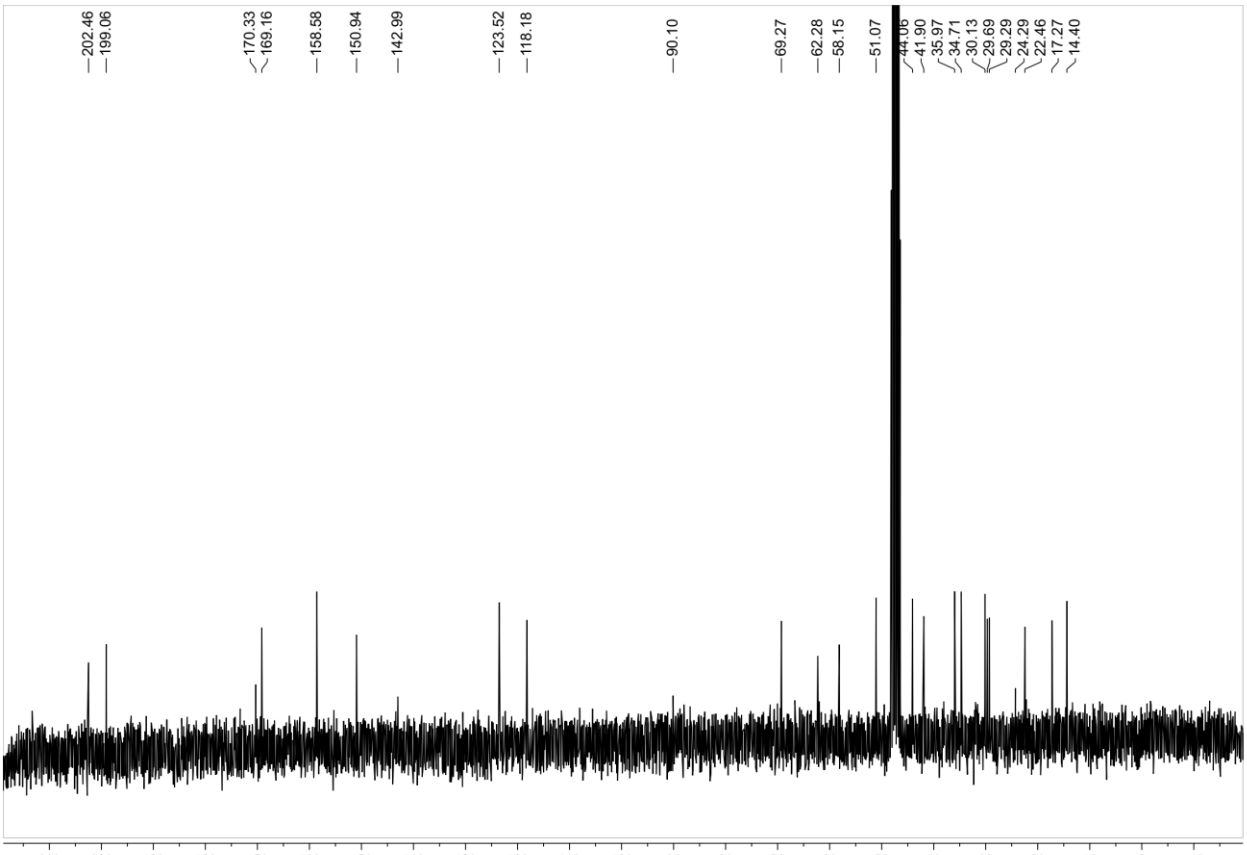
202.46
199.06
170.33
169.16
158.58
150.94
142.99
123.52
118.18
90.10
69.27
62.28
58.15
51.07
44.06
41.90
35.97
34.71
30.13
29.69
29.29
24.29
22.46
17.27
14.40
210
200
190
180
170
160
150
140
130
120
110
100
90
80
70
60
50
40
30
20
10
0
-10

**Synthesis of Laxiflorin -biotin analogues**

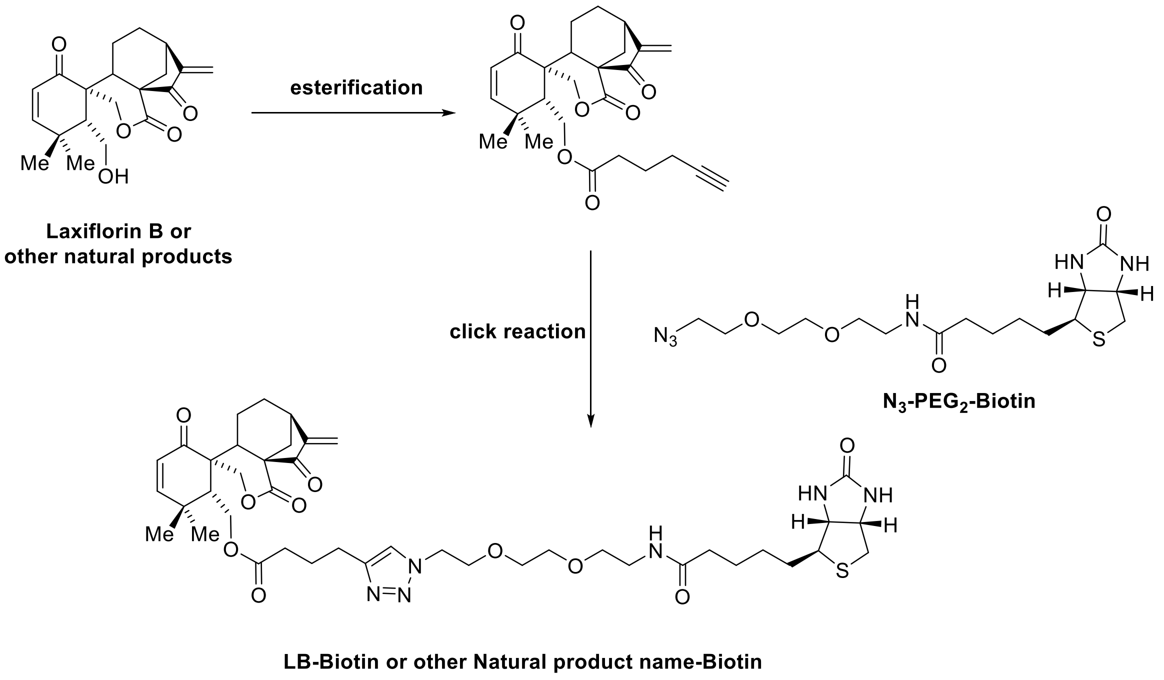


To a stirred solution of Laxiflorin B (17 mg, 0.05 mmol) (or other natural product) in DCM (1 mL) at room temperature was added 5-Hexynoic acid (5.5 mg, 0.05 mmol), EDCI (9.6 mg, 0.05 mmol) and DMAP (0.6 mg, 0.005 mmol). The resulting solution was stirred another 12 h until LB was disappeared monitored by TLC. The reaction was quenched by addition of saturated an aqueous $NaHCO_3$ (2 mL), followed by saturated brine (10 mL) and ethyl acetate (10 mL). The aqueous phase was extracted by ethyl acetate (10 mL×3) and the combined organic extracts were washed with brine, dried over $Na_2SO_4$, filtered and concentrated under reduced pressure. Silica gel flash column chromatography (silica gel, Hexanes/Ethyl acetate = 1:1) of the residue gave a desired product. To a solution of last step product in THF (2 mL) and $H_2O$ (1 mL) was added known compound $N_3$-$PEG_2$-Biotin (0.05 mmol) followed by sodium carboxymethyl cellulose (0.005 mmol). The resulting mixture was degased and added copper sulfate pentahydrate (1.25 mg, 0.005 mmol) under the argon atmosphere conditions. Keep the solution stirring another 12 h until the reactant was disappeared which was monitored by TLC. The reaction was quenched by addition of saturated an aqueous $NaHCO_3$ (2 mL), followed by ethyl acetate (10 mL). The aqueous phase was extracted by ethyl acetate (10 mL×3) and the combined organic extracts were washed with brine, dried over $Na_2SO_4$, filtered and concentrated under reduced pressure. Silica gel flash column chromatography (silica gel, MeOH/DCM = 1:10) of the residue gave a desired product.

**Laxiflorin B-biotin:** (35 mg, 0.042 mmol, 84%). $^1$H NMR (500 MHz, Methanol-$d_4$) δ 7.92 (t, *J* = 5.6 Hz, 1H), 7.85 (s, 1H), 6.78 (d, *J* = 10.2 Hz, 1H), 5.96 (s, 1H), 5.88 (d, *J* = 10.2 Hz, 1H), 5.59 (s, 1H), 4.71 (d, *J* = 11.3 Hz, 1H), 4.65 (d, *J* = 11.3 Hz, 1H), 4.56 (t, *J* = 5.1 Hz, 2H), 4.49 (dd, *J* = 7.9, 4.8 Hz, 1H), 4.41 (dd, *J* = 13.0, 3.8 Hz, 1H), 4.38 – 4.27 (m, 2H), 3.89 (t, *J* = 5.1 Hz, 2H), 3.66 – 3.59 (m, 2H), 3.59 – 3.54 (m, 2H), 3.50 (t, *J* = 5.5 Hz, 2H), 3.39 – 3.31 (m, 3H), 3.20 (ddd, *J* = 9.0, 5.8, 4.4 Hz, 1H), 3.14 (dd, *J* = 9.4, 4.8 Hz, 1H), 2.92 (dd, *J* = 12.7, 5.0 Hz, 1H), 2.81 – 2.64 (m, 4H), 2.42 (t, *J* = 4.3 Hz, 1H), 2.36 (dt, *J* = 10.1, 6.1 Hz, 3H), 2.28 (td, *J* = 14.8, 13.6, 5.9 Hz, 2H), 2.21 (t, *J* = 7.4 Hz, 2H), 2.05 – 1.91 (m, 2H), 1.79 – 1.54 (m, 6H), 1.51 – 1.37 (m, 3H), 1.28 (s, 3H), 1.23 (s, 3H). $^{13}$C NMR (125 MHz, MeOD) δ 202.4, 199.8, 174.7, 172.6, 170.7, 164.6, 159.1, 151.2, 146.6, 123.7, 122.8, 118.5, 70.0, 69.8, 69.8, 69.2, 69.0, 62.0, 60.3, 60.2, 58.5, 55.6, 51.5, 49.9, 44.3, 42.0, 39.7, 38.9, 36.3, 35.3, 35.0, 33.0, 30.5, 30.2, 29.4, 28.4, 28.1, 25.4, 24.3, 24.2, 22.5, 17.6. HRMS (ESI/$[M+Na]^+$) calcd. for $C_{42}H_{58}N_6NaO_{10}S$: 861.3827, found 861.3830.

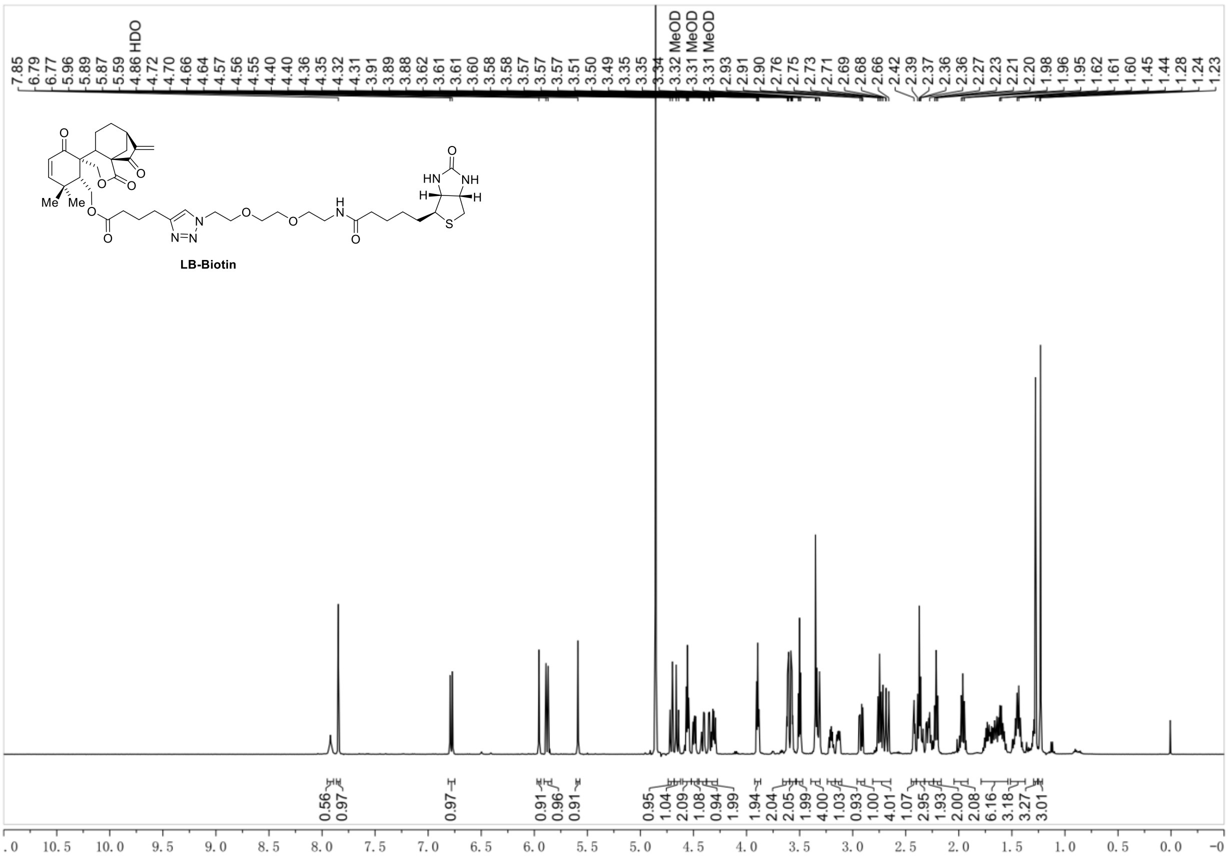
7.85
6.79
6.77
5.96
5.89
5.87
5.59
4.86 HDO
4.72
4.70
4.66
4.64
4.57
4.56
4.55
4.40
4.40
4.36
4.35
4.32
4.31
3.91
3.89
3.88
3.62
3.61
3.60
3.58
3.58
3.57
3.57
3.57
3.51
3.50
3.49
3.35
3.35
3.34
3.32 MeOD
3.31 MeOD
3.31 MeOD
2.93
2.91
2.90
2.76
2.75
2.73
2.71
2.69
2.68
2.66
2.42
2.39
2.37
2.36
2.36
2.27
2.23
2.21
2.20
1.98
1.96
1.95
1.62
1.61
1.60
1.45
1.44
1.28
1.24
1.23
LB-Biotin
0.56
0.97
0.97
0.91
0.96
0.91
0.95
1.04
2.09
1.08
0.94
1.99
1.94
2.04
2.05
1.99
4.00
1.03
0.93
1.00
4.01
1.07
2.95
1.93
2.00
2.08
6.16
3.18
3.27
3.01
10.5 10.0 9.5 9.0 8.5 8.0 7.5 7.0 6.5 6.0 5.5 5.0 4.5 4.0 3.5 3.0 2.5 2.0 1.5 1.0 0.5 0.0

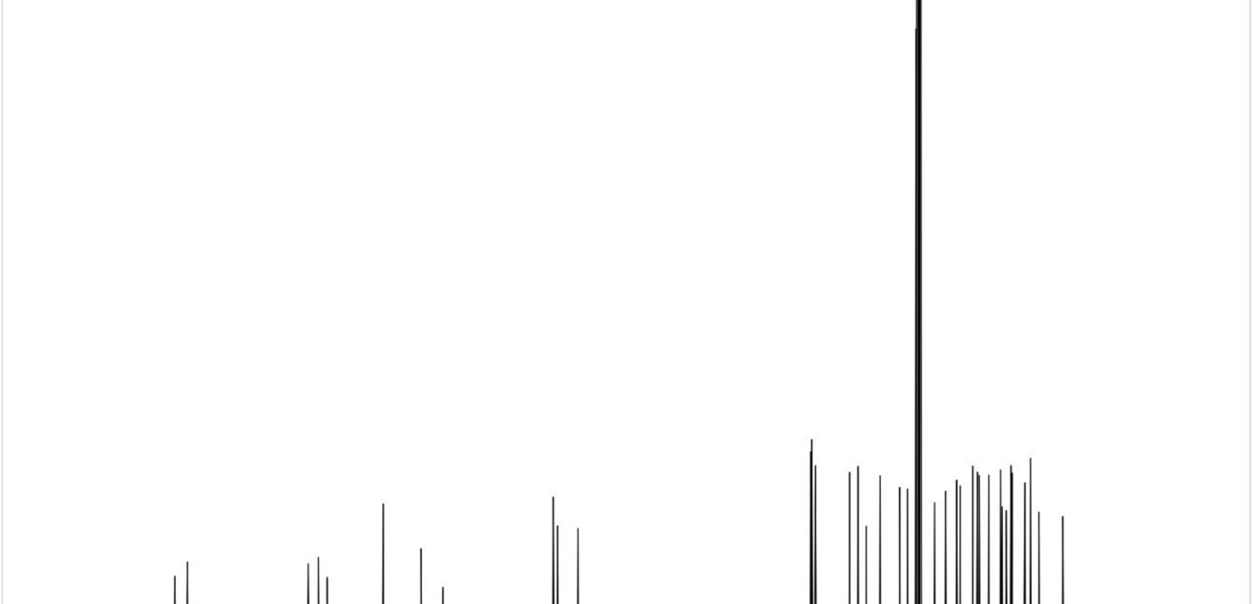
202.44
199.84
174.67
172.56
170.75
164.63
159.05
151.18
146.62
123.66
122.76
118.52
70.04
69.84
69.80
69.20
69.04
61.96
60.27
60.21
58.49
55.59
51.54
49.91
44.26
41.97
39.67
38.93
36.31
35.35
34.96
32.96
30.52
30.18
29.36
28.35
28.10
25.44
24.28
24.21
22.53
17.57
230 220 210 200 190 180 170 160 150 140 130 120 110 100 90 80 70 60 50 40 30 20 10 0 −10

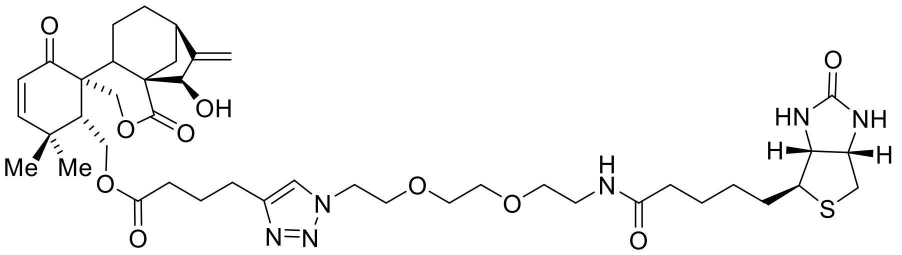

**LA-Biotin**

**LA-Biotin**: 33 mg, 0.04 mmol, totally yield 80%.$^{1}$H NMR (500 MHz, Methanol-$d_4$) δ 7.83 (s, 1H), 6.75 (d, *J* = 10.2 Hz, 1H), 5.85 (d, *J* = 10.2 Hz, 1H), 5.07 (dt, *J* = 10.7, 1.8 Hz, 2H), 4.85 (s, 2H), 4.68 – 4.53 (m, 3H), 4.52 – 4.43 (m, 2H), 4.44 – 4.33 (m, 2H), 4.31 (dd, *J* = 7.9, 4.5 Hz, 1H), 3.90 (t, *J* = 5.1 Hz, 2H), 3.70 – 3.54 (m, 4H), 3.51 (t, *J* = 5.6 Hz, 2H), 3.36 – 3.28 (m, 7H), 3.21 (ddd, *J* = 8.9, 5.8, 4.4 Hz, 1H), 2.93 (dd, *J* = 12.7, 4.9 Hz, 1H), 2.75 (t, *J* = 7.6 Hz, 2H), 2.70 (dd, *J* = 11.7, 6.8 Hz, 3H), 2.45 (t, *J* = 4.2 Hz, 1H), 2.41 (t, *J* = 7.5 Hz, 2H), 2.22 (t, *J* = 7.4 Hz, 2H), 2.09 – 1.87 (m, 2H), 1.80 – 1.55 (m, 4H), 1.44 (t, *J* = 7.7 Hz, 4H), 1.27 (s, 3H), 1.21 (s, 3H). $^{13}$C NMR (125 MHz, MeOD) δ 201.1, 176.8, 174.7, 172.9, 164.6, 159.2, 158.8, 146.8, 123.9, 122.8, 108.4, 81.6, 70.0, 69.8, 69.4, 69.2, 69.0, 62.0, 60.6, 60.2, 55.6, 52.3, 51.0, 49.9, 44.5, 39.7, 38.9, 36.1, 36.0, 35.3, 34.8, 33.1, 32.8, 30.6, 30.1, 28.3, 28.1, 25.4, 24.3, 24.2, 22.6, 16.4. HRMS (ESI/[M+Na]$^+$) calcd. for $C_{42}H_{60}N_6NaO_{10}S$: 863.3984, found 863.3987.

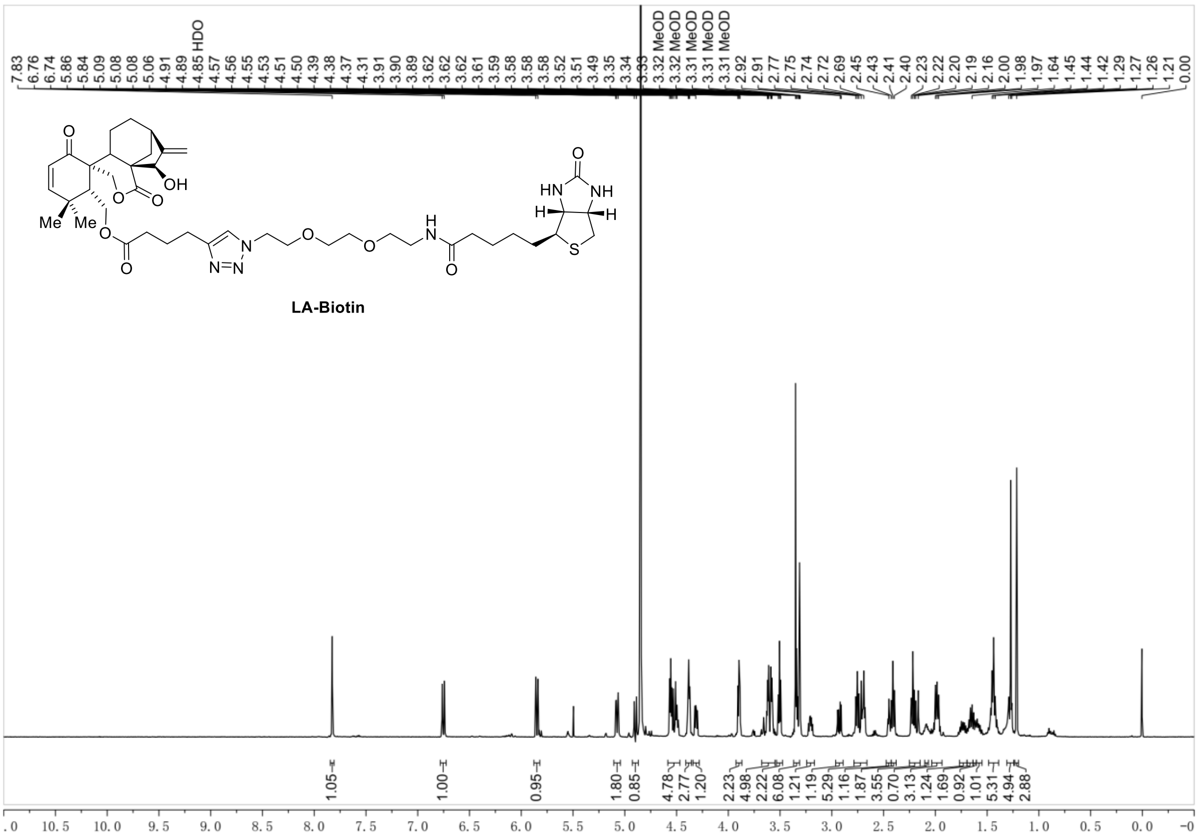

7.83
6.76
6.74
5.86
5.84
5.09
5.08
5.08
5.06
4.91
4.89
4.85 HDO
4.57
4.56
4.55
4.53
4.51
4.50
4.39
4.38
4.37
4.31
3.91
3.90
3.89
3.62
3.62
3.62
3.61
3.59
3.58
3.58
3.58
3.52
3.51
3.49
3.35
3.34
3.33
3.32 MeOD
3.32 MeOD
3.31 MeOD
3.31 MeOD
3.31 MeOD
2.92
2.91
2.77
2.75
2.74
2.72
2.69
2.45
2.43
2.41
2.40
2.23
2.22
2.20
2.19
2.16
2.00
1.98
1.97
1.64
1.45
1.44
1.42
1.29
1.27
1.26
1.21
0.00
LA-Biotin
1.05
1.00
0.95
1.80
0.85
4.78
2.77
1.20
2.23
4.98
2.22
6.08
1.21
1.19
5.29
1.16
1.87
3.55
0.70
3.13
1.24
1.69
0.92
1.01
5.31
4.94
2.88
10.5 10.0 9.5 9.0 8.5 8.0 7.5 7.0 6.5 6.0 5.5 5.0 4.5 4.0 3.5 3.0 2.5 2.0 1.5 1.0 0.5 0.0


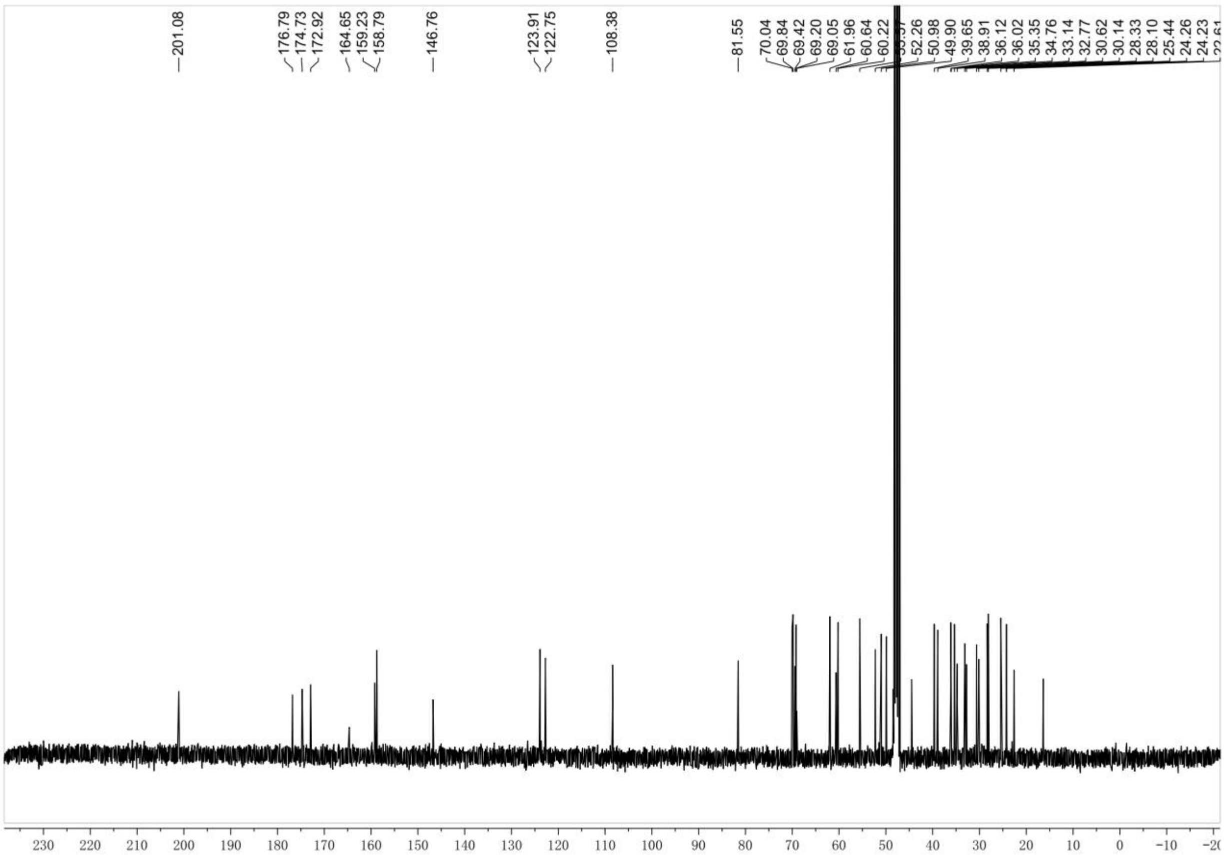

201.08
176.79
174.73
172.92
164.65
159.23
158.79
146.76
123.91
122.75
108.38
81.55
70.04
69.84
69.42
69.20
69.05
61.96
60.64
60.22
52.26
50.98
49.90
39.65
38.91
36.12
36.02
35.35
34.76
33.14
32.77
30.62
30.14
28.33
28.10
25.44
24.26
24.23
230 220 210 200 190 180 170 160 150 140 130 120 110 100 90 80 70 60 50 40 30 20 10 0 −10

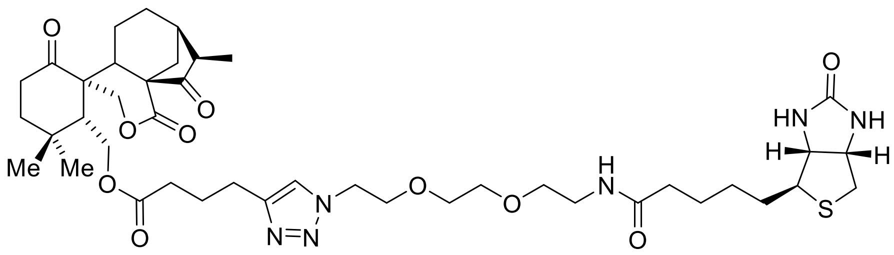

**LB-Di-Biotin**

**LB-Di-Biotin**:32 mg, 0.038 mmol, totally yield 76%.$^1$H NMR (400 MHz, DMSO-$d_6$) δ 7.81 (s, 1H), 7.77 (t, *J* = 5.6 Hz, 1H), 6.37 (s, 1H), 6.32 (s, 1H), 4.71 (d, *J* = 11.6 Hz, 1H), 4.45 (t, *J* = 5.3 Hz, 2H), 4.33 – 4.25 (m, 2H), 4.24 (dd, *J* = 12.6, 3.8 Hz, 1H), 4.19 – 4.07 (m, 2H), 3.78 (t, *J* = 5.3 Hz, 2H), 3.50 (dd, *J* = 6.2, 3.6 Hz, 3H), 3.46 (dd, *J* = 6.0, 3.5 Hz, 2H), 3.35 (t, *J* = 5.9 Hz, 2H), 3.15 (q, *J* = 5.8 Hz, 2H), 3.12 – 3.03 (m, 1H), 2.80 (dd, *J* = 12.4, 5.0 Hz, 1H), 2.74 – 2.50 (m, 5H), 2.32 (t, *J* = 7.5 Hz, 2H), 2.22 (ddd, *J* = 21.4, 11.7, 5.1 Hz, 2H), 2.09 – 1.90 (m, 3H), 1.90 – 1.78 (m, 2H), 1.74 (dt, *J* = 29.5, 7.5 Hz, 1H), 1.62 – 1.52 (m, 1H), 1.51 – 1.35 (m, 3H), 1.29 (s, 1H), 1.23 (s, 2H), 1.08 – 0.99 (m, 9H). $^{13}$C NMR (100 MHz, DMSO) δ 217.1, 212.1, 172.8, 172.7, 169.8, 163.3, 146.5, 122.9, 70.1, 70.0, 69.7, 69.4, 68.9, 61.9, 61.6, 59.8, 58.9, 56.0, 53.7, 49.8, 48.3, 47.6, 43.1, 39.0, 37.1, 36.7, 35.7, 33.6, 32.2, 31.5, 28.8, 28.6, 25.8, 24.9, 24.8, 24.2, 20.1, 17.9, 12.0. HRMS (ESI/[M+Na]$^+$) calcd. for $C_{42}H_{62}N_6NaO_{10}S$: 865.4140, found 865.4145.

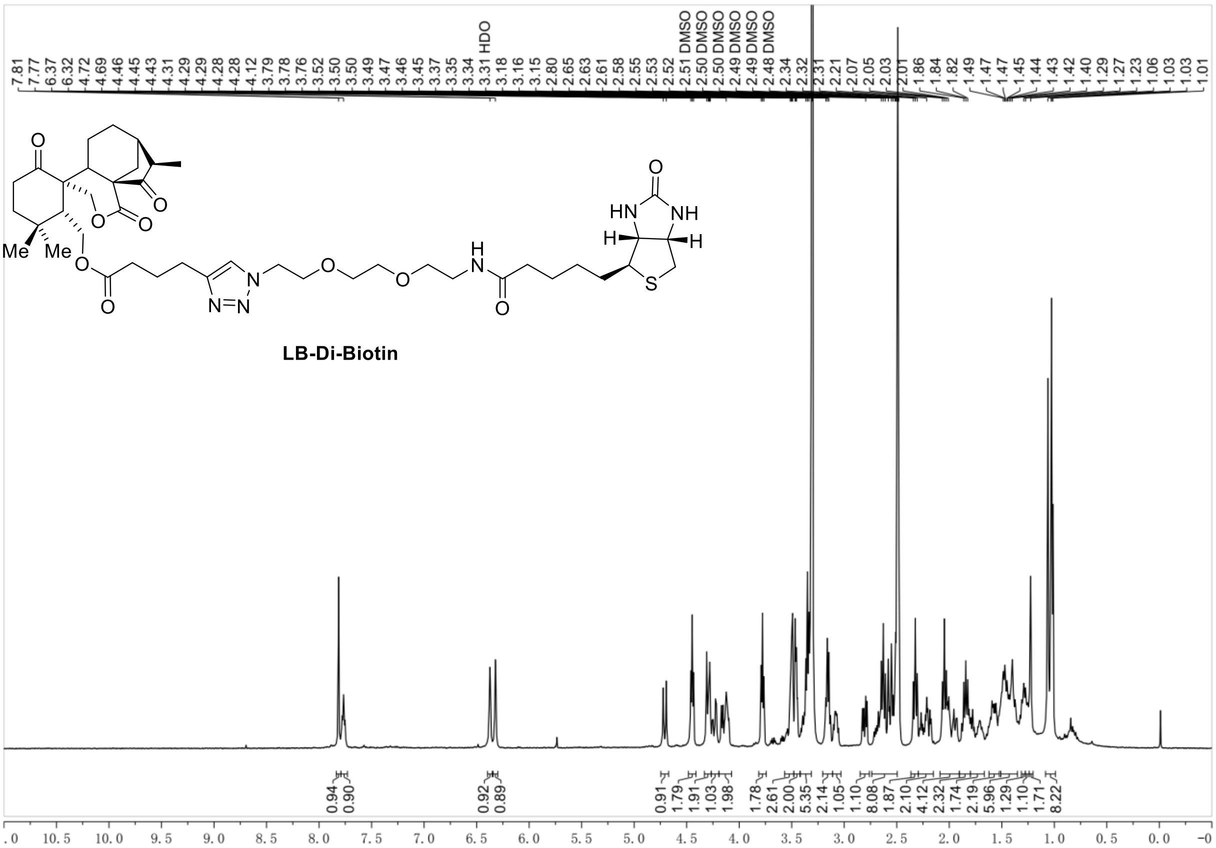
7.81
7.77
6.37
6.32
4.72
4.69
4.46
4.45
4.43
4.31
4.29
4.29
4.28
4.28
4.12
3.79
3.78
3.76
3.52
3.50
3.50
3.49
3.47
3.46
3.45
3.37
3.35
3.34
3.31 HDO
3.18
3.16
3.15
2.80
2.65
2.63
2.61
2.58
2.55
2.53
2.52
2.51 DMSO
2.50 DMSO
2.50 DMSO
2.49 DMSO
2.49 DMSO
2.48 DMSO
2.34
2.32
2.31
2.21
2.07
2.05
2.03
2.01
1.86
1.84
1.82
1.49
1.47
1.47
1.45
1.44
1.43
1.42
1.40
1.29
1.27
1.23
1.06
1.03
1.03
1.01
Me
Me
HN
NH
H
H
S
N
N=N
O
LB-Di-Biotin
0.94
0.90
0.92
0.89
0.91
1.79
1.91
1.03
1.98
1.78
2.61
2.00
5.35
2.14
1.05
1.10
8.08
1.87
2.10
4.12
2.32
1.74
2.19
5.96
1.29
1.10
1.71
8.22
10.5 10.0 9.5 9.0 8.5 8.0 7.5 7.0 6.5 6.0 5.5 5.0 4.5 4.0 3.5 3.0 2.5 2.0 1.5 1.0 0.5 0.0

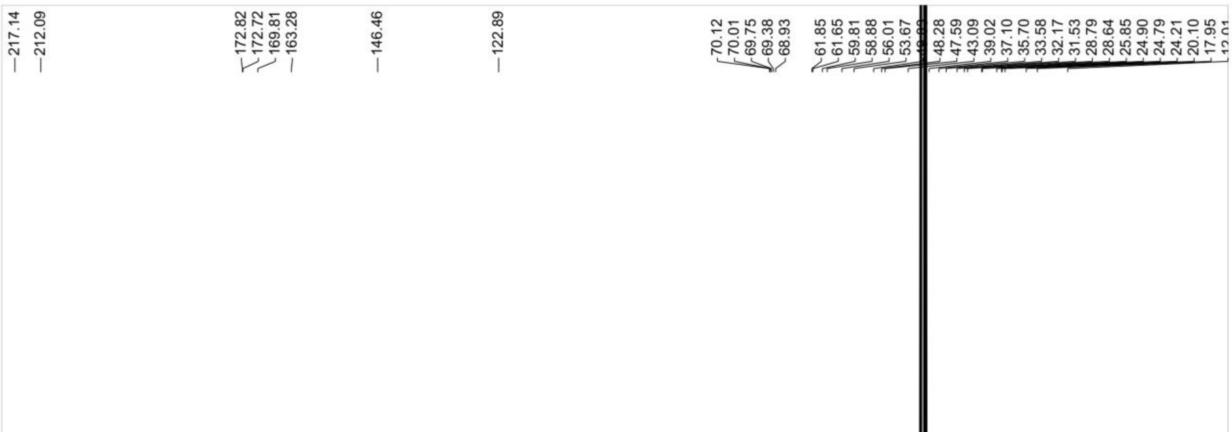
217.14
212.09
172.82
172.72
169.81
163.28
146.46
122.89
70.12
70.01
69.75
69.38
68.93
61.85
61.65
59.81
58.88
56.01
53.67
48.28
47.59
43.09
39.02
37.10
35.70
33.58
32.17
31.53
28.79
28.64
25.85
24.90
24.79
24.21
20.10
17.95
210 200 190 180 170 160 150 140 130 120 110 100 90 80 70 60 50 40 30 20 10 0 −10

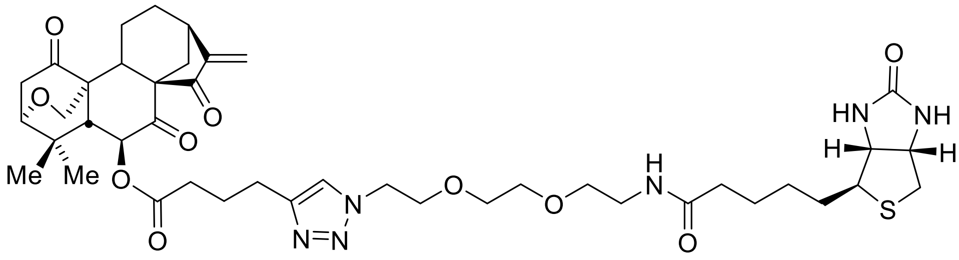

**LJ-Biotin**

**LJ-Biotin:**33 mg, 0.04 mmol, totally yield 80%.$^{1}$H NMR (400 MHz, Methanol-*d*$_4$) δ 7.88 (s, 1H), 6.04 (s, 1H), 5.71 (d, *J* = 12.6 Hz, 1H), 5.49 (s, 1H), 4.90 (s, 1H), 4.59 (t, *J* = 5.1 Hz, 2H), 4.52 (dd, *J* = 7.9, 4.8 Hz, 1H), 4.33 (dd, *J* = 7.9, 4.4 Hz, 1H), 4.19 (dd, *J* = 9.9, 1.7 Hz, 1H), 3.92 (t, *J* = 5.1 Hz, 2H), 3.83 – 3.71 (m, 1H), 3.75 – 3.54 (m, 5H), 3.53 (t, *J* = 5.5 Hz, 2H), 3.48 – 3.34 (m, 3H), 3.22 (ddt, *J* = 12.0, 6.2, 3.7 Hz, 2H), 2.95 (dd, *J* = 12.7, 4.9 Hz, 1H), 2.91 – 2.80 (m, 3H), 2.78 – 2.63 (m, 3H), 2.56 (td, *J* = 7.3, 4.4 Hz, 2H), 2.41 (d, *J* = 12.0 Hz, 1H), 2.24 (t, *J* = 7.3 Hz, 2H), 2.16 (dd, *J* = 12.7, 1.5 Hz, 1H), 2.13 – 2.02 (m, 2H), 1.94 (dd, *J* = 12.2, 4.9 Hz, 1H), 1.83 – 1.72 (m, 3H), 1.75 – 1.64 (m, 2H), 1.61 (dtd, *J* = 13.4, 9.0, 7.6, 4.7 Hz, 2H), 1.53 – 1.39 (m, 3H), 1.38 – 1.23 (s, 3H), 1.12 (s, 3H). $^{13}$C NMR (100 MHz, MeOD) δ 208.0, 201.6, 200.0, 174.8, 172.0, 164.7, 148.2, 123.0, 116.3, 77.0, 73.6, 70.1, 69.9, 69.3, 69.1, 62.0, 60.9, 60.3, 60.1, 55.7, 51.7, 50.0, 41.3, 39.7, 39.0, 38.8, 37.3, 36.8, 35.9, 35.4, 32.7, 30.8, 28.4, 28.2, 28.1, 25.5, 24.4, 24.1, 22.0, 20.3. HRMS (ESI/[M+Na]$^{+}$) calcd. for $C_{42}H_{58}N_6NaO_{10}S$: 861.3827, found 861.3832.

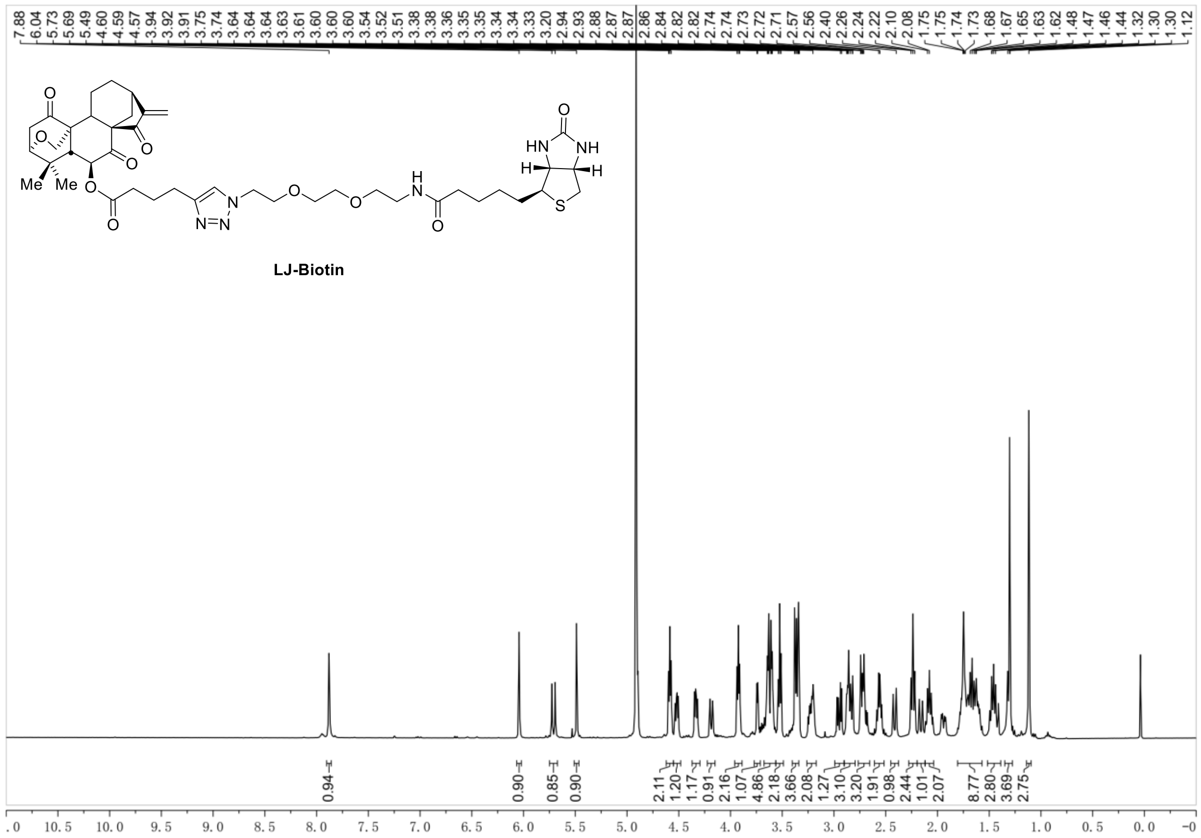
LJ-Biotin
7.88 6.04 5.73 5.69 5.49 4.60 4.59 4.57 3.94 3.92 3.91 3.75 3.74 3.64 3.64 3.63 3.61 3.60 3.60 3.54 3.52 3.51 3.38 3.38 3.36 3.35 3.35 3.34 3.34 3.33 3.20 2.94 2.93 2.88 2.87 2.87 2.86 2.84 2.82 2.82 2.74 2.74 2.73 2.72 2.71 2.57 2.56 2.40 2.26 2.24 2.22 2.10 2.08 1.75 1.75 1.74 1.73 1.68 1.67 1.65 1.63 1.62 1.48 1.47 1.46 1.44 1.32 1.30 1.30 1.12
0.94 0.90 0.85 0.90 2.11 1.20 1.17 0.91 2.16 1.07 4.86 2.18 3.66 2.08 1.27 3.10 3.20 1.91 0.98 2.44 1.01 2.07 8.77 2.80 3.69 2.75

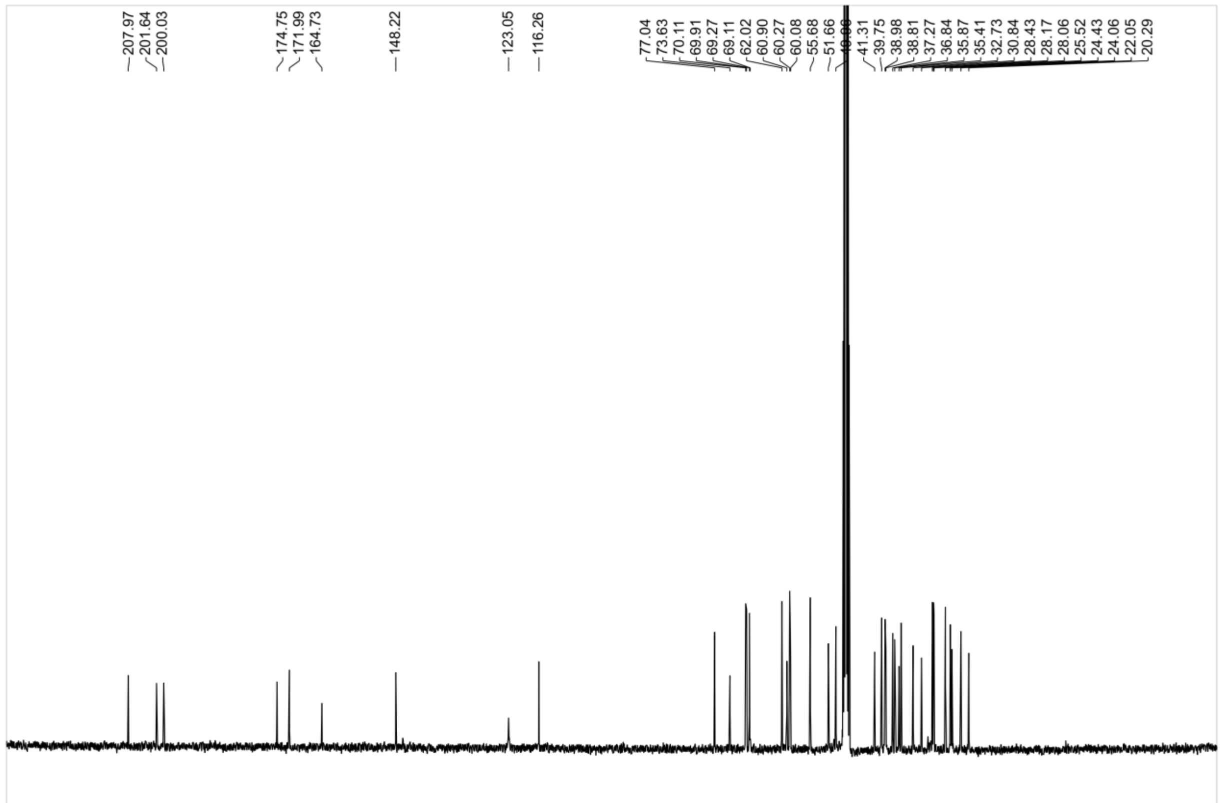
207.97 201.64 200.03 174.75 171.99 164.73 148.22 123.05 116.26 77.04 73.63 70.11 69.91 69.27 69.11 62.02 60.90 60.27 60.08 55.68 51.66 41.31 39.75 38.98 38.81 37.27 36.84 35.87 35.41 32.73 30.84 28.43 28.17 28.06 25.52 24.43 24.06 22.05 20.29

**Synthesis of Eriocalyxn B-biotin**

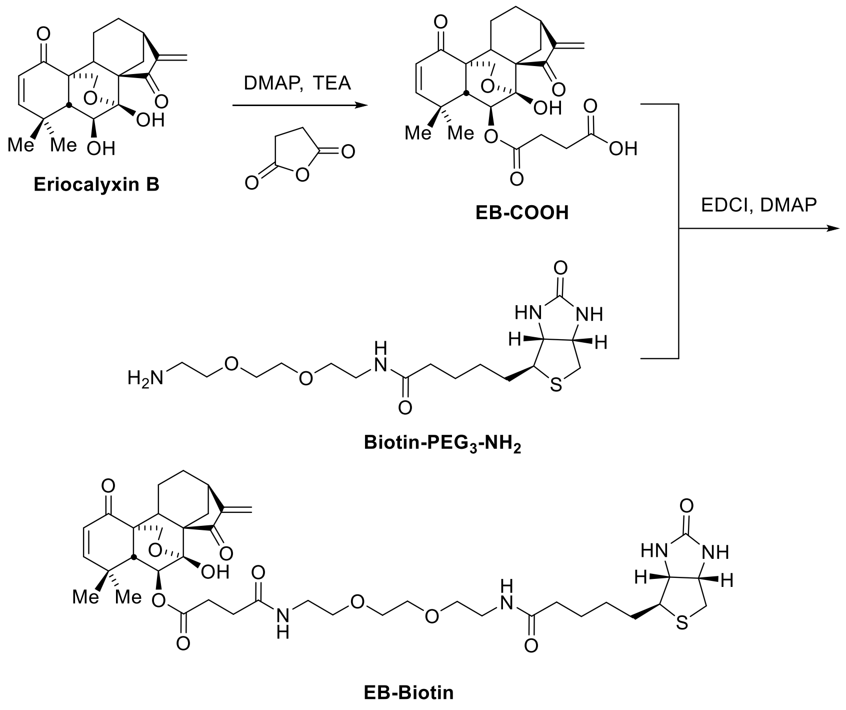


**EB-Biotin:** $^{1}$H NMR (400 MHz, Methanol-$d_4$) δ 6.89 (d, $J$ = 10.1 Hz, 1H), 5.99 – 5.88 (m, 1H), 5.85 (s, 1H), 5.56 – 5.44 (m, 1H), 5.38 (s, 1H), 4.53 (t, $J$ = 6.3 Hz, 2H), 4.34 (td, $J$ = 9.5, 8.9, 5.7 Hz, 3H), 4.14 – 4.03 (m, 2H), 3.77 – 3.66 (m, 4H), 3.58 (qd, $J$ = 6.5, 2.4 Hz, 8H), 3.40 (t, $J$ = 5.6 Hz, 6H), 3.23 (dt, $J$ = 13.2, 6.7 Hz, 2H), 2.96 (dd, $J$ = 12.8, 4.8 Hz, 2H), 2.82 (dd, $J$ = 14.1, 10.9 Hz, 4H), 2.73 (q, $J$ = 7.4, 6.7 Hz, 5H), 2.47 (d, $J$ = 8.9 Hz, 2H), 2.40 – 2.33 (m, 3H), 2.25 (d, $J$ = 7.2 Hz, 4H), 2.09 (dq, $J$ = 12.1, 5.8, 4.5 Hz, 4H), 1.75 (s, 3H), 1.73 – 1.58 (m, 12H), 1.53 – 1.40 (m, 16H), 1.29 (m, 5H), 1.17 (s, 3H). $^{13}$C NMR (100 MHz, MeOD) δ 203.0, 197.0, 175.3, 174.9, 173.5, 160.8, 154.0, 126.9, 114.5, 96.0, 72.9, 70.0, 69.3, 65.0, 62.0, 60.3, 58.1, 55.7, 52.9, 39.7, 39.0, 38.9, 35.4, 33.6, 31.7, 30.8, 30.2, 30.0, 29.4, 29.1, 28.7, 28.4, 28.3, 28.2, 27.8, 25.7, 25.5, 23.7, 22.4, 18.8, 13.1. HRMS (ESI/[M+Na]$^{+}$) calcd. for $C_{40}H_{56}N_4NaO_{11}S$: 822.3486, found 822.3484.

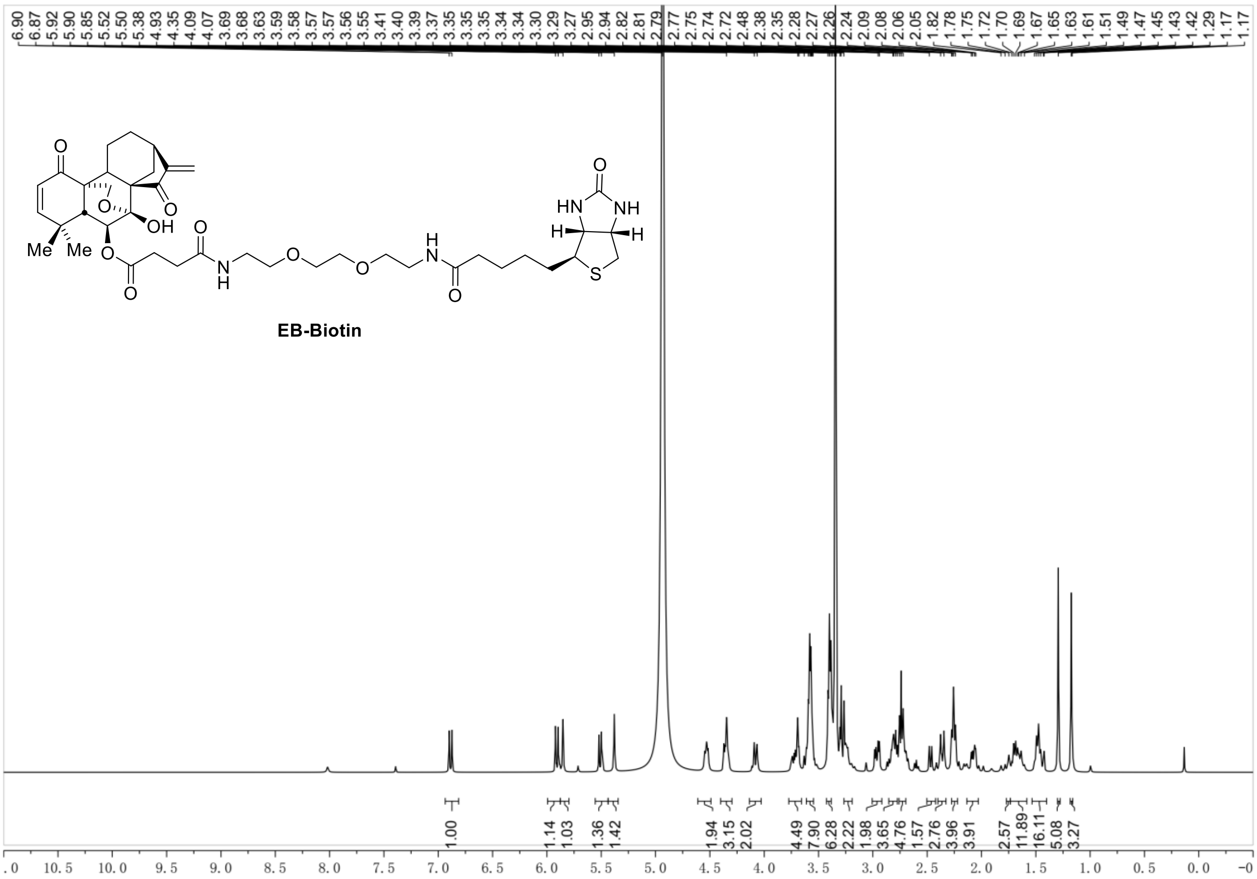
6.90 6.87 5.92 5.90 5.85 5.52 5.50 5.38 4.93 4.35 4.09 4.07 3.69 3.68 3.63 3.59 3.58 3.57 3.57 3.56 3.55 3.41 3.40 3.39 3.37 3.35 3.35 3.35 3.34 3.34 3.30 3.29 3.27 2.95 2.94 2.82 2.81 2.79 2.77 2.75 2.74 2.72 2.48 2.38 2.35 2.28 2.27 2.26 2.24 2.09 2.08 2.06 2.05 1.82 1.78 1.75 1.72 1.70 1.69 1.67 1.65 1.63 1.61 1.51 1.49 1.47 1.45 1.43 1.42 1.29 1.17 1.17
EB-Biotin
1.00 1.14 1.03 1.36 1.42 1.94 3.15 2.02 4.49 7.90 6.28 2.22 1.98 3.65 4.76 1.57 2.76 3.96 3.91 2.57 11.89 16.11 5.08 3.27

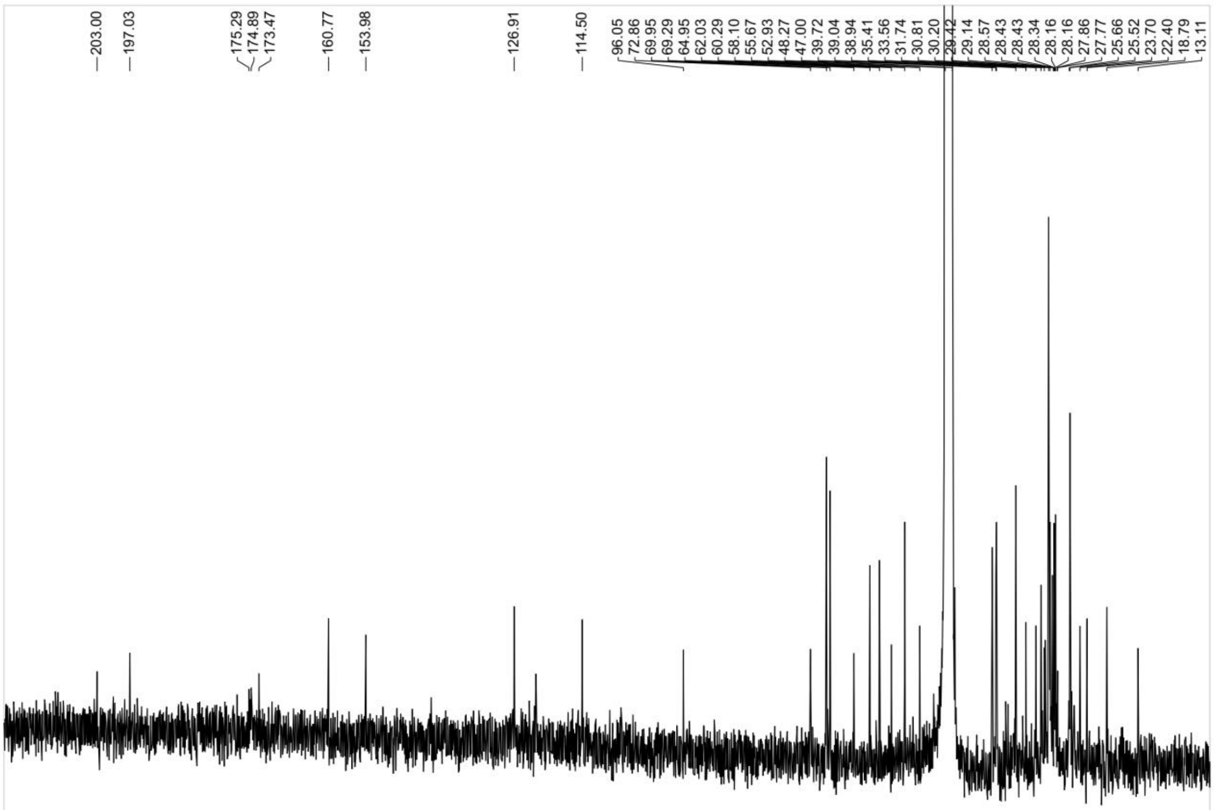
203.00 197.03 175.29 174.89 173.47 160.77 153.98 126.91 114.50 96.05 72.86 69.95 69.29 64.95 62.03 60.29 58.10 55.67 52.93 48.27 47.00 39.72 39.04 38.94 35.41 33.56 31.74 30.81 30.20 29.42 29.14 28.57 28.43 28.34 28.16 27.86 27.77 25.66 25.52 23.70 22.40 18.79 13.11

**Synthesis of Laxiflorin B series analogues**

To a stirred solution of Laxiflorin B (17 mg, 0.05 mmol) in DCM (1 mL) at room temperature was added acid (0.05 mmol), EDCI (9.6 mg, 0.05 mmol) and DMAP (0.6 mg, 0.005 mmol). The resulting solution was stirred another 12 h until LB was disappeared monitored by TLC. The reaction was quenched by addition of saturated an aqueous $NaHCO_3$ (2 mL), followed by saturated brine (10 mL) and ethyl acetate (10 mL). The aqueous phase was extracted by ethyl acetate (10 mL×3) and the combined organic extracts were washed with brine, dried over $Na_2SO_4$, filtered and concentrated under reduced pressure. Silica gel flash column chromatography (silica gel, Hexanes/Ethyl acetate = 1:2 or MeOH/DCM = 1:10) of the residue gave a desired product.

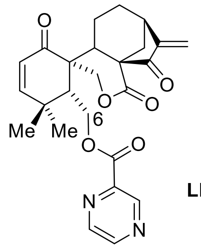


LB-1

**LB-1:**(18 mg, 0.04 mmol, 80%).$^1$H NMR (500 MHz, Chloroform-*d*) δ 9.24 (d, *J* = 1.5 Hz, 1H), 8.79 (d, *J* = 2.4 Hz, 1H), 8.71 (dd, *J* = 2.4, 1.5 Hz, 1H), 6.61 (d, *J* = 10.2 Hz, 1H), 5.96 – 5.90 (m, 2H), 5.44 (s, 1H), 4.83 (d, *J* = 11.2 Hz, 1H), 4.71 (d, *J* = 11.2 Hz, 1H), 4.67 – 4.57 (m, 2H), 3.07 (dd, *J* = 9.4, 4.8 Hz, 1H), 2.72 (d, *J* = 12.6 Hz, 1H), 2.52 – 2.45 (m, 2H), 2.39 (dd, *J* = 13.1, 4.6 Hz, 1H), 2.24 – 2.14 (m, 1H), 1.75 (tdd, *J* = 13.2, 11.9, 7.7 Hz, 1H), 1.56 (ddd, *J* = 13.4, 6.7, 4.6 Hz, 1H), 1.42 (td, *J* = 12.9, 6.7 Hz, 1H), 1.33 (s, 3H), 1.27 (s, 3H). $^{13}$C NMR (125 MHz, $CDCl_3$) δ 202.2, 199.0, 169.4, 163.9, 157.5, 150.7, 148.3, 146.2, 144.7, 142.8, 124.8, 119.0, 69.5, 62.2,

58.5, 51.8, 44.6, 42.3, 36.6, 35.1, 32.0, 30.1, 30.0, 29.8, 24.0, 18.3.HRMS (ESI/[M+Na]+) calcd. for $C_{20}H_{26}O_5Na$: 369.1678, found 369.1682. HRMS (ESI/[M+Na]+) calcd. for $C_{25}H_{26}N_2NaO_6$: 473.1683, found 473.1686

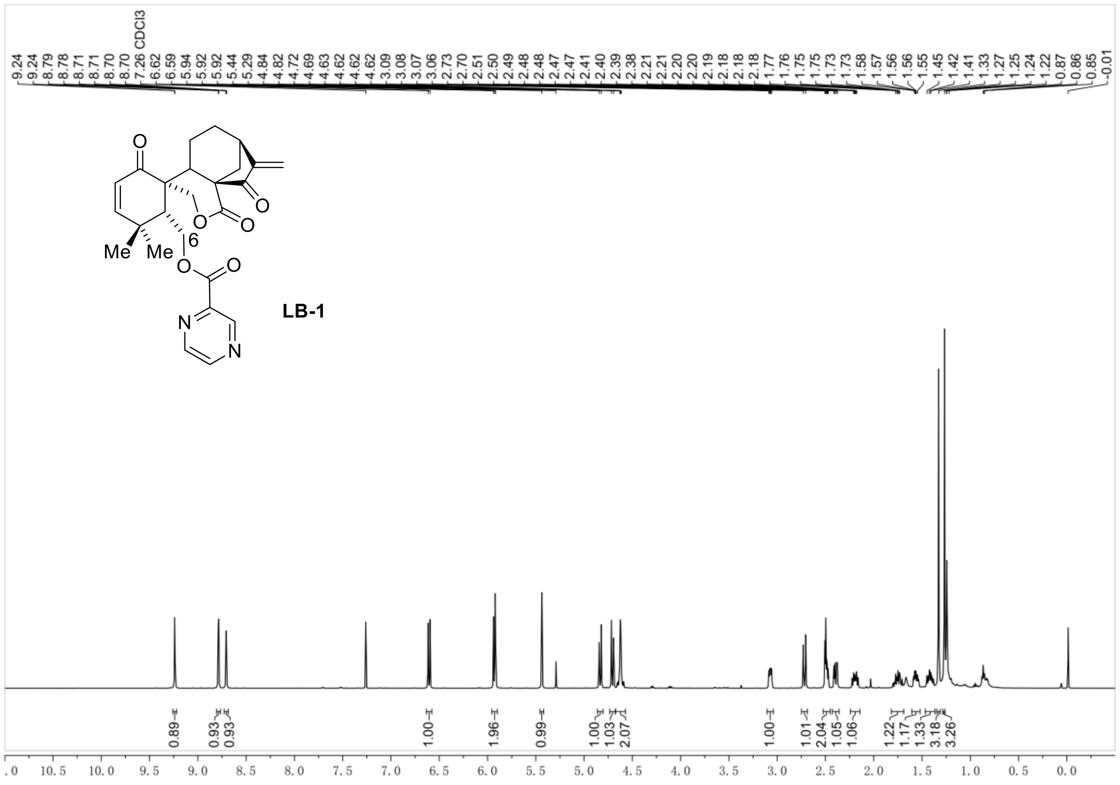


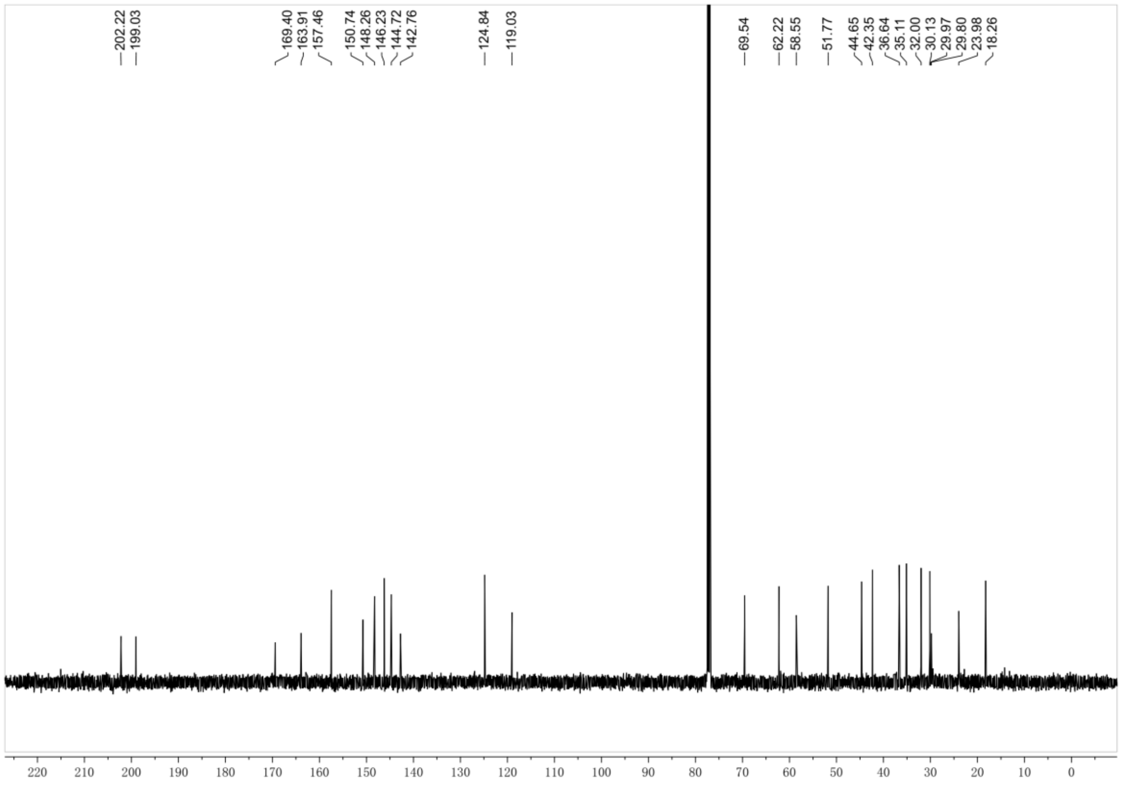

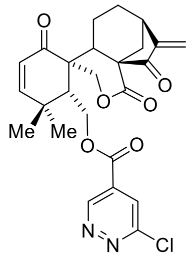

**LB-2**

**LB-2:**(20 mg, 0.041 mmol, 82%).$^1$H NMR (400 MHz, Chloroform-*d*) δ 9.39 (dd, *J* = 5.1, 1.7 Hz, 1H), 8.20 (dd, *J* = 8.5, 1.7 Hz, 1H), 7.71 (dd, *J* = 8.4, 5.1 Hz, 1H), 6.65 (d, *J* = 10.2 Hz, 1H), 5.96 (d, *J* = 10.2 Hz, 1H), 5.88 (s, 1H), 5.44 (s, 1H), 4.73 (d, *J* = 5.3 Hz, 3H), 4.64 (dd, *J* = 12.7, 5.9 Hz, 1H), 3.10 (dd, *J* = 9.4, 4.8 Hz, 1H), 2.74 (d, *J* = 12.6 Hz, 1H), 2.62 (dd, *J* = 6.0, 3.8 Hz, 1H), 2.47 (ddd, *J* = 20.7, 11.9, 5.5 Hz, 2H), 2.23 (dtd, *J* = 14.9, 8.2, 7.3, 4.0 Hz, 1H), 1.84 – 1.68 (m, 2H), 1.57 (td, *J* = 12.5, 11.8, 7.3 Hz, 1H), 1.38 (s, 3H), 1.32 (s, 3H). $^{13}$C NMR (100 MHz, $CDCl_3$) δ 202.5, 199.0, 169.4, 164.0, 157.6, 153.4, 151.1, 150.6, 127.7, 127.1, 124.8, 119.1, 69.3, 62.5, 58.6, 51.5, 44.5, 42.5, 36.5, 35.0, 31.9, 29.9, 29.8, 23.9, 18.2.HRMS (ESI/[M+Na]$^+$) calcd. for $C_{25}H_{25}ClN_2NaO_6$: 507.1293, found 507.1298.

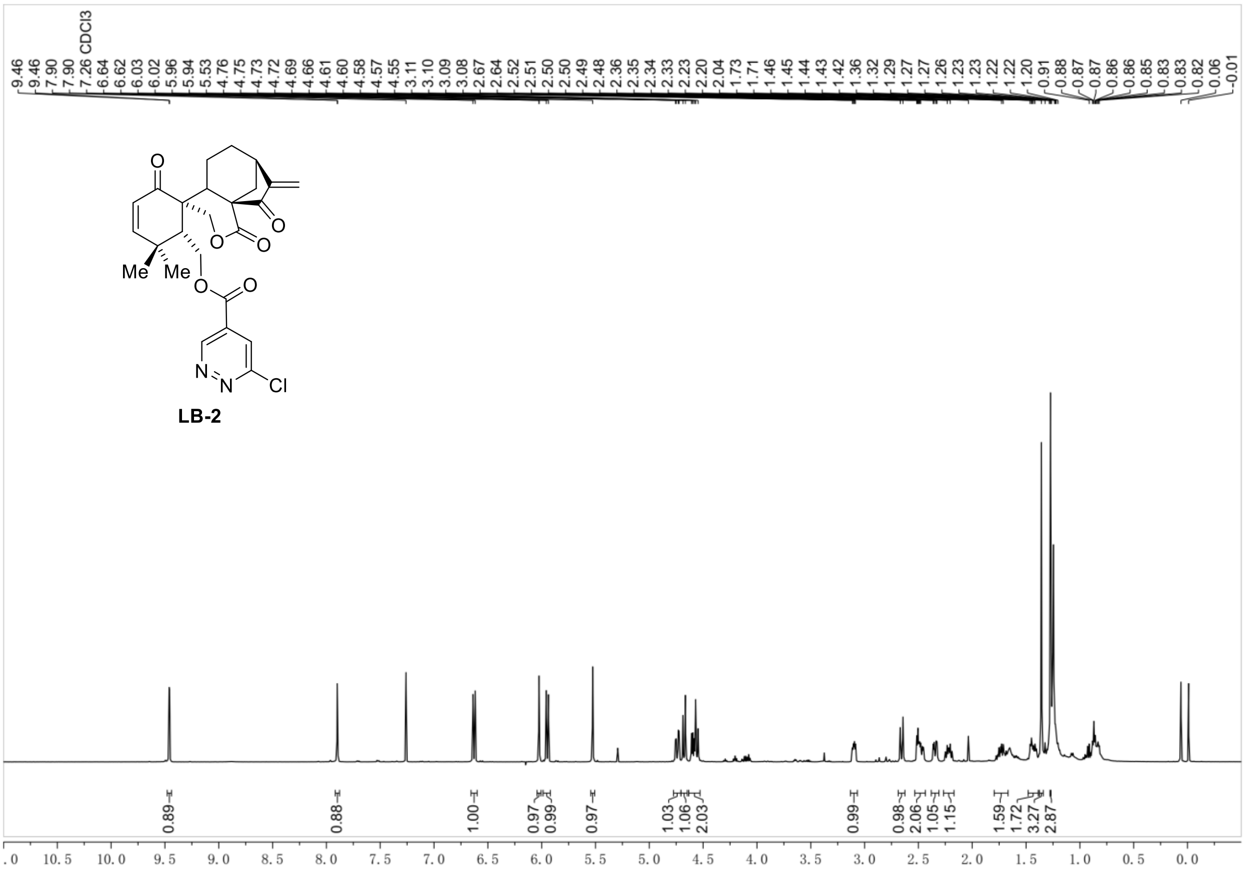
9.46
9.46
7.90
7.90
7.26 CDCl3
6.64
6.62
6.03
6.02
5.96
5.94
5.53
4.76
4.75
4.73
4.72
4.69
4.66
4.61
4.60
4.58
4.57
4.55
3.11
3.10
3.09
3.08
2.67
2.64
2.52
2.51
2.50
2.49
2.48
2.36
2.35
2.34
2.33
2.23
2.20
2.04
1.73
1.71
1.46
1.45
1.44
1.43
1.42
1.36
1.32
1.29
1.27
1.27
1.26
1.23
1.23
1.22
1.22
1.20
0.91
0.88
0.87
0.86
0.86
0.85
0.83
0.83
0.82
0.06
-0.01
O
O
O
O
Me
Me
O
O
N
N
Cl
LB-2
0.89
0.88
1.00
0.97
0.99
0.97
1.03
1.06
2.03
0.99
0.98
2.06
1.05
1.15
1.59
1.72
3.27
2.87
10.5
10.0
9.5
9.0
8.5
8.0
7.5
7.0
6.5
6.0
5.5
5.0
4.5
4.0
3.5
3.0
2.5
2.0
1.5
1.0
0.5
0.0

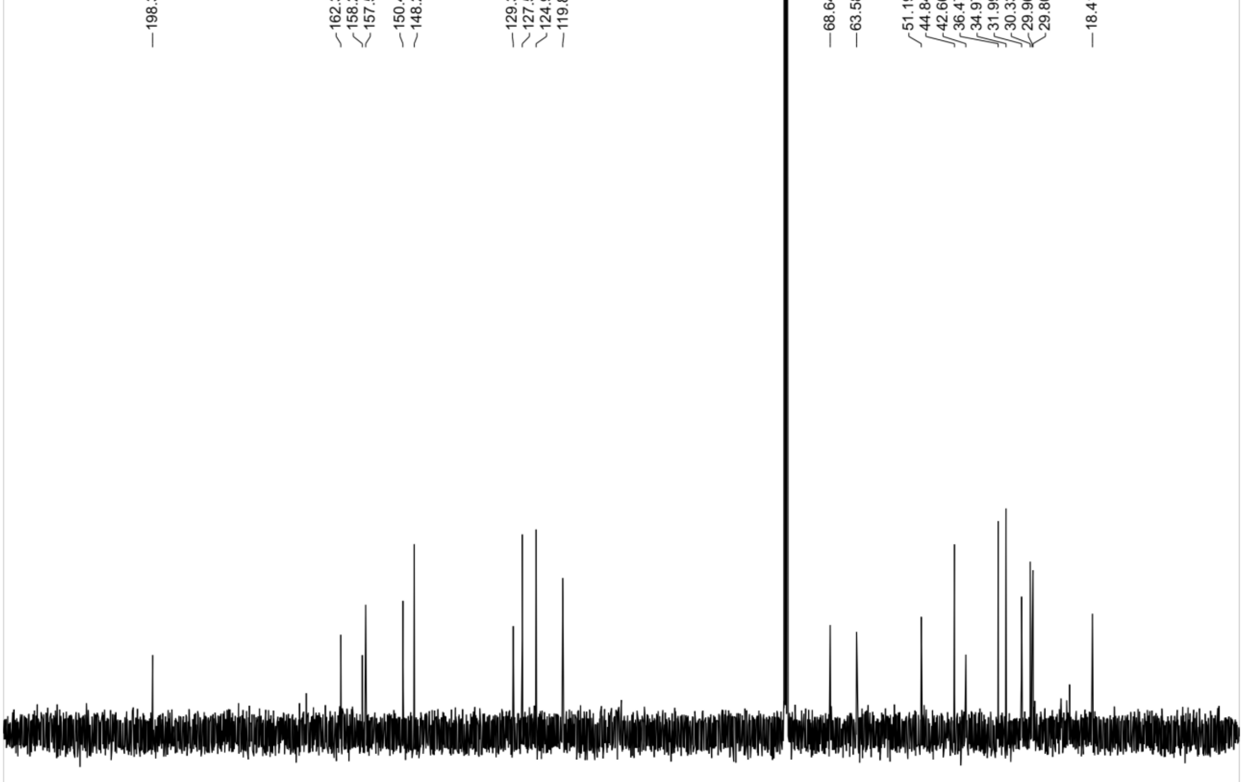
198.38
162.35
158.23
157.58
150.45
148.28
129.32
127.59
124.95
119.85
68.64
63.58
51.19
44.84
42.66
36.47
34.97
31.99
30.33
29.96
29.80
18.41
220
210
200
190
180
170
160
150
140
130
120
110
100
90
80
70
60
50
40
30
20
10
0

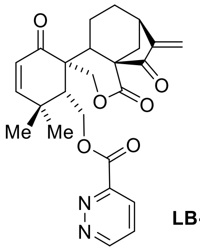

**LB-3**

**LB-3:**(18 mg, 0.04 mmol, 80%).$^{1}$H NMR (500 MHz, Chloroform-*d*) δ 9.46 (d, *J* = 1.8 Hz, 1H), 7.90 (d, *J* = 1.8 Hz, 1H), 6.63 (d, *J* = 10.2 Hz, 1H), 6.02 (d, *J* = 1.2 Hz, 1H), 5.95 (d, *J* = 10.3 Hz, 1H), 5.53 (s, 1H), 4.74 (dd, *J* = 12.8, 3.7 Hz, 1H), 4.68 (d, *J* = 11.2 Hz, 1H), 4.63 – 4.53 (m, 2H), 3.10 (dd, *J* = 9.2, 4.8 Hz, 1H), 2.66 (d, *J* = 12.7 Hz, 1H), 2.49 (ddd, *J* = 17.5, 9.3, 4.2 Hz, 2H), 2.34 (dd, *J* = 13.1, 4.5 Hz, 1H), 2.22 (dt, *J* = 13.5, 8.4 Hz, 1H), 1.80 – 1.67 (m, 1H), 1.48 – 1.39 (m, 2H), 1.36 (s, 3H), 1.27 (s, 3H). $^{13}$C NMR (125 MHz, $CDCl_3$) δ 198.4, 162.3, 158.2, 157.6, 150.5, 148.3, 129.3, 127.6, 124.9, 119.8, 68.6, 63.6, 51.2, 44.8, 42.7, 36.5, 35.0, 32.0, 30.3, 30.0, 29.8, 18.4. HRMS (ESI/[M+Na]$^+$) calcd. for $C_{25}H_{26}N_2NaO_6$: 473.1683, found 473.1685.

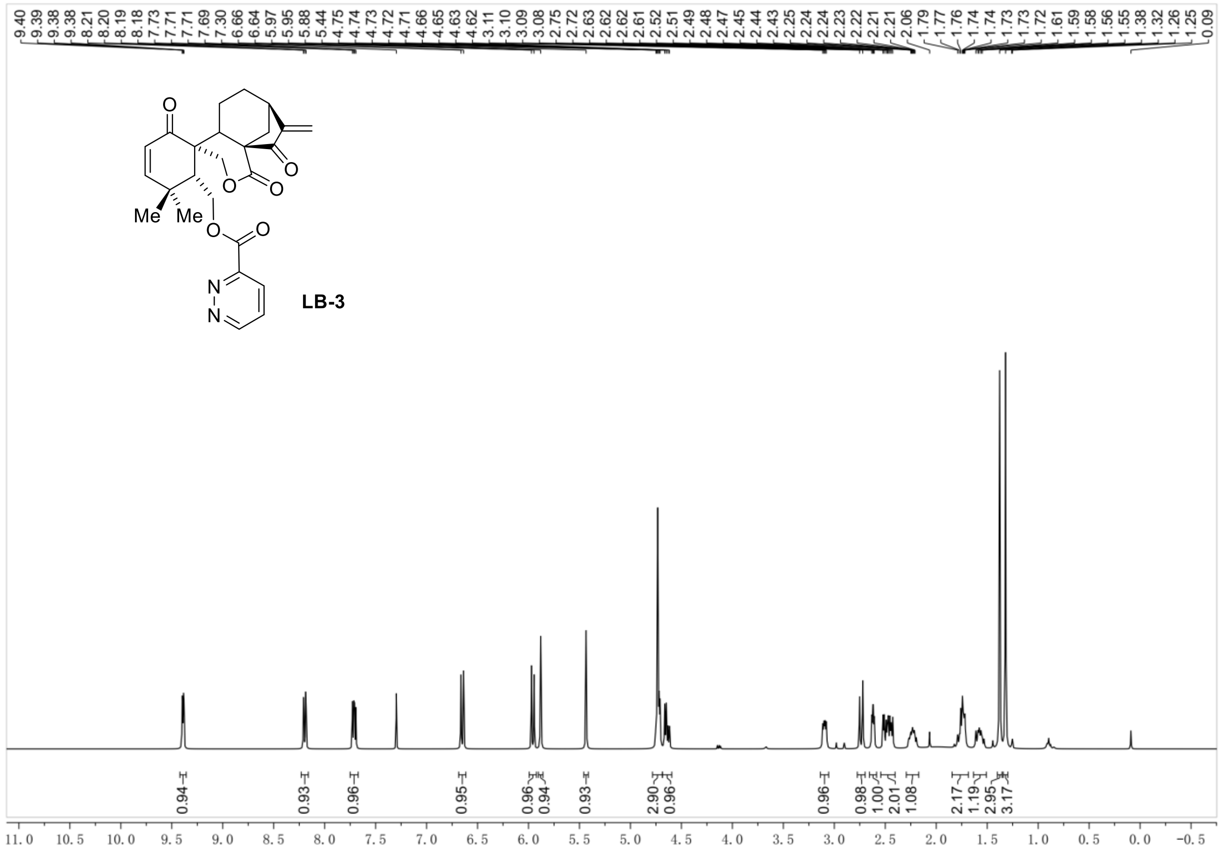
9.40 9.39 9.38 9.38 8.21 8.20 8.19 8.18 7.73 7.71 7.71 7.69 7.30 6.66 6.64 5.97 5.95 5.88 5.44 4.75 4.74 4.73 4.72 4.71 4.66 4.65 4.63 4.62 3.11 3.10 3.09 3.08 2.75 2.72 2.63 2.62 2.61 2.52 2.51 2.49 2.48 2.47 2.45 2.44 2.43 2.25 2.24 2.24 2.23 2.22 2.21 2.21 2.06 1.79 1.77 1.76 1.74 1.74 1.73 1.73 1.72 1.61 1.59 1.58 1.56 1.55 1.38 1.32 1.26 1.25 0.09
O
O
O
O
Me
Me
O
O
N
N
LB-3
0.94 0.93 0.96 0.95 0.96 0.94 0.93 2.90 0.96 0.96 0.98 1.00 2.01 1.08 2.17 1.19 2.95 3.17
11.0 10.5 10.0 9.5 9.0 8.5 8.0 7.5 7.0 6.5 6.0 5.5 5.0 4.5 4.0 3.5 3.0 2.5 2.0 1.5 1.0 0.5 0.0 -0.5

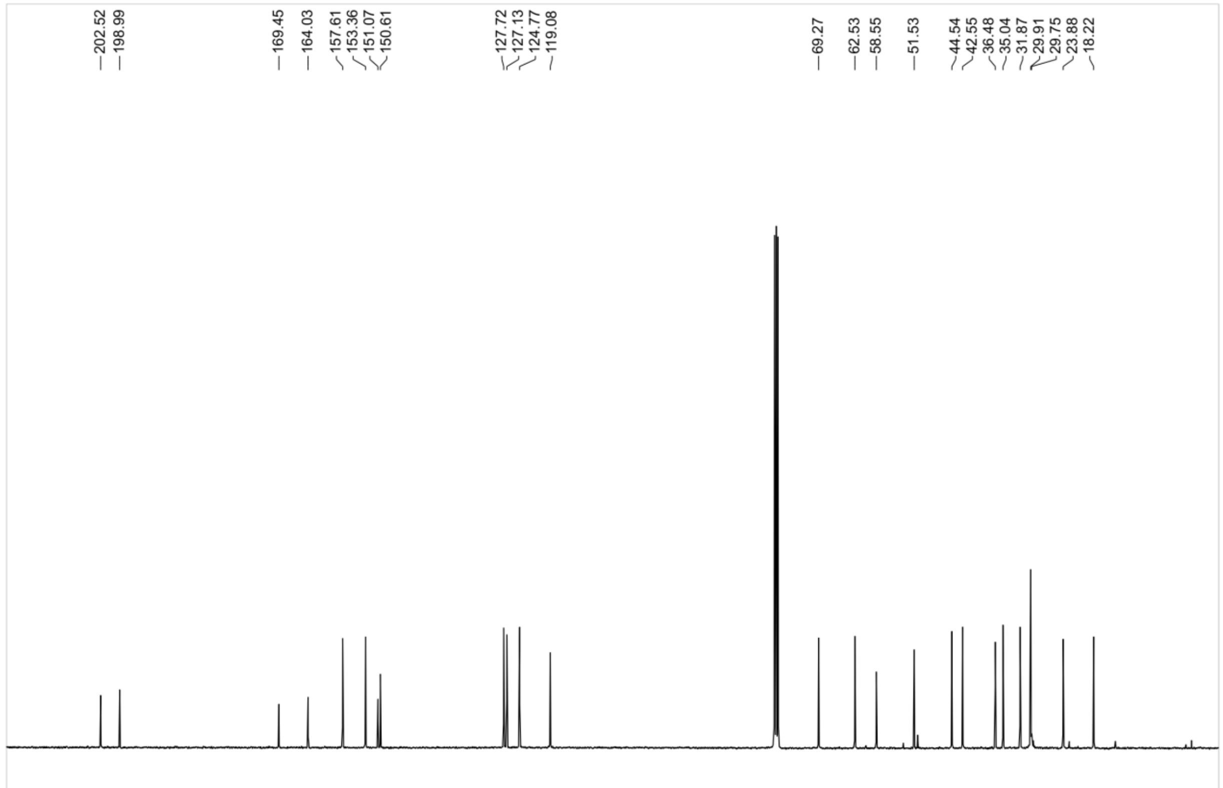
202.52 198.99 169.45 164.03 157.61 153.36 151.07 150.61 127.72 127.13 124.77 119.08 69.27 62.53 58.55 51.53 44.54 42.55 36.48 35.04 31.87 29.91 29.75 23.88 18.22
20 210 200 190 180 170 160 150 140 130 120 110 100 90 80 70 60 50 40 30 20 10 0

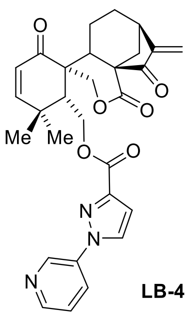


**LB-4**

**LB-4:**(18 mg, 0.035 mmol, 70%).[1]H NMR (500 MHz, Chloroform-*d*) δ 9.12 (s, 1H), 8.73 – 8.59 (m, 1H), 8.50 (s, 1H), 8.18 (ddd, *J* = 8.3, 2.6, 1.4 Hz, 1H), 8.10 (s, 1H), 7.52 – 7.44 (m, 1H), 6.61 (d, *J* = 10.2 Hz, 1H), 6.00 (s, 1H), 5.93 (d, *J* = 10.2 Hz, 1H), 5.48 (s, 1H), 4.79 (d, *J* = 11.1 Hz, 1H), 4.76 – 4.68 (m, 2H), 4.43 (dd, *J* = 12.8, 4.2 Hz, 1H), 3.11 (dd, *J* = 9.3, 4.8 Hz, 1H), 2.72 (d, *J* = 12.6 Hz, 1H), 2.53 – 2.35 (m, 3H), 2.23 (dt, *J* = 13.8, 8.5 Hz, 1H), 1.94 – 1.63 (m, 1H), 1.25 (s, 11H). [13]C NMR (125 MHz, $CDCl_3$) δ 202.7, 199.0, 169.3, 161.8, 157.7, 150.8, 149.0, 142.9, 141.1, 130.5, 127.2, 124.8, 124.3, 119.4, 116.5, 69.5, 60.7, 51.7, 44.8, 42.3, 36.7, 35.1, 32.0, 30.3, 29.8, 29.8, 29.5, 22.8, 18.3, 14.2, 1.1, 0.1.HRMS (ESI/$[M+Na]^+$) calcd. for $C_{29}H_29N_3NaO_6$: 538.1949, found 538.1956.

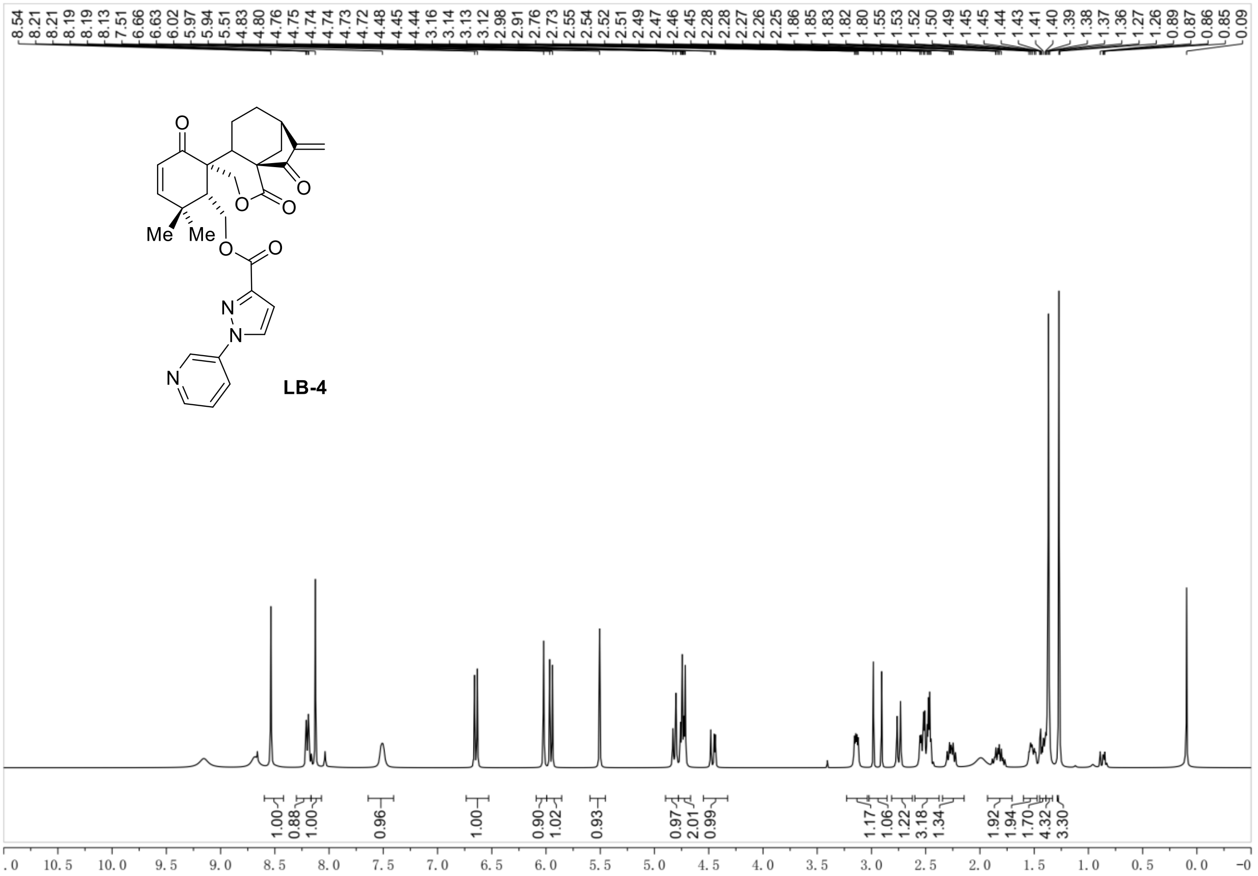
8.54 8.21 8.21 8.19 8.19 8.13 7.51 6.66 6.63 6.02 5.97 5.94 5.51 4.83 4.80 4.76 4.75 4.74 4.74 4.73 4.72 4.48 4.45 4.44 3.16 3.14 3.13 3.12 2.98 2.91 2.76 2.73 2.55 2.54 2.52 2.51 2.49 2.47 2.46 2.45 2.28 2.28 2.27 2.26 2.25 1.86 1.85 1.83 1.82 1.80 1.55 1.53 1.52 1.50 1.49 1.45 1.45 1.44 1.43 1.41 1.40 1.39 1.38 1.37 1.36 1.27 1.26 0.89 0.87 0.86 0.85 0.09
Me
Me
LB-4
1.00 0.88 1.00 0.96 1.00 0.90 1.02 0.93 0.97 2.01 0.99 1.17 1.06 1.22 3.18 1.34 1.92 1.94 1.70 4.32 3.30
10.5 10.0 9.5 9.0 8.5 8.0 7.5 7.0 6.5 6.0 5.5 5.0 4.5 4.0 3.5 3.0 2.5 2.0 1.5 1.0 0.5 0.0

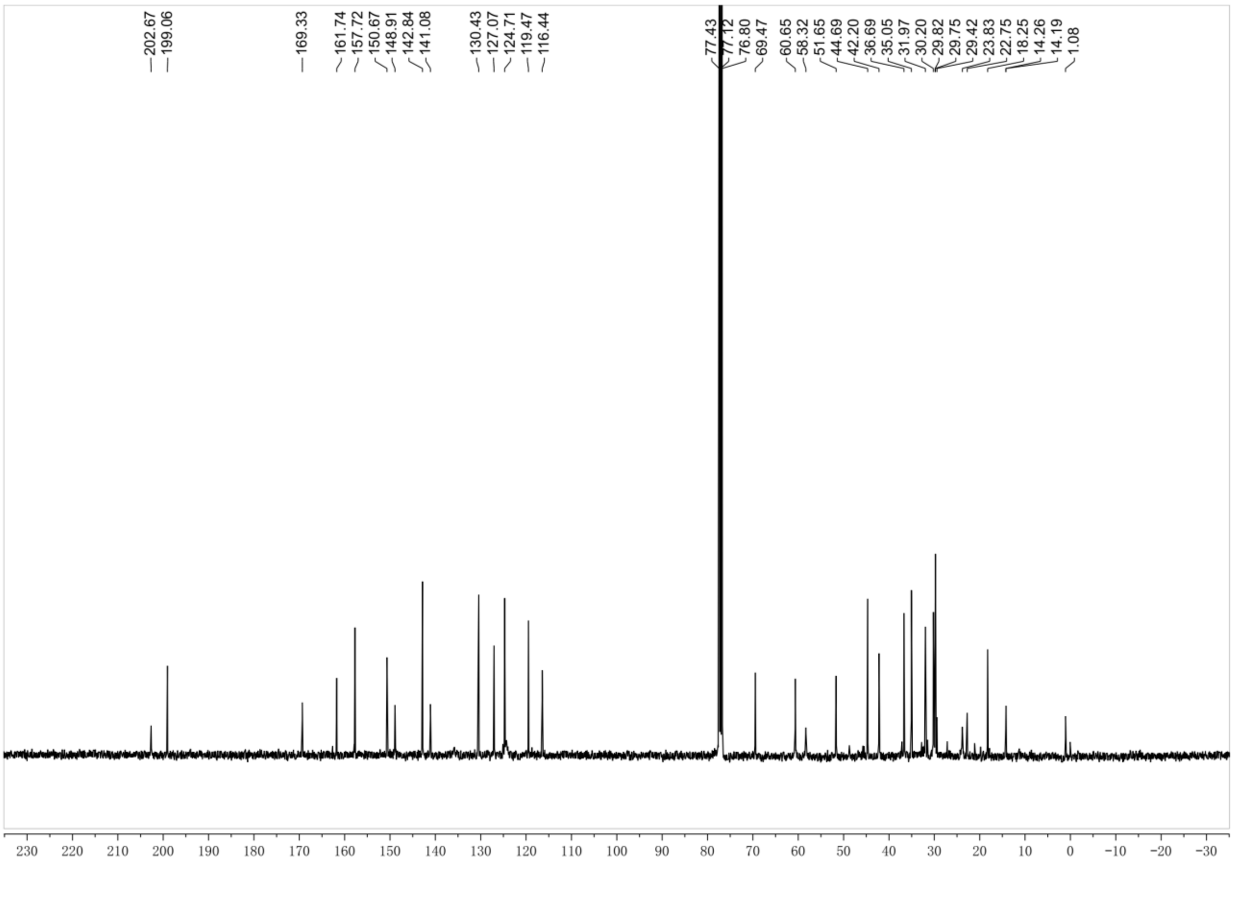
202.67 199.06 169.33 161.74 157.72 150.67 148.91 142.84 141.08 130.43 127.07 124.71 119.47 116.44 77.43 77.12 76.80 69.47 60.65 58.32 51.65 44.69 42.20 36.69 35.05 31.97 30.20 29.82 29.75 29.42 23.83 22.75 18.25 14.26 14.19 1.08
230 220 210 200 190 180 170 160 150 140 130 120 110 100 90 80 70 60 50 40 30 20 10 0 −10 −20 −30

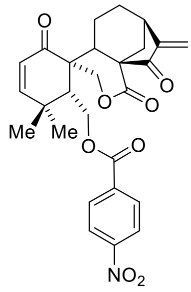

LB-5

**LB-5:**(18 mg, 0.038 mmol, 75%).$^{1}$H NMR (400 MHz, Chloroform-*d*) δ 8.37 – 8.26 (m, 2H), 8.19 – 8.08 (m, 2H), 6.63 (d, *J* = 10.2 Hz, 1H), 6.02 (s, 1H), 5.94 (d, *J* = 10.3 Hz, 1H), 5.49 (s, 1H), 4.72 – 4.56 (m, 3H), 3.09 (dd, *J* = 9.2, 4.8 Hz, 1H), 2.69 (d, *J* = 12.7 Hz, 1H), 2.54 – 2.45 (m, 2H), 2.41 (dd, *J* = 13.2, 4.5 Hz, 1H), 2.19 (dt, *J* = 13.9, 8.5 Hz, 1H), 1.75 (qd, *J* = 13.0, 7.8 Hz, 1H), 1.46 (dt, *J* = 12.9, 5.5 Hz, 1H), 1.37 (s, 3H), 1.28 (s, 3H). $^{13}$C NMR (100 MHz, $CDCl_3$) δ 202.3, 198.8, 169.1, 164.4, 157.6, 151.0, 150.8, 134.6, 130.9, 124.9, 124.0, 119.4, 69.1, 62.4, 58.4, 51.5, 44.7, 42.5, 36.6, 35.0, 32.0, 30.3, 29.9, 29.8, 24.3, 18.3.HRMS (ESI/$[M+Na]^+$) calcd. for $C_{27}H_{27}NNaO_8$: 516.1629, found 516.1633.

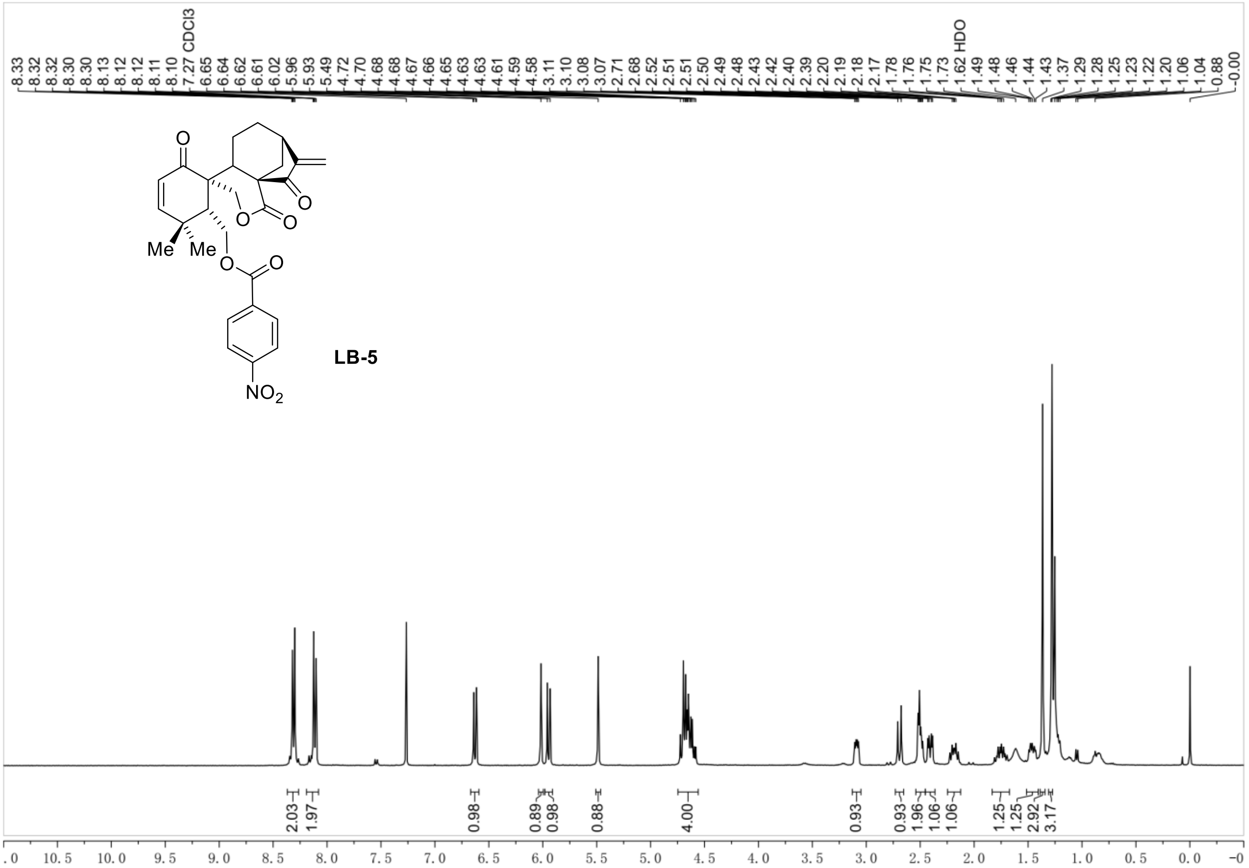
8.33 8.32 8.32 8.30 8.30 8.13 8.12 8.12 8.11 8.10 7.27 CDCl3 6.65 6.64 6.62 6.61 6.02 5.96 5.93 5.49 4.72 4.70 4.68 4.68 4.67 4.66 4.65 4.63 4.63 4.61 4.59 4.58 3.11 3.10 3.08 3.07 2.71 2.68 2.52 2.51 2.51 2.50 2.49 2.48 2.43 2.42 2.40 2.39 2.20 2.19 2.18 2.17 1.78 1.76 1.75 1.73 1.62 HDO 1.49 1.48 1.46 1.44 1.43 1.37 1.29 1.28 1.25 1.23 1.22 1.20 1.06 1.04 0.88 -0.00
Me
Me
$NO_2$
LB-5
2.03 1.97 0.98 0.89 0.98 0.88 4.00 0.93 0.93 1.96 1.06 1.06 1.25 1.25 2.92 3.17
.0 10.5 10.0 9.5 9.0 8.5 8.0 7.5 7.0 6.5 6.0 5.5 5.0 4.5 4.0 3.5 3.0 2.5 2.0 1.5 1.0 0.5 0.0 −0

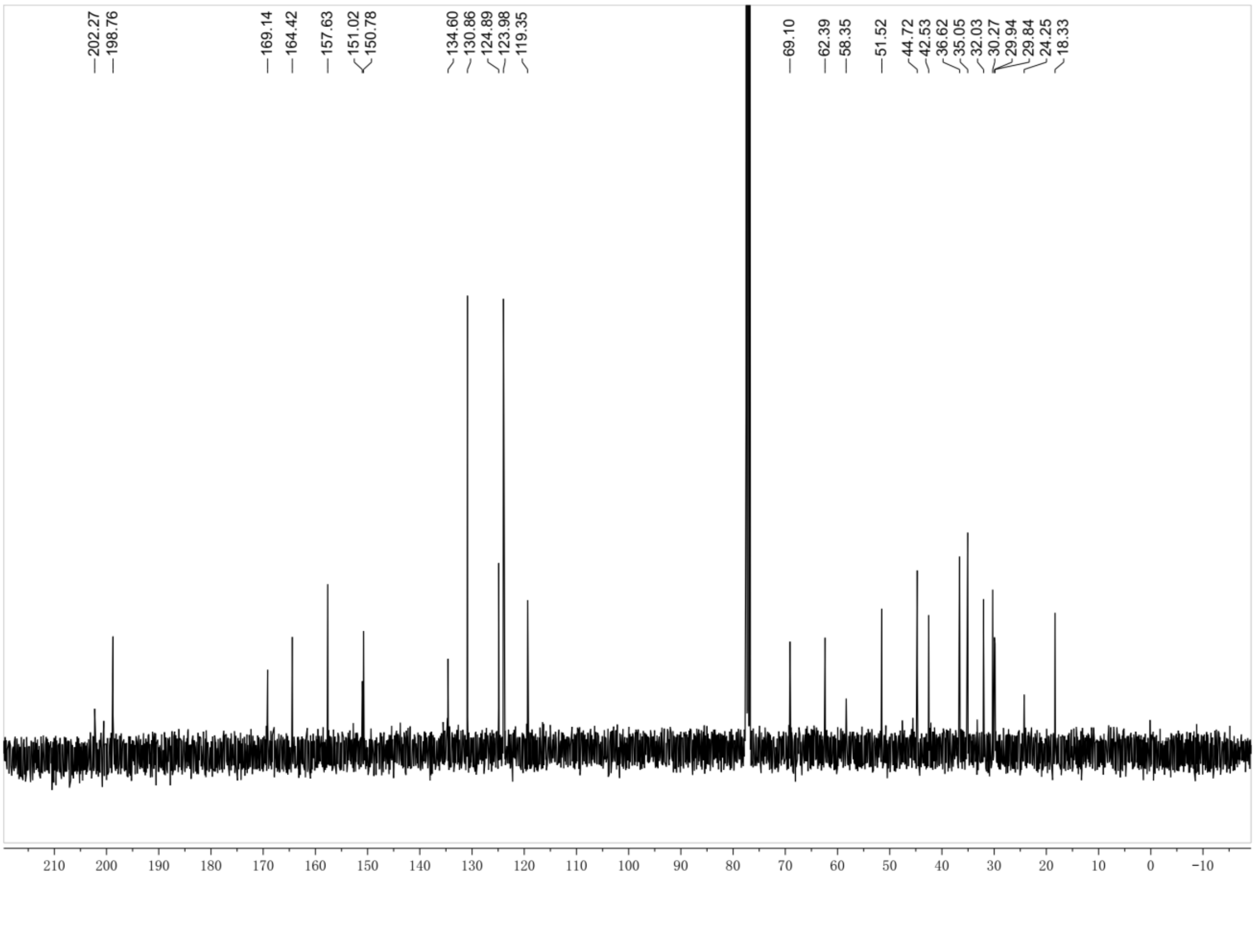
202.27 198.76 169.14 164.42 157.63 151.02 150.78 134.60 130.86 124.89 123.98 119.35 69.10 62.39 58.35 51.52 44.72 42.53 36.62 35.05 32.03 30.27 29.94 29.84 24.25 18.33
210 200 190 180 170 160 150 140 130 120 110 100 90 80 70 60 50 40 30 20 10 0 −10

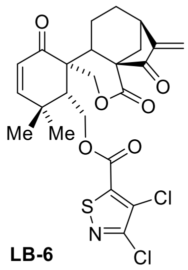

**LB-6**

**LB-6:**(22 mg, 0.0425 mmol, 85%).$^{1}$H NMR (400 MHz, Chloroform-*d*) δ 6.62 (d, *J* = 10.3 Hz, 1H), 6.03 (s, 1H), 5.94 (d, *J* = 10.2 Hz, 1H), 5.52 (s, 1H), 4.71 – 4.62 (m, 2H), 4.62 – 4.54 (m, 2H), 3.10 (dd, *J* = 9.3, 4.8 Hz, 1H), 2.70 (d, *J* = 12.6 Hz, 1H), 2.50 (dt, *J* = 8.9, 4.6 Hz, 2H), 2.31 – 2.16 (m, 2H), 1.82 – 1.66 (m, 1H), 1.51 – 1.42 (m, 1H), 1.45 – 1.34 (m, 1H), 1.34 (s, 3H), 1.29 (s, 6H), 1.26 (d, *J* = 7.1 Hz, 11H), 1.22 (s, 1H), 0.91 (s, 1H), 0.91 – 0.83 (m, 3H).$^{13}$C NMR (100 MHz, $CDCl_3$) δ 202.2, 198.5, 169.1, 157.5, 151.0, 150.6, 148.4, 127.0, 124.9, 119.7, 69.0, 62.9, 58.4, 51.4, 44.7, 42.5, 36.6, 35.0, 32.1, 30.2, 30.0, 29.8, 29.5, 24.3, 22.8, 18.3, 14.3. HRMS (ESI/$[M+Na]^+$) calcd. for $C_{24}H_{23}Cl_2NNaO_6S$: 546.0515, found 546.0518

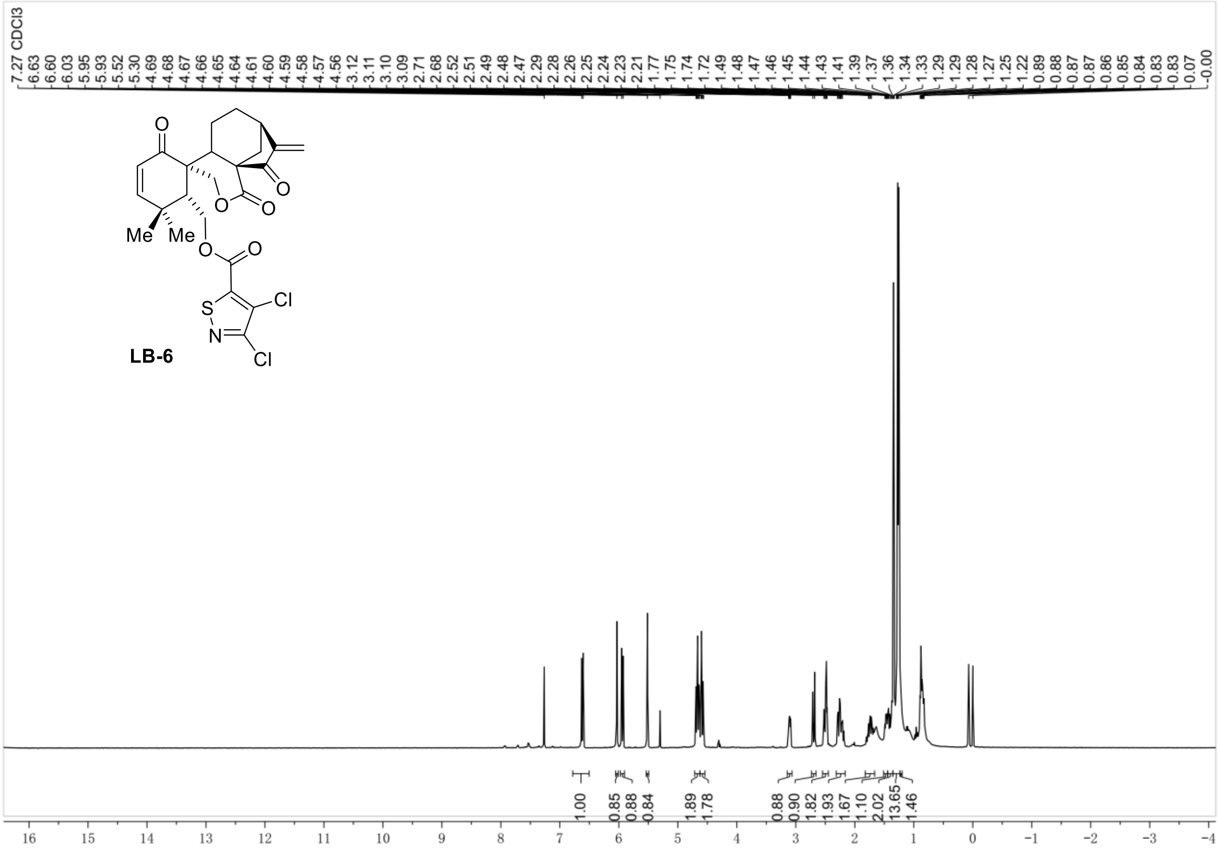
7.27 CDCl3
LB-6
Me
Me
O
S
N
Cl
Cl
1.00
0.85
0.88
0.84
1.89
1.78
0.88
0.90
1.82
1.93
1.67
1.10
2.02
13.65
1.46

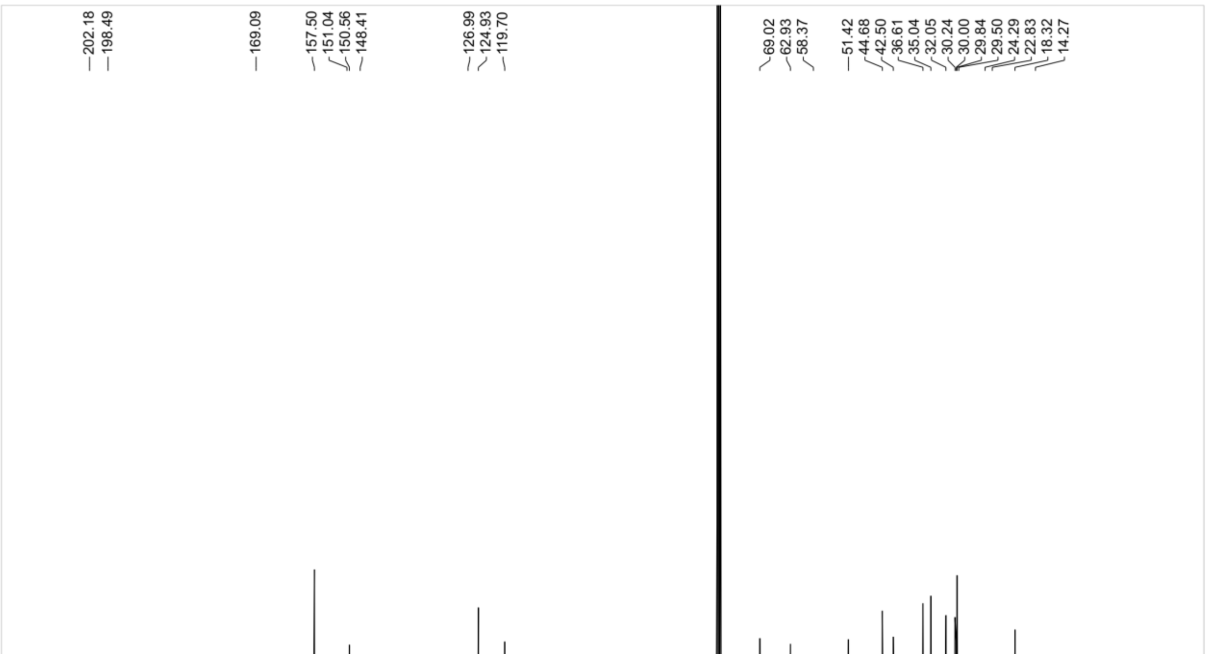
202.18
198.49
169.09
157.50
151.04
150.56
148.41
126.99
124.93
119.70
69.02
62.93
58.37
51.42
44.68
42.50
36.61
35.04
32.05
30.24
30.00
29.84
29.50
24.29
22.83
18.32
14.27

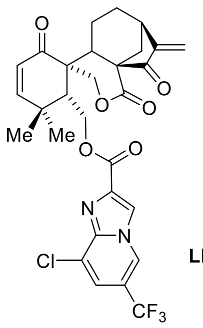

**LB-7**

**LB-7:**(22 mg, 0.0375 mmol, 75%).[1]H NMR (400 MHz, Chloroform-*d*) δ 8.60 – 8.55 (m, 1H), 8.38 (s, 1H), 7.51 (d, *J* = 1.4 Hz, 1H), 6.61 (d, *J* = 10.2 Hz, 1H), 5.96 – 5.89 (m, 2H), 5.43 (s, 1H), 4.84 – 4.73 (m, 2H), 4.72 (d, *J* = 11.0 Hz, 1H), 4.51 (dd, *J* = 12.8, 4.1 Hz, 1H), 3.10 (dd, *J* = 9.2, 4.7 Hz, 1H), 2.75 (d, *J* = 12.5 Hz, 1H), 2.54 – 2.41 (m, 3H), 2.21 (dt, *J* = 13.6, 8.0 Hz, 1H), 1.79 (qd, *J* = 13.4, 13.0, 7.2 Hz, 1H), 1.58 (ddt, *J* = 18.2, 12.8, 6.2 Hz, 2H), 1.25 (s, 8H), 0.07 (s, 1H). $^{13}$C NMR (100 MHz, $CDCl_3$) δ 199.1, 169.5, 161.6, 157.7, 150.8, 142.9, 137.8, 126.3, 124.8, 121.4, 120.3, 119.2, 118.6, 69.7, 61.5, 58.6, 51.8, 44.7, 42.4, 36.8, 35.2, 32.1, 30.1, 30.0, 29.8, 29.5, 23.9, 22.8, 18.3, 14.3.HRMS (ESI/[M+Na]$^+$) calcd. for $C_{29}H_{26}ClF_3N_2NaO_6$: 613.1324, found 613.1327.

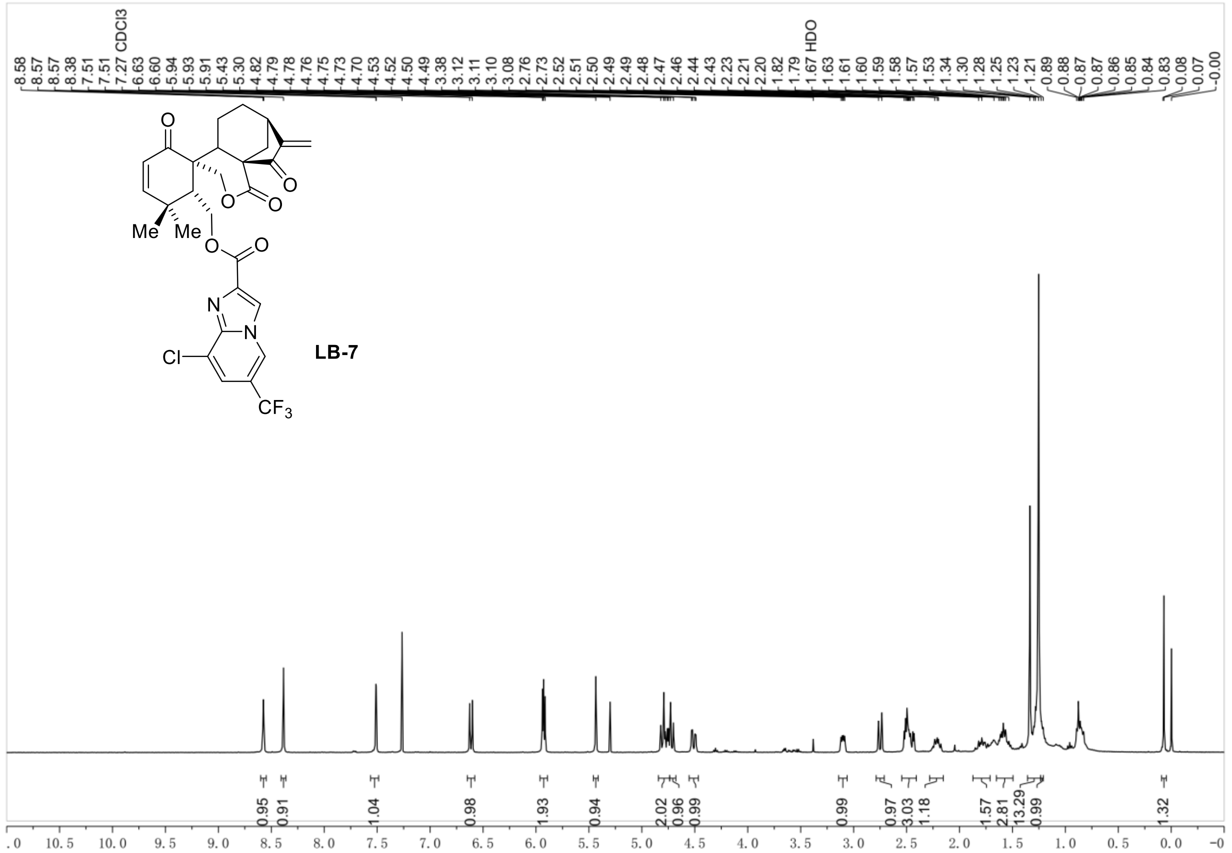

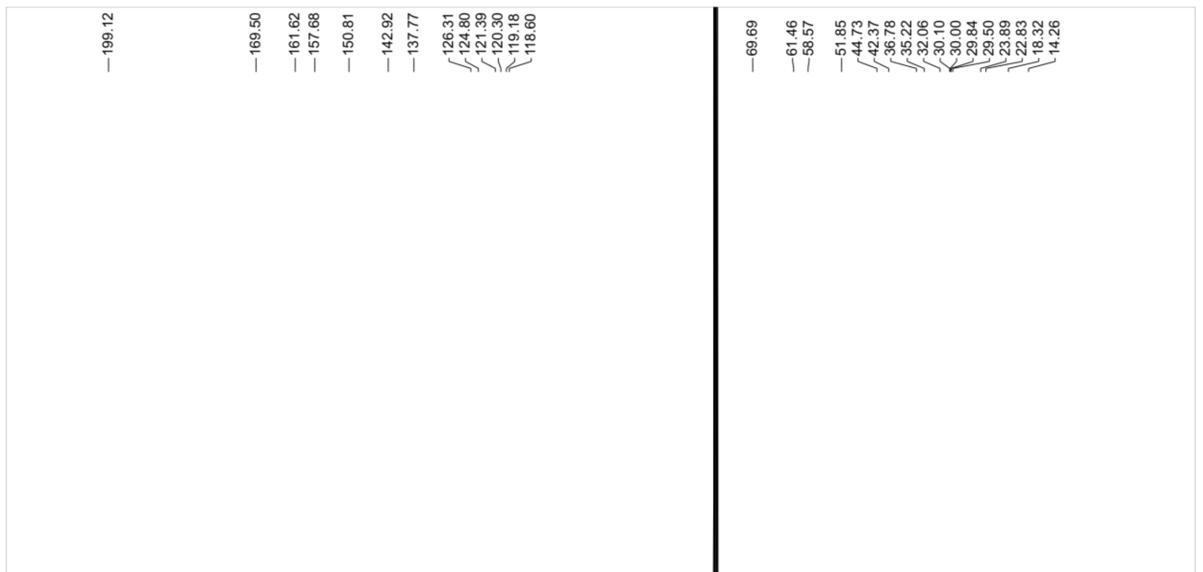
