## Supplementary Figures and Tables for "A Novel Selective ERK1/2 Inhibitor, Laxiflorin B, Targets EGFR Mutation Subtypes in Non-small-cell Lung Cancer"

### List of Contents (This file contains Supplementary Figure 1-12 and Table 1-5):

**Supplementary Figure S1.** Anti-cancer effect of Laxiflorin B in NSCLC cell lines

**Supplementary Figure S2.** Analysis of the effects of Laxiflorin B treatment signaling pathways and gene expression in NSCLC

**Supplementary Figure S3.** The effects of Laxiflorin B treatment in MAPKs and STAT-related pathways

**Supplementary Figure S4.** Identification of Laxiflorin B targeting site on ERK1 by LC-MS/MS

**Supplementary Figure S5.** The structure of biotin-labeled Laxiflorin B analogues

**Supplementary Figure S6.** Influence of Laxiflorin A on NSCLC cell lines

**Supplementary Figure S7.** The root means square deviation (RMSD) and probability of hydrogen bond formation between Laxiflorin B and ERK1/2

**Supplementary Figure S8.** Laxiflorin B sensitivity of BAD-KD (knockdown) NSCLC cells

**Supplementary Figure S9.** The influence of Laxiflorin B on the ErbB pathway

**Supplementary Figure S10.** In vivo study and toxicity of Laxiflorin B

**Supplementary Figure S11.** The modifiable capacity of Laxiflorin B

**Supplementary Table S1.** Down-regulated genes by Laxiflorin B treatment in PC9

**Supplementary Table S2.** $IC_{50}$ of Laxiflorin B in PC9 and HCC827 Bad-knockdown stable cells

**Supplementary Table S3.** IC50 of Laxiflorin B analogous in non-small cellular lung cancer cell lines

**Supplementary Table S4.** $IC_{50}$ of Laxiflorin B analogous in non-small cellular lung cancer cell lines

**Supplementary Table S5.** $IC_{50}$ of Laxiflorin B analogous comparing with ERK inhibitors in non-small cellular lung cancer cell lines

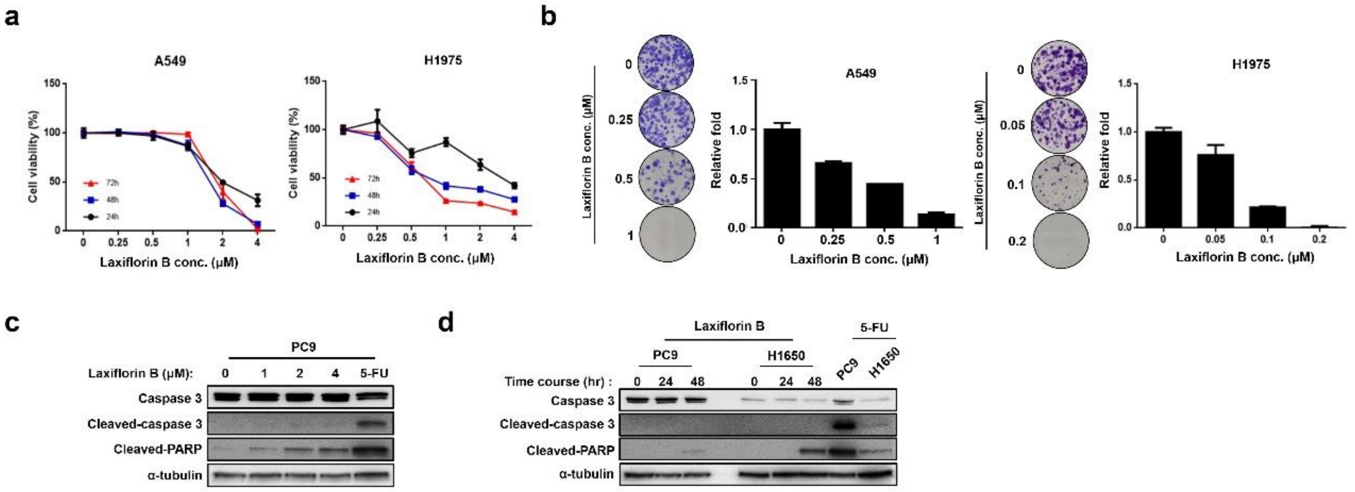


**Supplementary Figure S1. Anti-cancer effect of Laxiflorin B in NSCLC cell lines**

**a** Viability of A549 and H1975 cells after Laxiflorin B treatment. **b** Clonogenic assays of A549 and H1975 cell growth after Laxiflorin B treatment for 2 and 3 weeks, respectively. **c-d** Western blot analysis of the expression of the apoptosis-related proteins, caspase 3 and PARP in PC9 cells after Laxiflorin B treatment for 48 h or in PC9 and H1650 cells after treatment with 2 μM Laxiflorin B for 24 h and 48 h, respectively; 5-FU was used as a positive control. ***P* < 0.01.

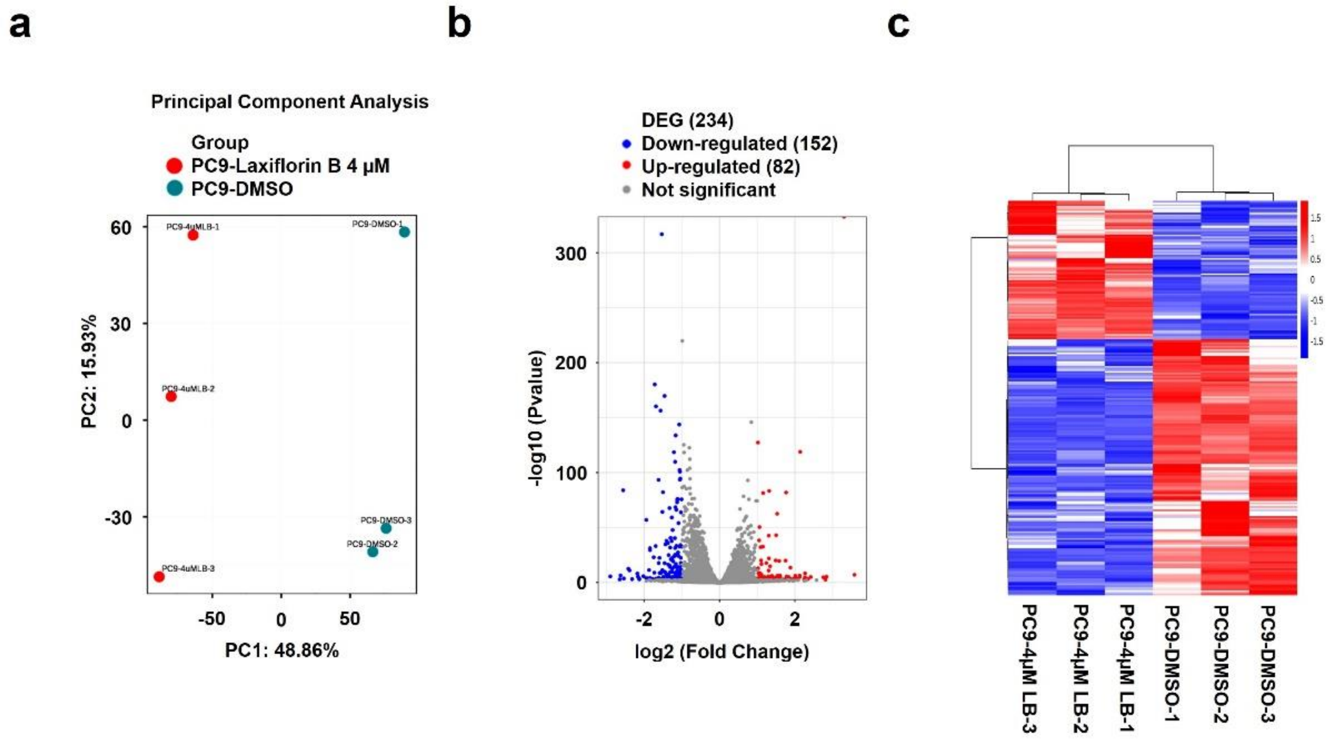


**Supplementary Figure S2. Analysis of the effects of Laxiflorin B treatment signaling pathways and gene expression in NSCLC.**

**a** Classification of control and Laxiflorin B-treated groups in PC9 was categorized by Principle Component Analysis of gene expression in PC9 cells after Laxiflorin B treatment; DMSO was used as a control. **b** Volcano plot showing the changes in gene expression in PC9 cells after Laxiflorin B treatment. **c** Cluster analysis of differentially expressed genes after Laxiflorin B treatment.

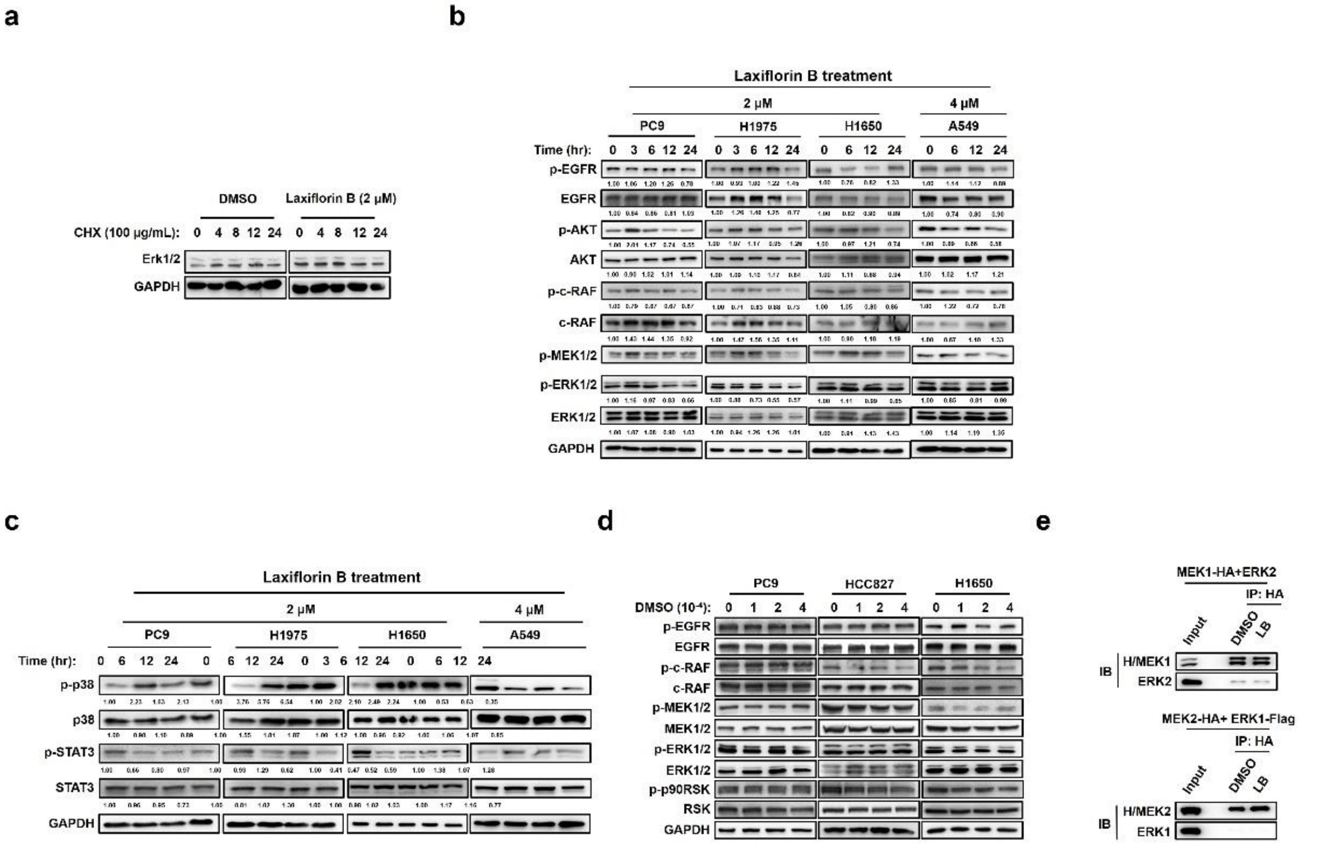


**Supplementary Figure S3. The effects of Laxiflorin B treatment in MAPKs and STAT-related pathways**

**a** Western blot analysis of ERK1/2 protein stability after Laxiflorin B treatment. **b** Western blot analysis of the expression of EGFR-associated molecules in PC9, H1975, H1650 and A549 cells after Laxiflorin B treatment. **c** Western blot analysis of the p38 and STAT3 signaling pathway status in PC9, H1975, H1650 and A549 cells after Laxiflorin B treatment. **d** Western blot analysis of the expression of EGFR-associated molecules in PC9, HCC827 and H1650 after DMSO treatment for 24 h. **e** Immunoprecipitation analysis of the interaction of MEK1 with ERK2 or MEK2 with ERK1 after Laxiflorin B treatment in HEK293 cells.

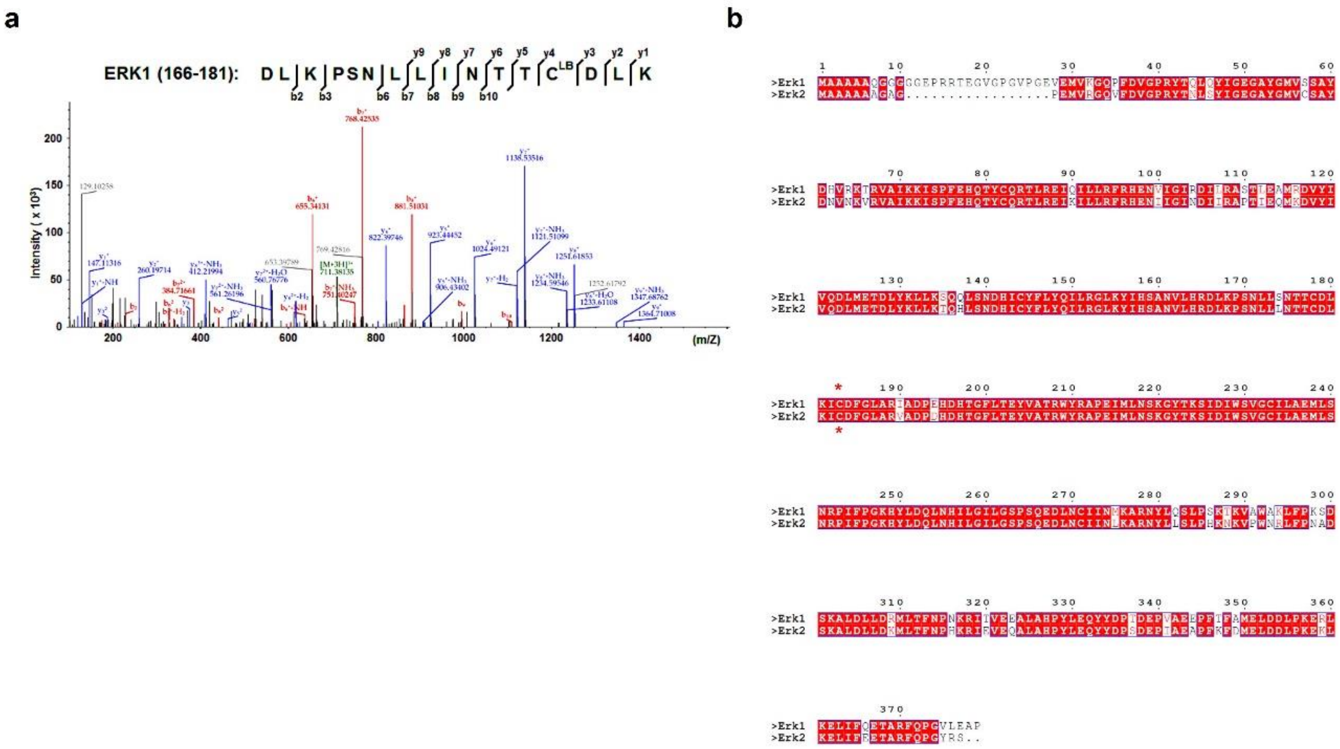


**Supplementary Figure S4. Identification of Laxiflorin B targeting site on ERK1 by LC-MS/MS**

**a** LC-MS/MS analysis showing that Laxiflorin B covalently binds Cys-178 of ERK1. **b** The homologues of ERKs show a high level of similarity in the peptide sequence of the ATP-binding pocket around Cys-183 of ERK1 corresponding to Cys-166 of ERK2.

**Laxiflorin B-Biotin**

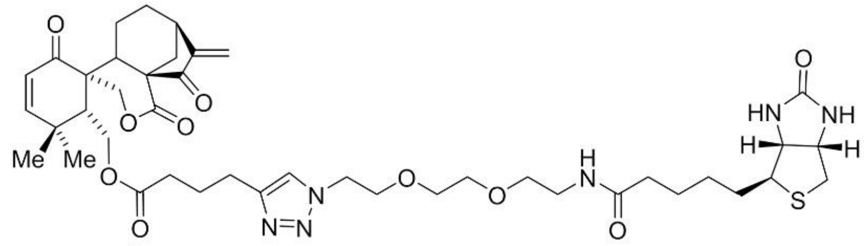

**Laxiflorin A-Biotin**

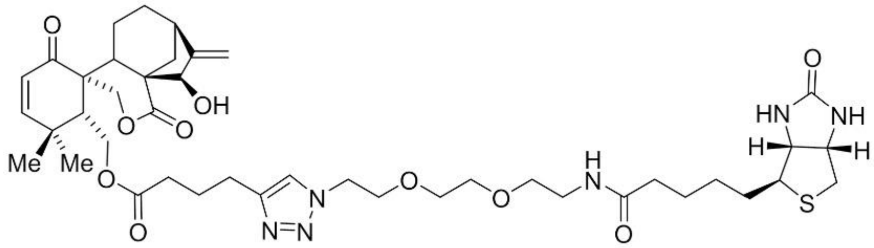

**Eriocalyxin B-Biotin**

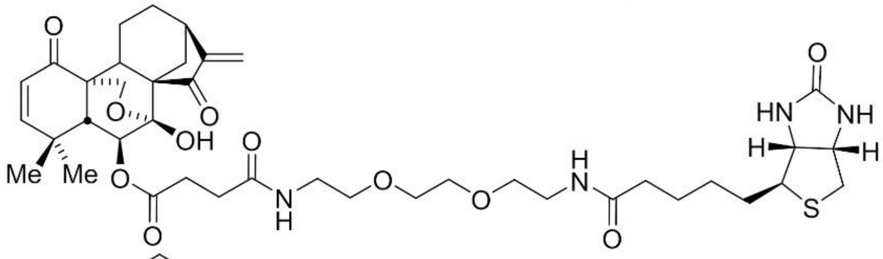

**Laxiflorin J-Biotin**

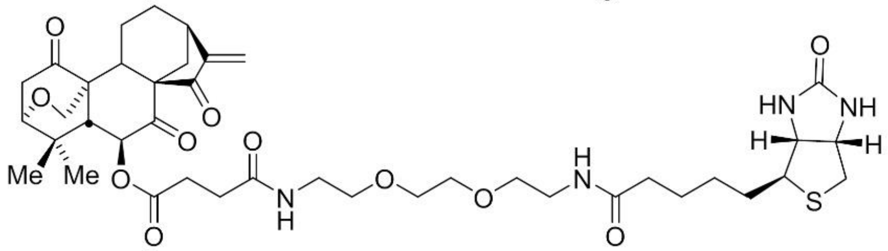

**Laxiflorin B-Di-PEG2-Biotin**

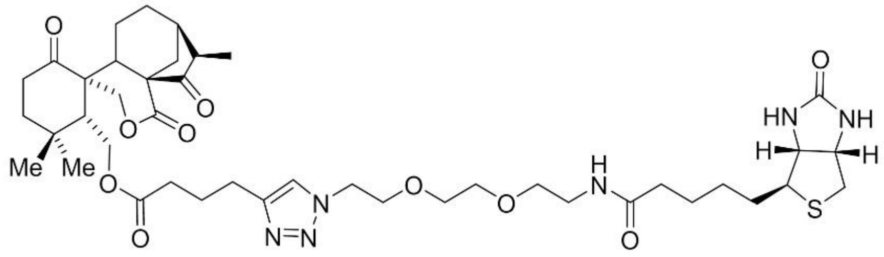

**Supplementary Figure S5. The structure of biotin-labeled Laxiflorin B analogues.**

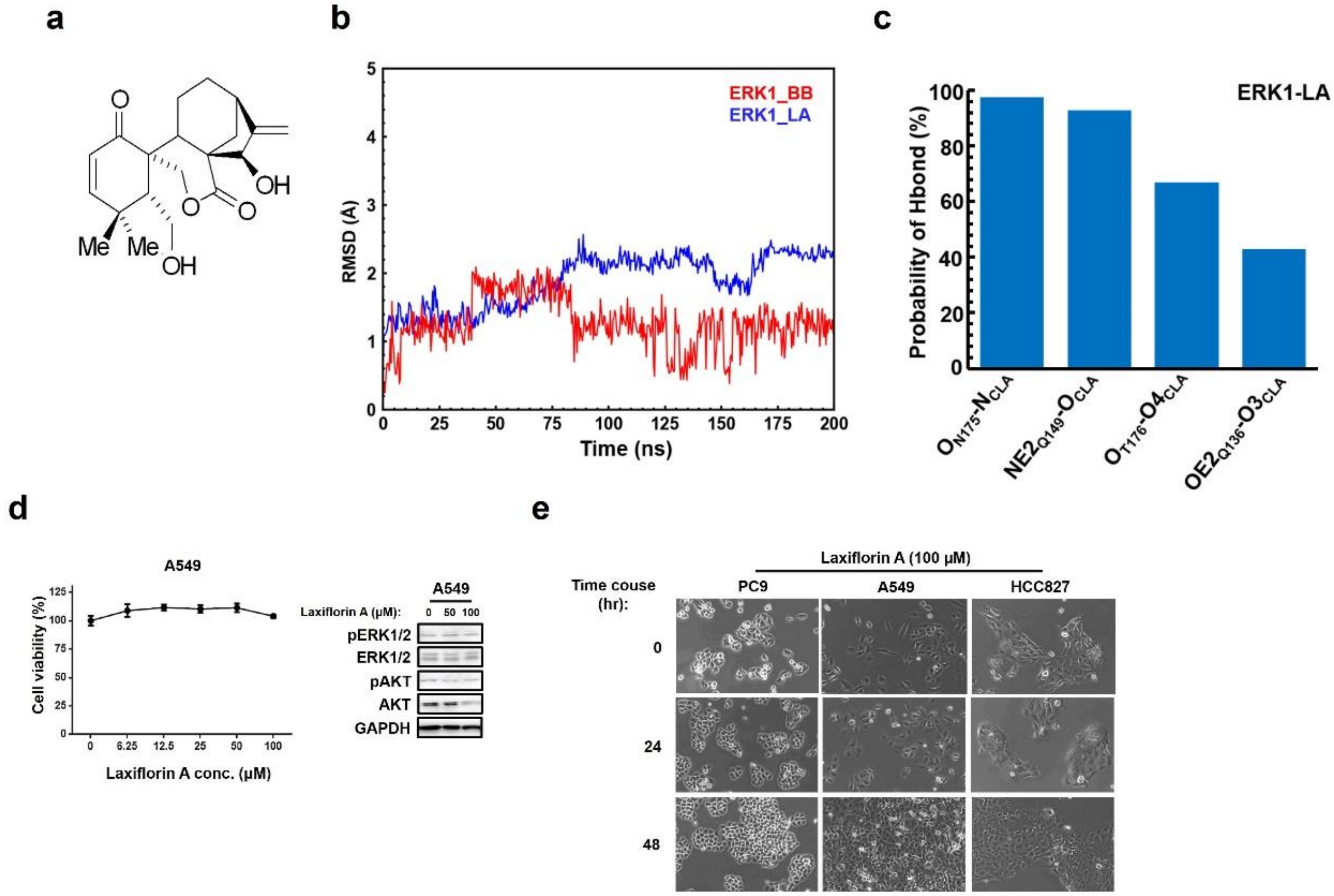


**Supplementary Figure S6. Influence of Laxiflorin A on NSCLC cell lines.**

**a** Molecular structure of Laxiflorin A. **b** Evolution of the RMSD values of backbone atoms in ERK1 and heavy atoms in Laxiflorin A with simulation time. **c** The probability of hydrogen bond formation between Laxiflorin A and residues of ERK1. **d** CCK-8 assay of cell viability and Western blot analysis of cell growth-related pathways after Laxiflorin A treatment of A549 cells for 48h. **e** Growth of A549, PC9 and HCC827 cells was monitored by microscopy after Laxiflorin A treatment for 24 and 48h.

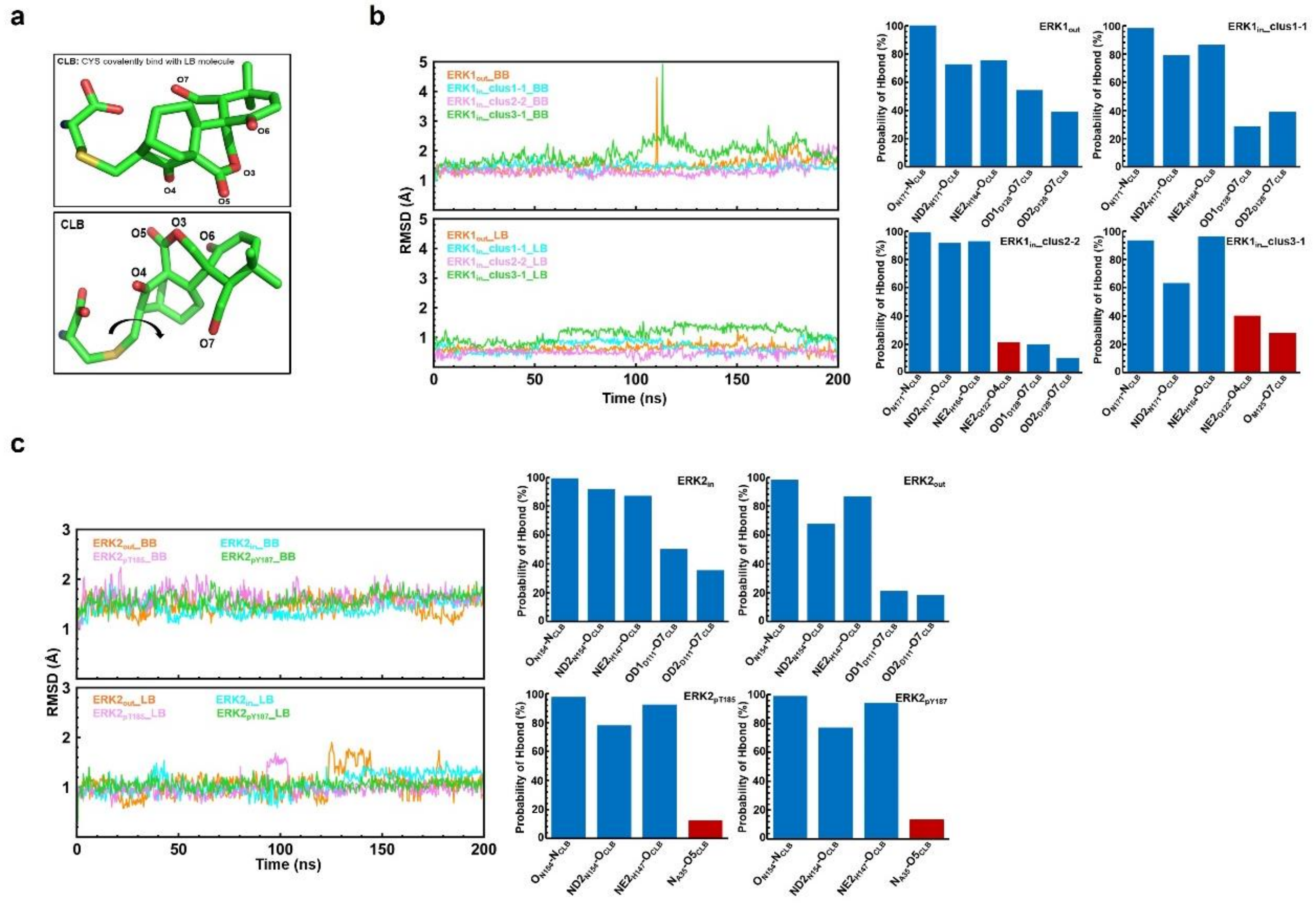


**Supplementary Figure S7. The root means square deviation (RMSD) and probability of hydrogen bond formation between Laxiflorin B and ERK1/2.**

**a** Schematic diagram of the “in” (upper panel) and “out” (lower panel) conformations of Laxiflorin B. **b** Evolution of the RMSD values of backbone atoms in ERK1 (upper left panel) and heavy atoms in Laxiflorin B (lower left panel) with simulation time. The probability of hydrogen bond formation between Laxiflorin B and residues in the binding pocket of ERK1 (right panels). Red bars show the non-conservative hydrogen bonds. **c** Evolution of the RMSD values of backbone atoms in ERK2 (upper left panel) and heavy atoms in Laxiflorin B (lower left panel) with simulation time. The probability of hydrogen bond formation between Laxiflorin B and residues in the binding pocket of ERK2 (right panels).

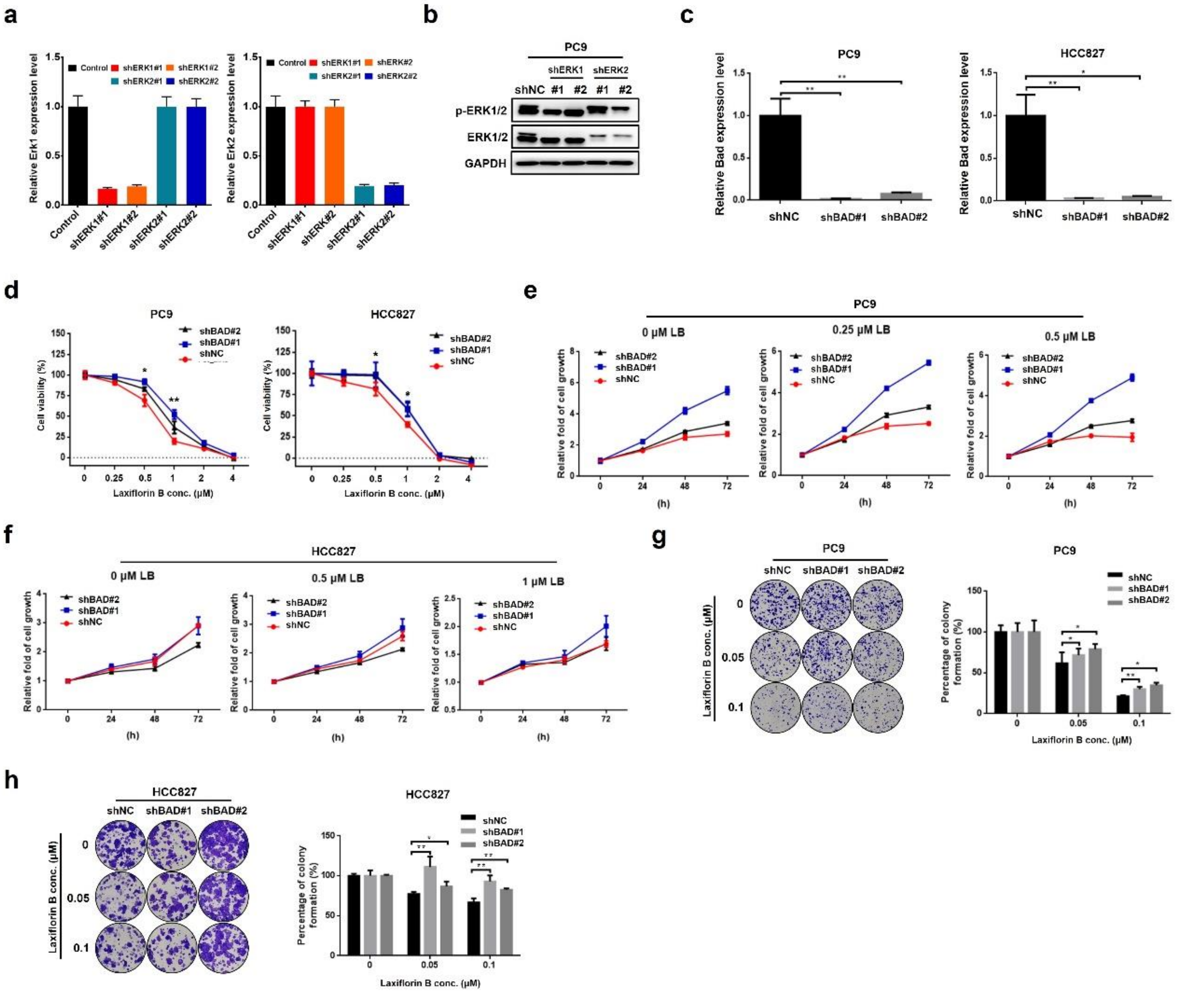


**Supplementary Figure S8. Laxiflorin B sensitivity of BAD-KD (knockdown) NSCLC cells**

**a-b** The knockdown specificity and efficiency of shERK1 and shERK2 were confirmed by qPCR and Western blotting in stably transfected PC9 cells. **c** qPCR analysis of BAD expression in PC9 and HCC827 stable cells stably transfected with shBAD#1 and shBAD#2. **d** CCK-8 assay of the sensitivity of stable BAD-KD PC9 and HCC827 cells to Laxiflorin B treatment for 48 h. **e-f** CCK-8 assay of stable BAD-KD PC9 and HCC827 cells after Laxiflorin B treatment for 48h. **g-h** 2-D Clonogenic assay of stable BAD-KD PC9 and HCC827 cells after Laxiflorin B treatment for 12 days. *$P < 0.05$; **$P < 0.01$.

a

b

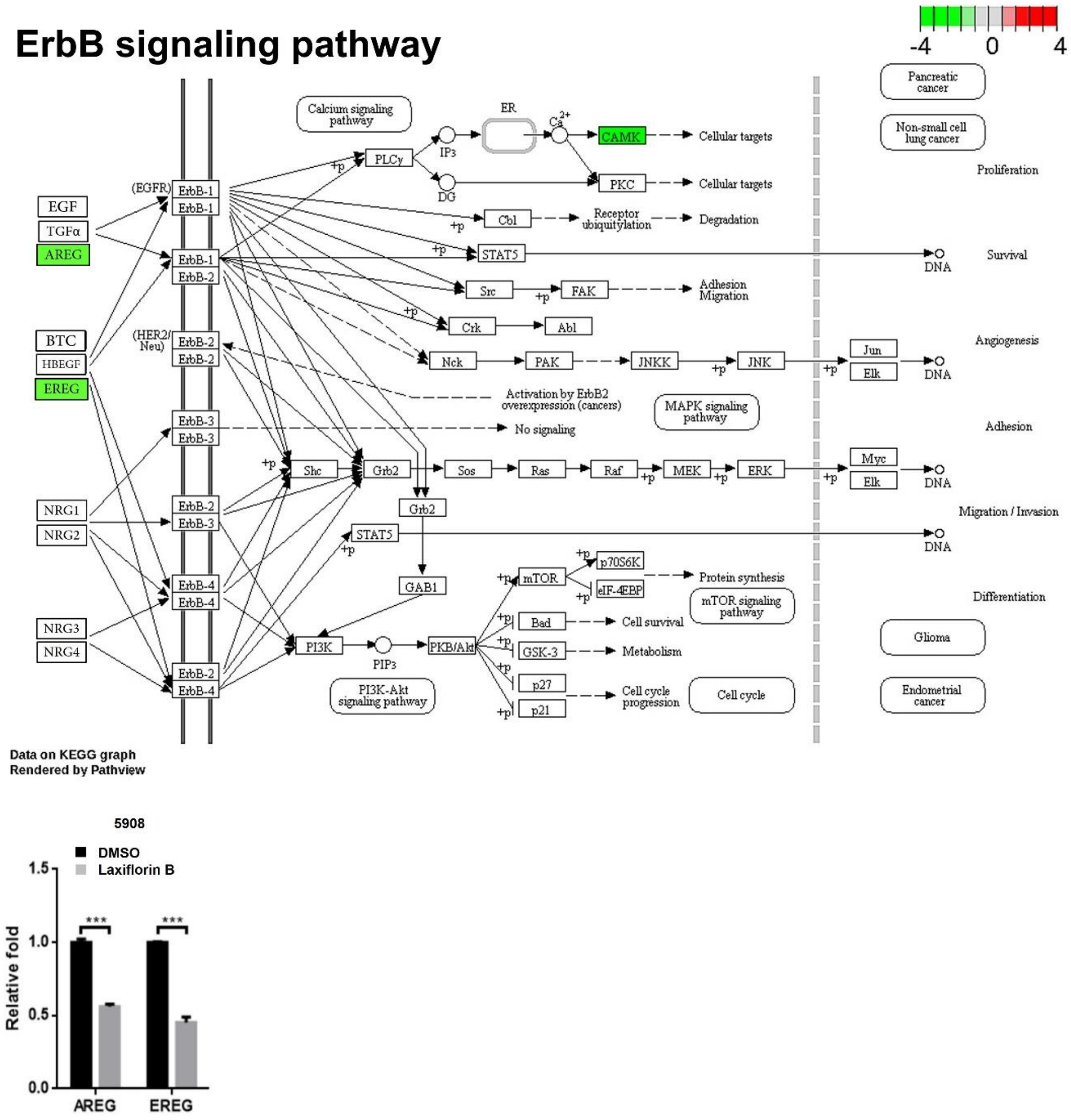


**Supplementary Figure S9. The influence of Laxiflorin B on the ErbB pathway.**

**a** KEGG analysis of gene expression in PC9 cells after Laxiflorin B treatment for 48 h. Downregulated genes in ErbB-associated pathways are shown in green. **b** The expression of *ARGE* and *ERGE* in 5908 cells after Laxiflorin B treatment for 48 h. *$P < 0.05$; ***$P < 0.001$.

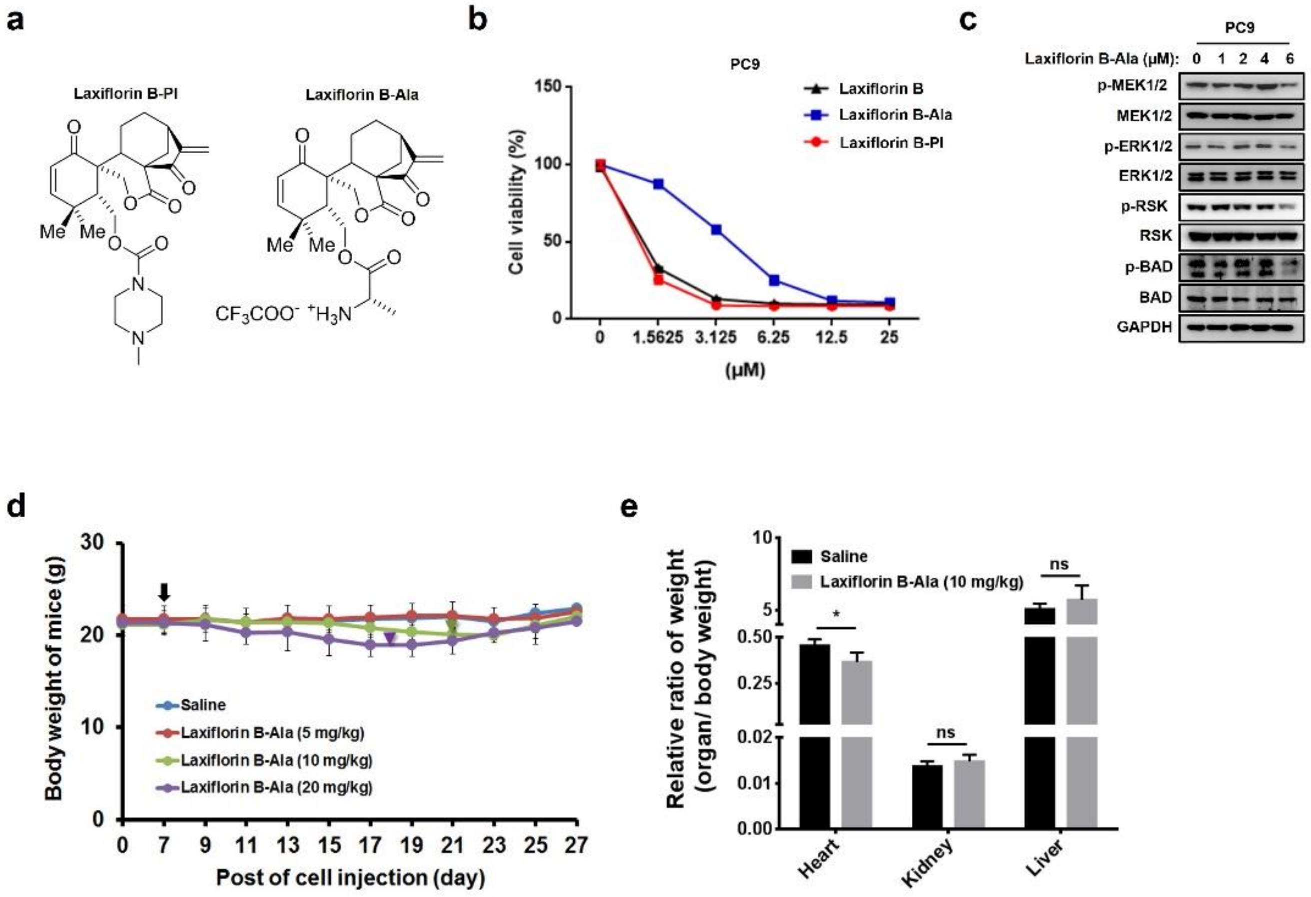


**Supplementary Figure S10.** ***In vivo*** **study and toxicity of Laxiflorin B.**

**a** Structures of Laxiflorin B-PI and Laxiflorin B-Ala: hydrophilic forms of Laxiflorin B. **b** CCK-8 assay of the effects of Laxiflorin B analogue on viability of PC9 cells for 48h. **c** Western bot analysis of the influence of Laxiflorin B-Ala on the MEK-ERK-RSK axis. **d** Body weight of nude mice monitored once every two days for 3 weeks after Laxiflorin B-Ala treatment. Black arrow indicates the initiation of Laxiflorin B treatment; green and purple inverted triangles indicate the termination of treatment in the 10 and 20 mg/kg groups on days 15 and 12, respectively. **e** Toxicity of Laxiflorin B-Ala monitored in the heart, kidney and liver of mice after Laxiflorin B treatment for 3 weeks. *$P < 0.05$; ns: not significant.

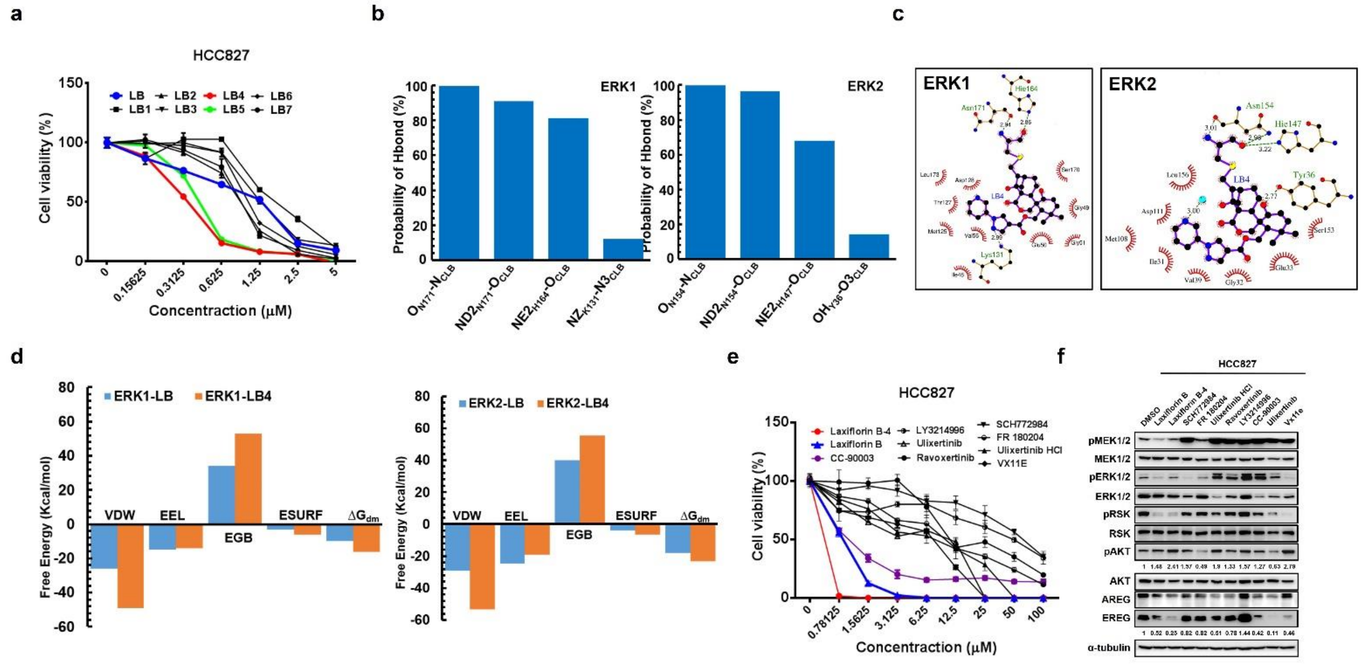


**Supplementary Figure S11. The modifiable capacity of Laxiflorin B.**

**a** $IC_{50}$ of Laxiflorin B analogues against HCC827 cells in CCK8 assays for 48h. **b** The probability of hydrogen bond formation between Laxiflorin B-4 and residues in the binding pocket of ERK1 and ERK2. **c** The detailed interactions between Laxiflorin B-4 and ERK1/2. The green dotted lines represent the hydrogen bonds. The hydrophobic interactions are shown as red arcs. **d** The noncovalent binding free energy of Laxiflorin B and Laxiflorin B-4. VDW and EEL represent the hydrophobic and electrostatic interactional energy in the gaseous phase. The sum of EGB and ESURF is the solvation free energy, where EGB is the electrostatic contribution and ESURF is the nonpolar contribution. **e** CCK-8 assay of the inhibitory effects of commercial ERK inhibitors, Laxiflorin B analogues on HCC827 cell viability for 48h. **f** Western blot analysis of the status of the MEK-ERK-RSK axis in HCC827 cells after treatment with commercial ERK inhibitors and Laxiflorin B analogues at 1 μM.

**Supplementary Table S1. Down-regulated genes by Laxiflorin B treatment in PC9**

| | logFC | logCPM | PValue | FDR | PC9-DMSO | | | PC9-LB (4μM) | | |
|---|---|---|---|---|---|---|---|---|---|---|
| **PDE7B** | -2.8976 | -0.9379 | 2.96E-06 | 2.11E-05 | 0.1795 | 0.0912 | 0.1407 | 0.0104 | 0.0104 | 0.0329 |
| **SCARNA2** | -2.649 | -1.5788 | 0.0014708 | 0.005725 | 0.767 | 0.4385 | 1.5266 | 0 | 0.1333 | 0.2811 |
| **BTBD19** | -2.6265 | -0.4757 | 3.99E-07 | 3.33E-06 | 0.9426 | 0.3514 | 0.7416 | 0.1073 | 0.1425 | 0.0751 |
| **FAM30A** | -2.5568 | 3.08258 | 1.20E-84 | 5.60E-82 | 1.5683 | 1.4687 | 1.4916 | 0.2281 | 0.297 | 0.264 |
| **MAGEA8** | -2.5387 | -1.1741 | 0.0002257 | 0.00109 | 0.2745 | 0.3778 | 0.1656 | 0 | 0.0265 | 0.1118 |
| **CAMK2A** | -2.4193 | 0.48499 | 3.10E-13 | 5.67E-12 | 0.5461 | 0.3426 | 0.482 | 0.0697 | 0.0926 | 0.0976 |
| **C11orf86** | -2.3884 | 0.57378 | 9.33E-12 | 1.47E-10 | 2.5575 | 1.2961 | 2.1169 | 0.4273 | 0.3783 | 0.349 |
| **KRT16P2** | -2.3201 | -1.3085 | 0.0008019 | 0.003353 | 0.4091 | 0.2338 | 0.2961 | 0.0357 | 0.0711 | 0.075 |
| **SLC34A2** | -2.1796 | -0.0215 | 8.27E-09 | 8.94E-08 | 0.3728 | 0.3108 | 0.3654 | 0.0949 | 0.0405 | 0.0996 |
| **TBX15** | -2.1651 | -0.5573 | 4.68E-06 | 3.22E-05 | 0.2212 | 0.2986 | 0.2835 | 0.0804 | 0 | 0.1013 |
| **LOC730338** | -2.1337 | -0.9381 | 0.0001168 | 0.000609 | 0.2647 | 0.4729 | 0.2994 | 0.0577 | 0.115 | 0.0606 |
| **AREG** | -2.1291 | -1.0811 | 0.0011945 | 0.004775 | 0.0864 | 0.2469 | 0.1407 | 0 | 0.0601 | 0.0475 |
| **SERPINB3** | -2.0195 | -0.0655 | 1.75E-06 | 1.30E-05 | 1.0945 | 0.4675 | 0.8537 | 0.1976 | 0.1968 | 0.2075 |
| **NTSR1** | -1.945 | -0.418 | 9.71E-05 | 0.000515 | 0.2026 | 0.1931 | 0.3385 | 0 | 0.1355 | 0.0571 |
| **EGR3** | -1.9442 | 3.90705 | 1.19E-57 | 2.31E-55 | 4.6243 | 5.2626 | 5.8882 | 1.5318 | 1.4883 | 1.19 |
| **SLC6A2** | -1.9117 | -0.8329 | 0.0002355 | 0.00113 | 0.2155 | 0.2211 | 0.12 | 0.0289 | 0.0288 | 0.0911 |
| **TNNI2** | -1.8857 | -0.9656 | 0.0006029 | 0.002605 | 0.9437 | 0.8991 | 0.6985 | 0.3743 | 0.2982 | 0 |
| **PHGR1** | -1.8696 | 0.72759 | 5.51E-12 | 8.96E-11 | 6.0723 | 5.7856 | 5.0918 | 1.5507 | 1.287 | 1.8998 |
| **SPRR3** | -1.8464 | -0.6659 | 7.53E-05 | 0.000409 | 0.9306 | 0.9549 | 0.8419 | 0.1874 | 0.1866 | 0.3935 |
| **SSPO** | -1.8456 | -0.7593 | 0.0010749 | 0.004338 | 0.0165 | 0.063 | 0.0636 | 0.0217 | 0.0072 | 0.0114 |
| **KRT13** | -1.8455 | 2.57707 | 7.74E-32 | 5.04E-30 | 5.6072 | 5.0517 | 5.7632 | 1.9304 | 1.5579 | 1.1882 |
| **CPZ** | -1.8439 | 2.23288 | 3.47E-30 | 2.11E-28 | 3.5445 | 3.3771 | 3.1529 | 1.0915 | 0.8283 | 0.9551 |
| **CKMT1B** | -1.7868 | -1.0222 | 0.0010297 | 0.004181 | 0.1826 | 0.2393 | 0.2065 | 0.0598 | 0.0595 | 0.0628 |
| **CYP21A2** | -1.7787 | -0.5363 | 0.000638 | 0.002742 | 0.7708 | 0.2754 | 0.3196 | 0.1681 | 0.0837 | 0.1471 |
| **FGFBP1** | -1.7471 | 2.02694 | 1.12E-23 | 4.59E-22 | 4.0535 | 4.8616 | 4.7458 | 1.4147 | 1.2432 | 1.5292 |
| **CYB5R2** | -1.7302 | 0.76965 | 1.38E-09 | 1.63E-08 | 1.5898 | 1.082 | 1.6781 | 0.5285 | 0.3947 | 0.4162 |
| **AOC1** | -1.7194 | 5.68947 | 6.72E-181 | 2.51E-177 | 30.598 | 31.492 | 32.663 | 10.485 | 9.9935 | 9.0922 |
| **LINC02015** | -1.6942 | -0.8586 | 0.0010038 | 0.004086 | 0.2397 | 0.2491 | 0.1971 | 0.1141 | 0.0379 | 0.0599 |

| | logFC | logCPM | PValue | FDR | PC9-DMSO | | | PC9-LB (4μM) | | |
|---|---|---|---|---|---|---|---|---|---|---|
| **MILR1** | -1.6937 | 2.98726 | 7.06E-34 | 5.17E-32 | 19.91 | 17.705 | 19.583 | 7.3943 | 5.8559 | 4.8644 |
| **H19** | -1.6869 | 7.00378 | 6.38E-161 | 1.59E-157 | 90.23 | 79.967 | 78.472 | 27.531 | 27.228 | 24.624 |
| **TMOD4** | -1.6686 | -0.5997 | 0.0002849 | 0.001341 | 0.889 | 0.5481 | 0.615 | 0.2738 | 0.1818 | 0.1916 |
| **NEURL3** | -1.6178 | 5.04477 | 5.12E-94 | 2.94E-91 | 32.876 | 37.26 | 31.942 | 11.798 | 11.443 | 10.931 |
| **C1orf167** | -1.6158 | -0.8079 | 0.0022511 | 0.008241 | 0.2068 | 0.092 | 0.1497 | 0.0722 | 0 | 0.0758 |
| **GABRP** | -1.6136 | -0.1423 | 5.27E-05 | 0.000297 | 0.2926 | 0.3531 | 0.4412 | 0.1872 | 0.1017 | 0.0715 |
| **PTGS2** | -1.6132 | 1.85619 | 2.17E-18 | 6.23E-17 | 1.1285 | 1.3338 | 1.0856 | 0.4113 | 0.4221 | 0.3534 |
| **ITPRID1** | -1.5919 | 2.33999 | 3.94E-23 | 1.53E-21 | 0.9809 | 1.0604 | 1.2373 | 0.3785 | 0.3606 | 0.3802 |
| **MUC5B-AS1** | -1.5884 | 0.73596 | 2.32E-09 | 2.68E-08 | 6.0863 | 5.0917 | 5.1035 | 1.9429 | 1.548 | 2.0401 |
| **RARRES1** | -1.5652 | 5.80437 | 4.86E-157 | 1.04E-153 | 48.103 | 49.881 | 47.433 | 16.474 | 18.335 | 15.679 |
| **LOC101928505** | -1.5587 | -0.666 | 0.0009949 | 0.004053 | 0.5815 | 0.2955 | 0.5261 | 0.1353 | 0.2021 | 0.1421 |
| **SYT8** | -1.5513 | 1.21197 | 2.06E-09 | 2.39E-08 | 2.2787 | 2.2501 | 1.8742 | 1.0122 | 0.8641 | 0.3416 |
| **EDAR** | -1.5488 | -0.759 | 0.0019067 | 0.007157 | 0.2379 | 0.1511 | 0.1435 | 0.1107 | 0.0413 | 0.0291 |
| **CAMK2B** | -1.5215 | 1.73246 | 5.36E-11 | 7.77E-10 | 1.9113 | 1.6187 | 2.3776 | 0.4864 | 0.9458 | 0.681 |
| **LOC105369332** | -1.5199 | -0.326 | 0.0003933 | 0.001783 | 1.6021 | 2.3987 | 1.4494 | 0.8986 | 0.2983 | 0.7339 |
| **RGS7** | -1.5176 | 1.83182 | 6.40E-12 | 1.03E-10 | 1.7559 | 2.3619 | 2.0558 | 0.8787 | 0.8976 | 0.4259 |
| **PLAT** | -1.5152 | 4.43613 | 6.78E-65 | 1.69E-62 | 10.741 | 9.6167 | 10.72 | 3.9567 | 3.3595 | 3.8675 |
| **TRIM31** | -1.5022 | 4.96819 | 7.42E-83 | 3.26E-80 | 26.132 | 23.682 | 26.209 | 9.9311 | 8.4734 | 9.1937 |
| **MUC5AC** | -1.4906 | 0.22918 | 6.51E-06 | 4.35E-05 | 0.2717 | 0.2071 | 0.3146 | 0.0948 | 0.1133 | 0.0797 |
| **FGD2** | -1.4846 | -0.5572 | 0.0020303 | 0.007551 | 0.4243 | 0.2223 | 0.1919 | 0.1111 | 0.0553 | 0.1361 |
| **KRT9** | -1.4844 | -0.0076 | 2.74E-05 | 0.000162 | 0.5614 | 0.4814 | 0.6603 | 0.2939 | 0.1708 | 0.1543 |
| **HPGD** | -1.4587 | 7.38025 | 2.04E-170 | 6.10E-167 | 82.373 | 89.771 | 90.825 | 34.54 | 30.562 | 33.314 |
| **SMAD9** | -1.457 | 1.99132 | 2.62E-18 | 7.46E-17 | 1.0697 | 1.0303 | 1.0209 | 0.3385 | 0.3984 | 0.4308 |
| **MICALCL** | -1.45 | 1.12312 | 1.99E-10 | 2.67E-09 | 0.5122 | 0.4981 | 0.502 | 0.2328 | 0.1669 | 0.1662 |
| **TNC** | -1.4359 | 3.65752 | 2.38E-35 | 1.86E-33 | 2.2211 | 1.914 | 2.3249 | 0.8665 | 0.8564 | 0.7293 |
| **ANK1** | -1.429 | 1.10995 | 2.38E-09 | 2.74E-08 | 0.8326 | 1.1503 | 0.9228 | 0.3269 | 0.3437 | 0.4387 |
| **SMG7-AS1** | -1.4221 | -0.5163 | 0.002439 | 0.008819 | 0.3149 | 0.2 | 0.5223 | 0.1374 | 0.114 | 0.1443 |
| **CHD5** | -1.4065 | 3.55184 | 1.28E-38 | 1.16E-36 | 1.7856 | 1.6951 | 1.6747 | 0.7647 | 0.6195 | 0.6113 |
| **RIPOR2** | -1.4025 | 0.89702 | 2.25E-07 | 1.95E-06 | 0.4924 | 0.389 | 0.5432 | 0.1886 | 0.24 | 0.121 |
| **BDNF-AS** | -1.4018 | -0.3991 | 0.000741 | 0.00312 | 0.4443 | 0.4233 | 0.3517 | 0.1212 | 0.1931 | 0.1527 |

| | logFC | logCPM | PValue | FDR | PC9-DMSO | | | PC9-LB (4μM) | | |
|---|---|---|---|---|---|---|---|---|---|---|
| **GPR78** | -1.3722 | 3.69731 | 1.65E-36 | 1.37E-34 | 3.9015 | 4.4381 | 4.4826 | 1.4756 | 1.8821 | 1.7397 |
| **LOC101927954** | -1.3667 | -0.1899 | 0.0014049 | 0.005507 | 0.3712 | 0.3705 | 0.1759 | 0.1851 | 0.0768 | 0.0972 |
| **RAB37** | -1.3601 | 2.57591 | 6.16E-17 | 1.57E-15 | 2.539 | 3.2456 | 2.5459 | 1.2923 | 1.1216 | 0.9111 |
| **CALML5** | -1.3595 | 1.58411 | 8.77E-08 | 8.10E-07 | 5.676 | 5.127 | 3.6013 | 2.7014 | 1.3452 | 1.6884 |
| **NTNG2** | -1.3558 | 0.00691 | 0.0007285 | 0.003072 | 0.3034 | 0.3548 | 0.1747 | 0.0602 | 0.1079 | 0.1643 |
| **PECAM1** | -1.3513 | 3.16424 | 1.32E-25 | 6.12E-24 | 2.8791 | 2.5915 | 3.1153 | 1.073 | 1.0561 | 1.3388 |
| **FAM131B** | -1.3496 | -0.1423 | 0.0004222 | 0.001898 | 0.2355 | 0.2524 | 0.293 | 0.0771 | 0.0895 | 0.1483 |
| **WFDC10B** | -1.3407 | 0.36802 | 1.26E-05 | 7.94E-05 | 4.187 | 3.3459 | 3.1771 | 1.7678 | 1.1737 | 1.3612 |
| **HOTS** | -1.3272 | 4.70128 | 2.00E-68 | 5.65E-66 | 9.7128 | 9.1594 | 9.6721 | 3.5722 | 4.1482 | 3.9787 |
| **OGDHL** | -1.3235 | 0.03477 | 0.0007782 | 0.003264 | 0.4477 | 0.2218 | 0.4212 | 0.2031 | 0.0622 | 0.1805 |
| **EXOC3L4** | -1.3218 | 2.15028 | 6.35E-16 | 1.51E-14 | 2.3982 | 2.1659 | 2.6443 | 0.9809 | 0.9335 | 1.0528 |
| **CALHM3** | -1.3127 | 2.10122 | 9.09E-15 | 1.97E-13 | 3.3929 | 3.79 | 3.6694 | 1.8715 | 1.2539 | 1.3578 |
| **LOC91370** | -1.3097 | 0.15447 | 9.75E-05 | 0.000517 | 1.8901 | 1.9574 | 1.6356 | 0.8604 | 0.9997 | 0.3765 |
| **EREG** | -1.3095 | 3.96572 | 1.79E-32 | 1.21E-30 | 4.3295 | 5.1763 | 4.8647 | 1.6451 | 2.1118 | 2.2137 |
| **PTGIS** | -1.3085 | 1.71907 | 3.60E-10 | 4.62E-09 | 0.6362 | 0.8927 | 0.9104 | 0.4037 | 0.3216 | 0.2861 |
| **LINC00887** | -1.3039 | 0.02067 | 0.0003398 | 0.001566 | 0.4277 | 0.4482 | 0.4063 | 0.1493 | 0.2973 | 0.0784 |
| **C1orf116** | -1.2989 | 4.53844 | 3.03E-41 | 3.17E-39 | 5.1297 | 6.4051 | 6.3469 | 2.4832 | 2.3713 | 2.6289 |
| **HBEGF** | -1.292 | 3.09153 | 8.30E-22 | 2.99E-20 | 4.5355 | 5.0242 | 5.6359 | 1.788 | 2.3268 | 2.278 |
| **EFNB2** | -1.2709 | 3.90817 | 1.99E-38 | 1.78E-36 | 4.7728 | 5.0232 | 4.6237 | 1.8324 | 1.9781 | 2.3412 |
| **S100A8** | -1.2706 | 0.8895 | 3.94E-05 | 0.000228 | 4.653 | 4.0923 | 4.9652 | 0.8328 | 1.9699 | 3.0607 |
| **AATK** | -1.2682 | 0.49429 | 1.23E-05 | 7.73E-05 | 0.3292 | 0.3377 | 0.4123 | 0.1436 | 0.143 | 0.174 |
| **IL1RL1** | -1.2653 | 4.58549 | 6.74E-60 | 1.44E-57 | 8.4855 | 7.843 | 8.5523 | 3.4875 | 3.68 | 3.4731 |
| **SPRR1A** | -1.2605 | -0.0075 | 0.0008334 | 0.00347 | 2.5127 | 1.5322 | 1.8186 | 1.2278 | 0.8734 | 0.3683 |
| **HOXA-AS2** | -1.2585 | 2.3028 | 2.08E-13 | 3.88E-12 | 6.209 | 7.7315 | 5.6174 | 2.7893 | 2.5108 | 3.0978 |
| **CRLF2** | -1.2521 | 0.51441 | 1.17E-05 | 7.40E-05 | 2.6364 | 2.0409 | 2.4597 | 1.1502 | 1.1455 | 0.7548 |
| **ZBTB7C** | -1.2476 | 3.18445 | 2.64E-23 | 1.05E-21 | 2.6642 | 2.2758 | 2.7903 | 1.0421 | 1.1063 | 1.2024 |
| **MAMDC4** | -1.2458 | 1.5739 | 3.33E-10 | 4.32E-09 | 1.1201 | 1.0339 | 1.1401 | 0.6109 | 0.3954 | 0.4169 |
| **LOC107001062** | -1.2449 | 0.81875 | 1.21E-05 | 7.65E-05 | 4.0871 | 4.7139 | 2.9192 | 2.1585 | 1.2151 | 1.6753 |
| **ARSI** | -1.2444 | 4.76664 | 1.72E-48 | 2.39E-46 | 11.672 | 13.586 | 11.403 | 5.0021 | 5.5392 | 5.3662 |
| **ATG9B** | -1.2331 | 2.77695 | 4.88E-20 | 1.58E-18 | 2.2667 | 1.9987 | 1.9616 | 0.9951 | 0.832 | 0.8901 |

| | logFC | logCPM | PValue | FDR | PC9-DMSO | | | PC9-LB (4μM) | | |
|---|---|---|---|---|---|---|---|---|---|---|
| **VWF** | -1.2224 | -0.3261 | 0.0024559 | 0.008863 | 0.1021 | 0.0972 | 0.1253 | 0.0445 | 0.038 | 0.0601 |
| **CALML3-AS1** | -1.2217 | 0.67434 | 1.75E-05 | 0.000107 | 0.4478 | 0.6997 | 0.5834 | 0.2969 | 0.1868 | 0.279 |
| **DKK1** | -1.2118 | 7.79464 | 3.15E-119 | 3.14E-116 | 167.9 | 169.22 | 172.21 | 80.351 | 76.86 | 68.839 |
| **MGAM** | -1.2062 | 0.04847 | 0.0027629 | 0.009829 | 0.2583 | 0.1041 | 0.2337 | 0.13 | 0.0691 | 0.0637 |
| **ATP6V1FNB** | -1.2061 | 2.80805 | 6.68E-15 | 1.47E-13 | 2.8538 | 2.4819 | 2.1316 | 1.3029 | 1.0237 | 0.9881 |
| **CXCL17** | -1.1922 | 3.88224 | 7.42E-25 | 3.27E-23 | 19.051 | 17.465 | 20.39 | 9.2817 | 6.8938 | 9.4709 |
| **HLA-F** | -1.1892 | 0.076 | 0.001939 | 0.007268 | 1.8535 | 1.4006 | 0.6939 | 0.5019 | 0.6665 | 0.5856 |
| **MT1A** | -1.1857 | -0.1423 | 0.0009989 | 0.004068 | 2.7658 | 2.9452 | 2.6494 | 1.2776 | 1.131 | 1.3415 |
| **LOC692247** | -1.1844 | -0.0511 | 0.001693 | 0.006469 | 1.9232 | 3.0921 | 1.6312 | 1.3634 | 0.8356 | 0.7709 |
| **CREB3L1** | -1.1843 | 3.98785 | 4.19E-41 | 4.32E-39 | 7.9502 | 8.106 | 8.5962 | 3.6377 | 3.6229 | 3.9085 |
| **RN7SL2** | -1.182 | 8.31884 | 3.06E-09 | 3.48E-08 | 2049.6 | 1075.8 | 1264.2 | 791.33 | 714.71 | 484.07 |
| **PLA2G2F** | -1.1794 | 3.49582 | 4.65E-32 | 3.08E-30 | 5.8052 | 5.5988 | 5.7023 | 2.543 | 2.6974 | 2.5183 |
| **UNC5B** | -1.1783 | 6.01008 | 1.60E-110 | 1.33E-107 | 13.104 | 12.629 | 13.134 | 5.688 | 6.2298 | 5.7397 |
| **FSTL4** | -1.178 | 3.29166 | 5.06E-26 | 2.41E-24 | 2.5445 | 2.322 | 2.6047 | 1.1779 | 1.1419 | 1.0726 |
| **SYT12** | -1.1696 | 5.94728 | 3.53E-69 | 1.04E-66 | 24.944 | 21.955 | 21.316 | 10.106 | 10.831 | 10.231 |
| **CAPN8** | -1.1645 | 6.67102 | 1.51E-134 | 2.26E-131 | 69.148 | 65.533 | 64.557 | 30.047 | 31.044 | 30.287 |
| **PADI1** | -1.1592 | 2.8154 | 1.42E-17 | 3.83E-16 | 2.5462 | 2.5696 | 2.3642 | 1.1255 | 1.3538 | 0.9516 |
| **AREG** | -1.1565 | 2.26965 | 1.39E-08 | 1.45E-07 | 4.6504 | 5.8247 | 5.3418 | 2.7355 | 3.0422 | 1.4841 |
| **TMEM178A** | -1.1514 | 0.94974 | 6.16E-06 | 4.14E-05 | 1.8197 | 1.328 | 1.4011 | 0.6757 | 0.572 | 0.8514 |
| **PDZD7** | -1.1492 | 1.17413 | 1.58E-07 | 1.40E-06 | 1.4101 | 1.4928 | 1.4175 | 0.6835 | 0.708 | 0.6029 |
| **LOC101927751** | -1.1465 | 5.31932 | 6.94E-67 | 1.82E-64 | 13.133 | 12.82 | 12.991 | 5.3754 | 6.4589 | 6.2622 |
| **TNFSF10** | -1.1443 | 4.69155 | 4.66E-34 | 3.44E-32 | 15.82 | 19.186 | 20.355 | 8.5352 | 8.3539 | 8.9006 |
| **MUC6** | -1.1318 | -0.174 | 0.0024543 | 0.00886 | 0.1286 | 0.1608 | 0.1091 | 0.0701 | 0.0419 | 0.0736 |
| **LMF1** | -1.1302 | 2.16959 | 5.37E-11 | 7.77E-10 | 2.579 | 2.3424 | 1.8753 | 1.1356 | 1.0472 | 0.9937 |
| **CASP14** | -1.1297 | 1.21768 | 4.71E-07 | 3.88E-06 | 1.0656 | 0.8005 | 1.0382 | 0.4291 | 0.4452 | 0.4881 |
| **CRYBG2** | -1.1265 | 5.31435 | 1.04E-73 | 3.32E-71 | 10.637 | 10.229 | 10.113 | 5.1016 | 4.5899 | 4.8956 |
| **SPACA6** | -1.1245 | 3.15833 | 2.47E-14 | 5.07E-13 | 6.4951 | 5.3133 | 6.6479 | 2.6619 | 3.5633 | 2.4645 |
| **PRR22** | -1.124 | 0.46397 | 0.0004416 | 0.001976 | 1.6892 | 1.1624 | 0.9764 | 0.696 | 0.4485 | 0.6449 |
| **EGFL7** | -1.119 | 5.2281 | 4.85E-76 | 1.65E-73 | 38.596 | 39.084 | 39.619 | 19.343 | 18.318 | 17.863 |
| **CCDC71L** | -1.1146 | 5.78955 | 8.94E-77 | 3.26E-74 | 10.315 | 11.518 | 11.116 | 5.064 | 5.5519 | 5.0321 |

| | logFC | logCPM | PValue | FDR | PC9-DMSO | | | PC9-LB (4μM) | | |
|---|---|---|---|---|---|---|---|---|---|---|
| **UNC5B-AS1** | -1.1049 | 1.34073 | 9.33E-08 | 8.56E-07 | 5.178 | 5.2181 | 4.9549 | 2.8672 | 2.1633 | 2.2808 |
| **ITGB2** | -1.1008 | 3.88079 | 4.40E-28 | 2.35E-26 | 6.8807 | 6.8298 | 6.8056 | 3.7837 | 2.9031 | 3.1419 |
| **GASK1A** | -1.0977 | 2.43988 | 1.00E-14 | 2.16E-13 | 2.1862 | 1.9924 | 2.2359 | 1.0118 | 1.0738 | 0.9928 |
| **SLC23A3** | -1.0963 | 0.25295 | 0.001426 | 0.005575 | 0.5676 | 0.9014 | 0.6847 | 0.4127 | 0.4385 | 0.1734 |
| **MYO15B** | -1.0892 | 4.51679 | 6.25E-25 | 2.77E-23 | 3.0798 | 3.5457 | 3.0919 | 1.821 | 1.303 | 1.5718 |
| **S100A4** | -1.0891 | 4.72126 | 1.01E-54 | 1.80E-52 | 64.655 | 61.71 | 63.042 | 30.3 | 31.269 | 29.933 |
| **TNFRSF19** | -1.0876 | 2.0875 | 3.58E-10 | 4.60E-09 | 1.25 | 1.2483 | 1.4442 | 0.6439 | 0.4973 | 0.7727 |
| **FOXJ1** | -1.0869 | 0.41133 | 0.0002623 | 0.001247 | 0.7723 | 0.6646 | 0.4959 | 0.3478 | 0.3031 | 0.2739 |
| **ALDH1A3** | -1.069 | 6.99514 | 1.88E-144 | 3.12E-141 | 51.858 | 51.745 | 52.381 | 25.969 | 25.659 | 24.826 |
| **ATP6V1B1** | -1.0656 | 4.1404 | 1.89E-33 | 1.36E-31 | 12.196 | 11.878 | 13.112 | 6.3967 | 5.6368 | 6.2526 |
| **RN7SL1** | -1.0616 | 9.18893 | 6.82E-10 | 8.37E-09 | 3458.1 | 1885 | 2462.5 | 1460.6 | 1380.9 | 1006 |
| **CRAT** | -1.061 | 6.59694 | 4.71E-103 | 3.52E-100 | 43.609 | 43.986 | 47.612 | 22.49 | 21.852 | 22.341 |
| **GJA5** | -1.0519 | 2.65854 | 1.08E-14 | 2.30E-13 | 2.4119 | 2.6485 | 2.8848 | 1.3911 | 1.2256 | 1.3296 |
| **MIR205HG** | -1.0518 | 6.09213 | 1.55E-101 | 1.06E-98 | 96.081 | 100.71 | 99.101 | 50.617 | 47.564 | 48.615 |
| **CTDSP1** | -1.0517 | 6.26004 | 2.91E-94 | 1.74E-91 | 39.051 | 37.556 | 39.794 | 19.583 | 19.865 | 18.281 |
| **YBX2** | -1.05 | 0.42178 | 0.0002294 | 0.001106 | 0.9348 | 1.1134 | 0.9515 | 0.4078 | 0.5416 | 0.5353 |
| **EME2** | -1.0492 | 2.31371 | 1.82E-11 | 2.79E-10 | 5.6514 | 5.7615 | 5.8288 | 2.4162 | 2.652 | 3.521 |
| **SERPINB5** | -1.0391 | 6.6566 | 2.75E-95 | 1.79E-92 | 49.827 | 53.646 | 52.269 | 25.738 | 24.551 | 27.711 |
| **MAPK15** | -1.0371 | 4.18546 | 1.10E-34 | 8.37E-33 | 12.982 | 13.648 | 12.274 | 6.5498 | 6.6429 | 6.2782 |
| **MMP28** | -1.0365 | 1.88621 | 7.79E-08 | 7.25E-07 | 1.9839 | 1.583 | 1.9968 | 1.0602 | 1.0344 | 0.6816 |
| **PPT2-EGFL8** | -1.0333 | 1.57857 | 6.85E-07 | 5.51E-06 | 1.6059 | 1.3726 | 1.282 | 0.577 | 0.9235 | 0.6275 |
| **MICAL2** | -1.0287 | 4.31613 | 1.85E-18 | 5.38E-17 | 6.7724 | 5.934 | 7.0397 | 2.6124 | 3.7565 | 3.5939 |
| **BPIFB1** | -1.0278 | 3.94452 | 3.69E-23 | 1.44E-21 | 11.268 | 12.83 | 13.249 | 6.4402 | 5.4928 | 6.9423 |
| **TCIM** | -1.0251 | 5.60625 | 1.33E-64 | 3.27E-62 | 33.111 | 36.146 | 36.84 | 17.119 | 18.427 | 18.105 |
| **ERN2** | -1.0222 | 0.46394 | 0.0019018 | 0.007142 | 0.5952 | 0.5139 | 0.3702 | 0.1947 | 0.1778 | 0.3749 |
| **ZBED2** | -1.0217 | 1.18683 | 0.0005395 | 0.00236 | 1.164 | 1.8299 | 0.9478 | 0.6602 | 0.4552 | 0.8798 |
| **EGR4** | -1.0165 | 0.27663 | 0.0022329 | 0.008189 | 0.5804 | 0.553 | 0.8402 | 0.2279 | 0.3278 | 0.452 |
| **AK5** | -1.0162 | 0.54395 | 0.0005754 | 0.002499 | 0.5729 | 0.4211 | 0.385 | 0.2428 | 0.2845 | 0.165 |
| **WNT10A** | -1.0075 | 2.74634 | 6.03E-12 | 9.73E-11 | 4.4854 | 4.2437 | 4.1431 | 2.1621 | 2.6712 | 1.7243 |

**Supplementary Table S2. $IC_{50}$ of Laxiflorin B in PC9 and HCC827 BAD-knockdown stable cells**

| | | PC9 | | | HCC827 | | |
|---|---|---|---|---|---|---|---|
| | | shNC | shBad#1 | shBad#2 | shNC | shBad#1 | shBad#2 |
| $IC_{50}$ (μM) | 24h | 3.175 | 1.997 | 3.048 | 1.875 | 2.081 | 2.918 |
| | 48h | 1.028 | 1.156 | 1.388 | 1.217 | 1.26 | 1.471 |
| | 72h | 0.649 | 1.076 | 0.8523 | 0.8335 | 1.062 | 1.059 |

**Supplementary Table S3. IC50 of Laxiflorin B analogous in non-small cellular lung cancer cell lines for 48h**

| Status | | PC9 |
|---|---|---|
| EGFR | | Ex19del |
| KRAS | | WT |
| $IC_{50}$ (μM) | Laxiflorin B | 1.734 |
| | Laxiflorin B-PI | 2.771 |
| | Laxiflorin B-Ala | 3.848 |

**Supplementary Table S4. $IC_{50}$ of Laxiflorin B analogous in non-small cellular lung cancer cell lines for 48h**

| | Compound | PC9 | HCC827 |
|---|---|---|---|
| | LB | 2.125 | 0.974 |
| | LB1 | 4.021 | 1.801 |
| | LB2 | 1.518 | 0.883 |
| | LB3 | 1.05 | 1.358 |
| | LB4 | 0.605 | 0.337 |
| | LB5 | 0.785 | 0.414 |
| $IC_{50}$ | LB6 | 2.101 | 1.067 |
| (μM) | LB7 | 1.092 | 0.899 |

**Supplementary Table S5. $IC_{50}$ of Laxiflorin B analogous and ERK inhibitors in non-small cellular lung cancer cell lines for 48h**

| | PC9 | HCC827 | $K_i$ of ERK1 and 2 | $IC_{50}$of ERK1 and 2 | Clinical trial |
|---|---|---|---|---|---|
| Laxiflorin B | 5.034 | 0.844 | | | |
| Laxiflorin B-4 | 2.118 | <0.5 | | | |
| CC-90003 | 4.776 | 0.639 | | 10-20 nM | Phase I |
| VX11E | 12.23 | 4.443 | < 2 nM | | |
| Ulixertinib | 18.9 | 4.575 | 0.3 and 0.04 nM | <0.3 nM | Phase I/II |
| Ulixertinib HCl | 27.72 | 5.85 | 0.3 and 0.04 nM | <0.3 nM | |
| FR 180204 | 54.81 | 9.962 | 0.31 and 0.14 μM | 0.51 and 0.33 μM | |
| Ravoxertinib | 75.74 | 21.25 | | 6. 1and 3.1 nM | Phase I |
| LY3214996 | 122.4 | 49.4 | | 5 nM | Phase I |
| SCH772984 | 167.5 | 60.4 | | 4 and 1 nM | |
